# The circadian clock regulates scavenging of fluid-borne substrates by brain border-associated macrophages

**DOI:** 10.64898/2025.12.08.693074

**Authors:** Helena J. Barr, Sydney K. Caldwell, Justin M. Reimertz, Constanze Depp, Alec J. Walker, Samuel E. Marsh, Anushree S. Gupte, Maximilian Hingerl, Jasmin Patel, Sang-O Park, Sarah Ferraro, Jonathan Lipton, Beth Stevens

## Abstract

Circadian disruptions perturb the brain and immune system and increase the risk of developing Alzheimer’s Disease (AD), yet whether this involves dysregulation of brain immunity remains less clear. Here, we perform single-cell RNA sequencing of the brain immune compartment around the day-night cycle and identify brain border-associated macrophages (BAMs) as highly rhythmic cells. During the rest phase, we find that BAMs exhibit coordinated upregulation of endocytic genes and enhanced uptake of extracellular fluid-borne material including amyloid-beta (Aβ). Rhythmicity in BAM scavenging is regulated by the clock gene *Bmal1*, mediated by the endocytic receptor CD206, and perturbed with age. In a mouse model of AD, we show that deletion of *Bmal1* in BAMs worsens perivascular and leptomeningeal Aβ plaque burden. Our results identify endocytosis as a specialized and rhythmic BAM function and identify perturbed timing of brain border immune functions as a potential mechanism by which circadian disruptions precipitate amyloidosis.

## INTRODUCTION

Circadian rhythmicity is a ubiquitous organizing principle that allows cells and organisms to anticipate, rather than simply respond to, predictable changes in light and temperature^1–3^. In mammals, circadian rhythms are genetically encoded by a 24-hour transcription-translation feedback loop driven by the clock genes *Bmal1* and *Clock* and their repressors, including the period and cryptochrome proteins (PER1, PER2, CRY1, CRY2) and the REV-ERB nuclear receptors (REV-ERBα, REV-ERBβ), that together synchronize physiology to time-of-day^2^. Accumulating epidemiological evidence supports a causative link between circadian disruptions and neurodegenerative disease^4^. Night-shift work, sleep fragmentation, and lack of rhythmicity in daily activities are all associated with an increased risk of developing dementia or Alzheimer’s Disease (AD)^5–13^, and sleep perturbations or clock gene deletion worsen amyloid-beta (Aβ) plaque deposition in mouse models of AD^14–18^. However, the cellular and molecular mechanisms linking circadian disruption to neurodegenerative disease remain unclear.

The brain’s innate immune system contributes to pathogenic protein clearance^19–21^, and mounting evidence from peripheral tissues indicates that innate immunity is regulated by the circadian clock. In peripheral macrophages, toll-like receptor expression, bacterial phagocytosis, and the production of cytokines and matrix metalloproteinases have all been found to occur under circadian regulation^22–28^. Microglia, the parenchyma-dwelling leukocyte of the central nervous system (CNS), exhibit diurnal oscillations in gene expression, inflammatory signaling, and amyloid-beta (Aβ) phagocytosis^29,30^. However, large knowledge gaps remain regarding diurnal rhythms in brain immunity: little to no research has interrogated circadian rhythms in non-microglial immune cells of the CNS despite emerging evidence that border-dwelling macrophages may help prevent Aβ accumulation^31,32^.

To address this gap, we generated a single-cell RNA-sequencing (scRNA-seq) atlas of brain immune niches every six hours around the day-night cycle, enriched for non-microglial leukocytes. We identified brain border-associated macrophages (BAMs) as highly rhythmic immune cells in both transcriptional phenotype and function. BAMs populate the leptomeninges and perivascular spaces of the brain and, unlike monocyte-derived macrophages (MDMs), are long-lived cells of embryonic origin^33–35^. We show that BAMs are highly effective scavengers of fluid-borne substrates including Aβ, far surpassing other brain macrophages in acute engulfment assays. We find that BAM engulfment exhibits diurnal rhythmicity: it is heightened during the rest phase, regulated by the clock gene *Bmal1*, and accompanied by a coordinated wave of upregulation of endocytic genes that is recapitulated in other organs. Furthermore, we identify the endocytic receptor CD206 as a mediator of the rapid scavenging capacity of BAMs and its rhythmicity that is vulnerable to aging-related downregulation. Finally, we show that *Bmal1* deletion in BAMs in the 5xFAD mouse model of AD increases early perivascular and leptomeningeal Aβ plaque deposition. Together, our scRNA-seq atlas and functional studies nominate endocytosis as a specialized and rhythmic function of brain border tissue-resident macrophages. These results add to a handful of studies nominating BAMs as regulators of Aβ plaque deposition^32,36^ and link circadian rhythms in innate immunity to the pathogenesis of amyloidosis such as AD.

## RESULTS

### A single-cell atlas reveals diurnal rhythmicity in the brain immune compartment

To comprehensively characterize how brain immune niches change around the clock, we first performed single-cell RNA sequencing (scRNA-seq) of the brain and leptomeninges of healthy adult mice at Zeitgeber time (ZT) points ZT0, ZT6, ZT12, and ZT18 (**Figure 1A**). Zeitgeber Time refers to the number of hours elapsed from the beginning of the light cycle; this approach thus sampled tissue around a single 24-hour light-dark cycle covering the rest phase (ZT0, ZT6) and active phase (ZT12, ZT18) of mice, a nocturnal species. To sensitively sample rarer immune populations including BAMs and infiltrating leukocytes, we used fluorescence-activated cell sorting (FACS) in combination with acute labeling of circulating immune cells to de-enrich microglia while still excluding contamination from blood-borne leukocytes (**Figure S1A,B**). To arrest cellular transcription at the time of sample collection, prevent dissociation-induced gene signatures, and ensure transcriptional profiles were faithful to collection time, samples were treated with validated transcriptional and translational inhibitors^37^ throughout the collection pipeline.

**Figure 1.**
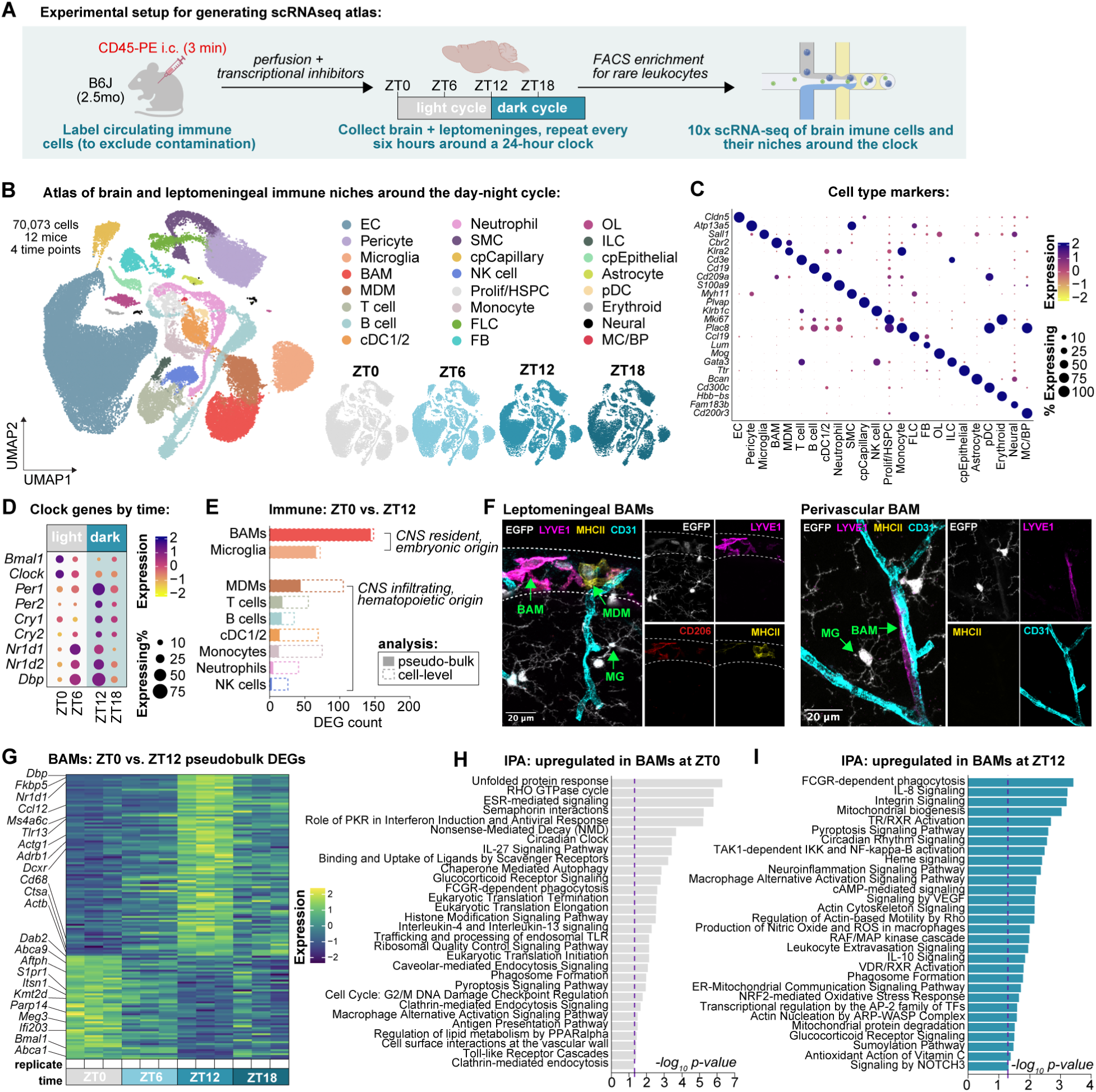
A single-cell atlas reveals diurnal rhythmicity in the brain immune compartment (**A**) Experimental approach. Circulating CD45+ cells were acutely labeled prior by intracardiac (i.c.) injection to transcardial perfusion to identify and exclude any blood contamination during FACS. Illustration made with BioRender. **(B and C**) UMAP of cell types and heat map of cell type markers (n = 12 mice, 3 independent samples per time point). EC = endothelial cell, BAM = border-associated macrophage, MDM = monocyte-derived macrophage, cDC1/2 = classical dendritic cell type 1/2, SMC = smooth muscle cell, cp = choroid plexus, NK = natural killer, HSPC = hematopoietic stem/progenitor cell, FLC = fibroblast-like cell, FB = fibroblast, OL = oligodendrocyte lineage, ILC = innate lymphoid cell, pDC = plasmacytoid DC, MC/BP = mast cell/basophil. Inset plot shows atlas split by time. Note: relative cellular representation in this dataset cannot be used as a proxy for *in vivo* cell counts due to FACS enrichment strategy. (D) Heat map of clock gene expression across time (n = 3 mice per time point, independent samples). (E) Number of differentially expressed genes (DEGs) per immune cell type between ZT0 and ZT12. Analysis was done both via pseudobulk (filled bars, n = 3 independent replicates per time point) and via cell-level analysis (dashed bars, cell number matched across cell types via randomized sub-sampling for analysis, see **Methods**). (F) Immunohistochemistry of leptomeningeal and perivascular BAMs from *Cx3cr1*^EGFP^ mice with intravenously delivered CD31-BV421 prior to cardiac perfusion. BAMs are EGFP^lo^, LYVE1^+^, CD206^+^, and MHC-II^lo^ or negative. MDMs are EGFP^lo^, LYVE1^-^, CD206^lo^, MHC-II^hi^. At steady state, cortical microglia (MG) are EGFP^hi^, LYVE1^-^, CD206^-^, and MHC-II^-^. Dashed line denotes leptomeninges. (G) Heat map of all ZT0 vs. ZT12 DEGs within BAMs (from pseudobulk analysis per replicate, n = 3 independent mice per time point, see **Methods** and **Table S2**). Selected genes of interest highlighted. (**H and I**) Ingenuity Pathway Analysis (IPA; Qiagen) of DEGs (from cell-level analysis) upregulated in BAMs at ZT0 and ZT12. Selected pathways shown, full list in **Table S3**. FCGR = Fcγ receptor, ROS = reactive oxygen species, TF = transcription factor. Minimum of 2 defining genes per pathway. *See also **Figure S1** and **Tables S1-3***.

Following quality control and doublet removal, we identified 70,073 cells (**Figure 1B,C, Table S1**) composed of both immune cells and cells composing their niches from three biological replicates per time point. Immune cells sampled included microglia, BAMs, MDMs, T cells, B cells, conventional dendritic cells (cDC1/2), neutrophils, natural killer (NK) cells, and monocytes, as well as a small number of plasmacytoid DCs, CCR7+ DCs, and hematopoietic stem and progenitor cells (HSPCs), innate lymphoid cells (ILCs), erythroid cells, and mast cells/basophils (MC/BP; **Figure S1C-E**). Niche cells sampled included endothelial cells (ECs), pericytes, smooth muscle cells (SMCs), fibroblast-like cells (FLCs), fibroblasts (FBs), choroid plexus capillary and epithelial cells (cpCapillary, cpEpithelial), and a small number of oligodendrocyte lineage cells (OLs), astrocytes, and other neural cells (**Figure S1F-H**). Cells were annotated using markers established from spatial, ontological, and transcriptional profiling of the brain and its borders^35,38–43^. All major cell types were captured across time (**Figure 1B, Figure S1E,H**).

Across all cells, expression of core clock genes and downstream transcription factors were rhythmic: *Bmal1* and *Clock* were highest at ZT0, whereas the antiphase genes *Per1, Per2, Cry1, Cry2, Nr1d1, Nr1d2* and *Dbp* peaked at ZT12 (**Figure 1D**). These data indicate that robust diurnal rhythms in core circadian clock machinery were successfully captured in our atlas. We next compared how different brain-resident immune cell types changed around the clock. BAMs had the highest number of differentially expressed genes between ZT0 and ZT12 (**Figure 1E**), surpassing all other leukocytes and agnostic to whether this analysis was done on the level of individual cells or using a pseudobulk analysis approach (**Figure 1E**, see **Methods**). For the remainder of this manuscript, we will focus on describing the different ways in which the circadian clock acts to regulate BAMs as well as the consequences of disrupting this process; however, there are many other interesting changes occurring in other cell types which will be important to investigate in future studies.

BAMs are long-lived, yolk sac-derived macrophages that reside in the leptomeninges and along brain vasculature and can be distinguished by expression of the mannose receptor CD206 (encoded by *Mrc1*, **Figure 1F**). Transcriptionally, BAMs are identifiable by co-expression of genes including of *Ms4a7, Cd38, Lyve1, Ccl24, Egfl7,* and *Clec4n,* and de-enrichment for both microglia markers (*Sall1, Sparc, Serpine2*) and MDM markers (*Ccr2, Klra2;* **Figure S1I-J**). Pseudobulk comparison of BAMs at ZT0 versus ZT12 identified 144 differentially expressed genes (DEGs; **Figure 1G, Table S2**). At ZT12, BAMs upregulated genes involved in chemokine signaling (*Ccl12, Cmklr1*), innate immune sensing (*Tlr13*), phagocytosis (*Ncf1, Rac3, Rab7b, Cd68*), actin dynamics (*Actg1, Actb*), and glucocorticoid signaling (*Fkbp5, Sgk1*), as well as the antiphase clock (*Dbp, Per1, Per3, Nr1d1, Nr1d2*) and MS4 family genes (*Ms4a4a, Ms4a6c, Ms4a6b, Ms4a6d*). At ZT0, BAMs instead upregulated genes involved in DNA repair (*Parp14, Kmt2d*), chromatin remodeling (*Chd3*), lipid homeostasis (*Abca1, Abca9*), proteasomal quality control (*Hspa5, Hspa8*), and endocytosis (*Dab2, Itsn1, Aftph*), as well as the positive branch of the circadian feedback loop (*Bmal1*). Cell-level analysis and subsequent Ingenuity Pathway Analysis (IPA; Qiagen) on genes upregulated by BAMs at ZT0 versus ZT12 identified similar circadian segregation of function (**Figure 1H,I; Table S3**). BAMs at ZT12 were enriched for pathways including reactive oxygen species (ROS) production, neuroinflammatory signaling, oxidative stress response, and leukocyte extravasation whereas BAMs at ZT0 were enriched for canonical tissue repair functions including aggrephagy, ribosomal quality control, and chaperone-mediated autophagy. ZT0 was also enriched for pathways involving the binding and uptake of ligands by scavenger receptors and endocytosis. Together, these patterns of gene and pathway enrichment are consistent with a model whereby BAMs are primed tissue defense during the active phase and tissue repair during the rest phase, and nominate engulfment as a rhythmic BAM function.

### BAMs rapidly and rhythmically scavenge fluid-borne substrates

BAMs reside along routes of fluid egress from the brain where they sample draining parenchymal substrates. One such substrate is Aβ, a peptide involved in AD pathogenesis with which BAMs colocalize in AD tissue^44,45^. Aβ accumulates in brain interstitial fluid (ISF) during wakefulness and peaks in cerebrospinal fluid (CSF) at the onset of the rest phase^17,46,47^. Converging evidence from several model organisms also indicates that fluid flow in the brain is increased during sleep^48–52^. We thus hypothesized that BAMs may be rhythmically scavenging draining parenchymal peptides such as Aβ, and that this process may be heightened during the rest phase. To test this, we turned to the 5xFAD mouse model of AD, which exhibits progressive amyloidosis with plaque formation beginning around 3 months of age^53^. We opted for an early time point in disease pathogenesis (2 months of age) to preclude effects of plaque formation on drainage dynamics of Aβ. We measured *in vivo* engulfment using FEAST^54^, a flow cytometry assay relying on perfusion fixation prior to tissue dissociation and staining of intracellular material to ensure antigens are not taken up *ex vivo.* In the 5xFAD line, expression of human APP and PSEN1 is driven by the neuronal Thy1.2 promoter^55^. Using the 5xFAD mouse model – instead of alternatives such as the *App^NL-G-F^*line – ensures that Aβ detected within brain macrophages using a human-specific anti-Aβ antibody (clone 6E10) is exclusively neuronally-derived and engulfed (but not produced) by macrophages.

We performed FEAST on cortical/leptomeningeal microdissections from young adult male pre-plaque 5xFAD mice (aged P61-63; **Figure 2A**) at either the peak of the rest phase (ZT6) or the peak of the active phase (ZT18). Aβ was broadly detected within all macrophage subsets of the brain – BAMs, MDMs, and microglia – but signal intensity was highest within BAMs (**Figure S2A**). BAMs exhibited increased Aβ engulfment during the rest phase relative to the active phase, both in terms of the percent of Aβ+ BAMs and in the intensity of Aβ signal (**Figure 2B-C**). This effect was restricted to BAMs: microglia showed no significant difference in Aβ engulfment across time (**Figure S2B-C**), and even a slight trend toward heightened signal during the active phase as has been observed in later stages of amyloidosis^30^. Taken together, these data indicate that BAMs exhibit diurnal variation in Aβ uptake *in vivo*.

**Figure 2.**
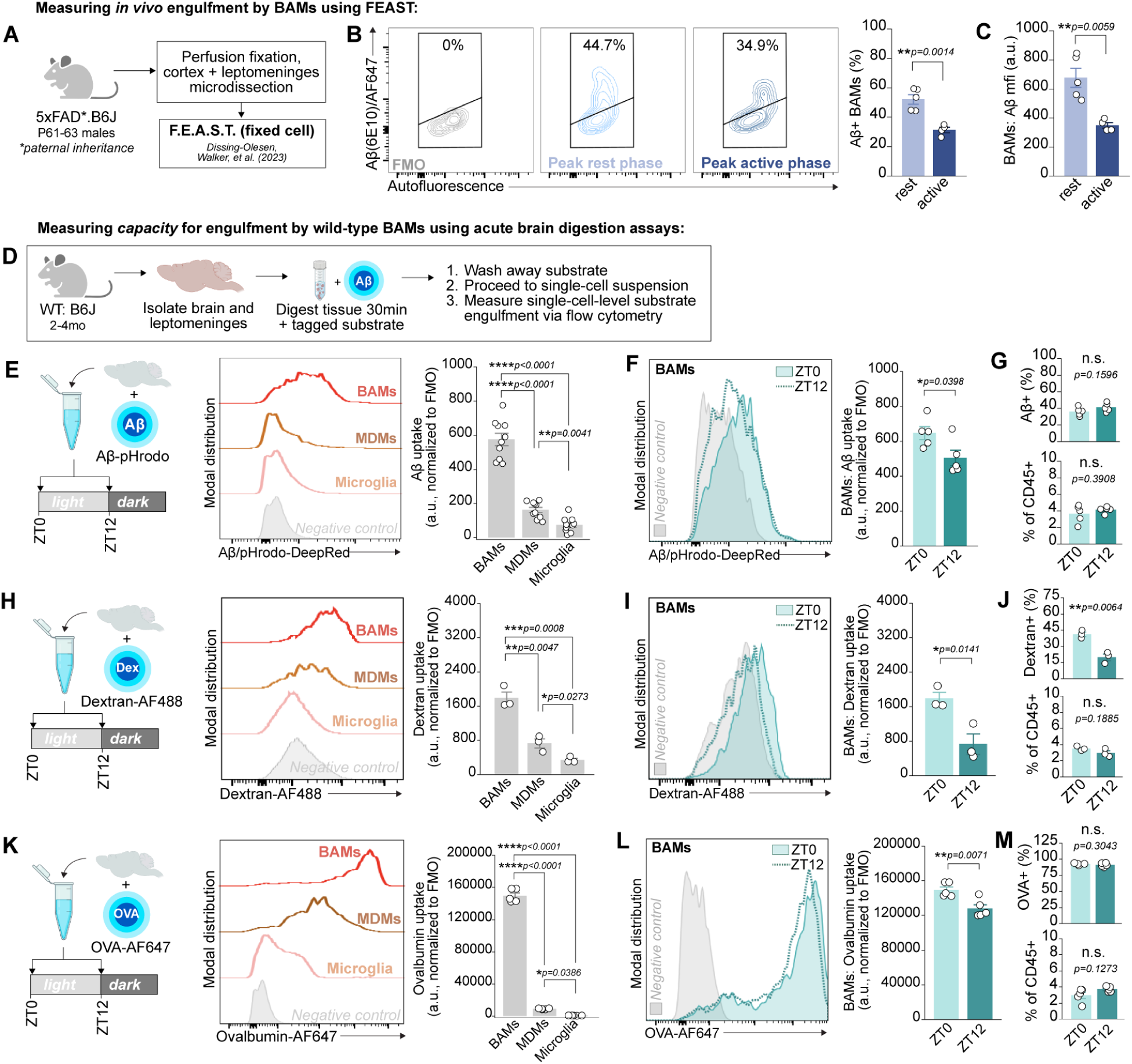
BAMs rapidly and rhythmically scavenge fluid-borne substrates **(A-C)** Experimental design for assessment of *in vivo* Aβ engulfment using FEAST, representative contour plots of Aβ signal within BAMs during the rest versus active phase, and group-level quantification of Aβ engulfment by BAMs. Both Aβ positivity and signal intensity in BAMs are increased during the rest phase (t(6.12)=5.53, p=0.0014 and t(4.69)=4.78, p=0.0059, respectively). Comparisons with two-tailed t-tests, n of 4-5 mice per time point. **(D)** Experimental design for assessing engulfment capacity of live brain macrophages using acutely isolated brain tissue. **(E)** Experimental design, representative histograms, and group-level quantification of Aβ uptake by cell type. Acute Aβ engulfment capacity is increased in BAMs relative to microglia and MDMs (F_2,18_=256.56, p<0.0001; mixed effects model with random variable of sample; n of 10 mice). **(F and G)** Representative histogram and group-level quantification of Aβ uptake by BAMs, the percent of Aβ+ BAMs, and BAM yield across time. Aβ uptake by BAMs is reduced at ZT12 relative to ZT0 (t(7.33)=-2.54, p=0.0372), but both Aβ-positivity and BAM yield are unchanged (p=0.1596 and p=0.3908, respectively). All comparisons with two-tailed t-tests, n of 5 mice per time point. (H) Experimental design, representative histogram, and group-level quantification of Dextran uptake by cell type. Acute Dextran engulfment capacity is increased in BAMs relative to microglia and MDMs (F_2,4_=65.97, p=0.0009; mixed effects model with random variable of sample; n of 3 mice). (**I and J**) Representative histogram and group-level quantification of Dextran uptake by BAMs, Dextran+ BAMs, and BAM yield across time. Dextran uptake and Dextran+ BAMs are reduced at ZT12 relative to ZT0 (t(3.84)=-4.27, p=0.0141 and t(3.56)=-5.75, p=0.0064, respectively), but BAM yield is unchanged (p=0.1885). All comparisons with two-tailed t-tests, n of 3 mice per time point. (**K**) Experimental design, representative histogram, and group-level quantification of OVA-AF647 uptake by cell type. Acute OVA-AF647 engulfment capacity is increased in BAMs relative to microglia and MDMs (F_2,12_=1582.822, p<0.0001; mixed effects model with random variable of sample; n of 5 mice). (**L and M**) Representative histogram and group-level quantification of OVA-AF647 uptake by BAMs, OVA+ BAMs, and BAM yield across time. OVA-AF647 uptake by BAMs is reduced at ZT12 relative to ZT0 (t(7.56)=-3.66, p=0.0071), but both OVA-positivity and BAM yield are unchanged (p=0.3043 and p=0.1273, respectively). All comparisons with two-tailed t-tests, n of 5 mice per time point. *For all panels: Unless otherwise specified, bars and error bars represent mean and SEM. Points represent individual mice. All post-hoc testing with Tukey’s HSD. Illustrations made with BioRender*. *\*\*\*\*p<0.0001, ***p<0.001, **p<0.01, *p<0.05. See also **Figure S2***.

Circadian rhythms enable changes in cellular function and physiology in anticipation of environmental changes rather than in response to them. We thus sought to interrogate whether the cell-intrinsic capacity of BAMs to engulf fluid-borne substrates may also be rhythmic, independent of changes in Aβ production and drainage^14,17,47,56–58^ as well as other variables including diurnal changes in lysosomal properties and epitope degradation (**Table S3**). To this end, we designed rapid engulfment assays wherein the brain and leptomeninges of wild-type mice were acutely isolated and immediately digested with the addition of an exogenous and fluorescently-tagged substrate for 30 minutes (**Figure 2D**; see **Methods**). Following tissue digestion, the substrate was washed away and its engulfment was measured on the single-cell level via flow cytometry. BAMs were identified using an antibody panel validated with our scRNA-seq atlas (**Figure S1K-M**). This approach allowed us to control both exposure duration and substrate concentration, thus measuring the intrinsic capacity of a cell to engulf extracellular material with the only variable being time of day. We also confirmed using primary cultured BAMs isolated from a reporter line (*Pf4*^Cre^ x Ai14) that BAMs were able to engulf Aβ in this time scale (**Figure S2D**). Within just 30 minutes of incubation with AF647-conjugated Aβ_1-42_, substantial Aβ signal localized to intracellular vesicles was visible *in vitro*. This confirmed that BAMs are capable of rapid substrate uptake on the order of minutes rather than hours and that acute engulfment assays measure a physiologically-relevant method of BAM scavenging.

Thus, we acutely digested wild-type brains for 30 minutes in the presence of pHrodo-conjugated synthetic Aβ_1-42_ fibrils at ZT0, the onset of the rest phase, versus ZT12, the onset of the active phase. This approach ensured that cells were exposed to an equal amount of Aβ_1-42_ across conditions and that fluorescent signal would only appear after substrate exposure to the acidic conditions of the endolysosomal compartment. BAMs by far surpassed both MDMs and microglia in Aβ uptake (**Figure 2E**). BAMs on average engulfed 4-fold more Aβ_1-42_ than MDMs and 11-fold more Aβ_1-42_ than microglia. Furthermore, we found that BAM scavenging was rhythmic: BAMs engulfed on average 28% more Aβ_1-42_ at ZT0 relative to ZT12 (**Figure 2F**). This difference was not likely due to any differences in fibril diffusion in the digest, as both the percentage of Aβ+ BAMs and BAM yield were unchanged across time (**Figure 2G**). Thus, BAM engulfment of Aβ is rhythmic and enhanced during the murine rest phase.

We then turned to other substrates to assess the replicability and dynamics of rhythmicity in BAM engulfment. In rapid engulfment assays using either AF488-conjugated Dextran (Dextran-AF488) or AF647-conjugated Ovalbumin (OVA-AF647), BAMs again exhibited vastly elevated substrate uptake relative to MDMs and microglia (**Figure 2H,K**). This extremely rapid scavenging capacity of BAMs was reminiscent of the highly endocytic function of peripheral CD206+ cell types including dendritic cells, liver sinusoidal endothelial cells (LSECs), and vascular-associated fat macrophages (VAMs)^59–62^. Indeed, IPA on genes upregulated in BAMs versus other CNS macrophages highlighted their enrichment for transcripts related to clathrin-mediated endocytosis (CME), a form of engulfment that can internalize ligands including ovalbumin via the BAM-enriched mannose receptor CD206 on timescale of minutes, nominating CME as a likely route of rapid uptake of fluid-borne substrates by BAMs. When larger engulfment substrates were used, such as AF488-conjugated *E.coli* BioParticles that exceed the cargo limit for CME and require phagocytic uptake, BAMs showed far reduced uptake within the 30-minute assays and no longer exhibited heightened uptake relative to MDMs and microglia (**Figure S2F-G**). Thus, BAMs are expert scavengers of extracellular fluid-borne antigens and share features of highly endocytic cells.

Rhythmicity in BAM engulfment also replicated across substrates and varying experimental conditions Uptake of both Dextran-AF488 and OVA-AF647 was elevated in BAMs harvested at ZT0 relative to ZT12, with no difference in BAM yield to explain this difference (**Figure 2I-M**). Ovalbumin uptake by BAMs was reduced at ZT12 relative to both ZT0 and ZT6, suggesting that increased scavenging is a persistent feature of BAMs during the rest phase (**Figure S2H**). While heightened BAM engulfment at ZT0 replicated across all substrates (Aβ, Dextran, Ovalbumin), this was not the case for microglia and MDMs (**Figure S2I**), suggesting that rhythmicity in rapid scavenging capacity is specifically a distinctive feature of BAM functionality. The diurnal change in BAM engulfment was highly consistent across experimental replicates: ovalbumin uptake was on average reduced by 27.5% at ZT12 relative to ZT0 (95% confidence interval (CI): 21.9-33.1%; **Figure S2J**). The diurnal change replicated whether assays were performed in parallel from reverse light cycle rooms or in sequence from the same animal room, suggesting that the change of engulfment was attributable to neither cage changing schedules nor housing room foot traffic (**Figure S2K**). Enhanced engulfment at ZT0 versus ZT12 was also maintained across sexes and at a different research institution (**Figure S2L,M**). Taken together, these data indicate that BAMs are expert and rhythmic scavengers of small, fluid-borne substrates. Unlike phagocytosis, which peaks in macrophages during the active phase^26^, we find that the capacity for endocytic uptake of small cargo is heightened during the rest phase in BAMs.

### Diurnal change in BAM engulfment is regulated by the clock gene *Bmal1*

Circadian rhythms persist in the absence of external lighting cues. To interrogate the source of rhythmicity in BAM engulfment, we first tested whether the diurnal change persisted in the absence of external lighting cues by keeping mice in total darkness after the expected onset of the light cycle. Under constant conditions, mice showed reduced ovalbumin uptake at Circadian Time 12 (CT12, referring to the hours elapsed since anticipated time of light exposure) relative to CT0 (**Figure 3A**). We also assessed whether the diurnal change may be sleep-dependent by subjecting mice to acute and total sleep deprivation via gentle handling and novel object introduction (see **Methods**). Mice were sleep deprived either for the first six hours of the light cycle (ZT0 to ZT6) or the six hours leading up to the dark cycle (ZT6 to ZT12). Sleep deprivation had no effect on the reduction in OVA-AF647 uptake by BAMs between ZT6 and ZT12 (**Figure 3B**). However, we did observe a circadian-independent effect of acute sleep deprivation on BAM engulfment, suggesting that multiple environmental factors influence BAM engulfment beyond its rhythmicity. Together, these experiments indicate that the rhythmic scavenging of extracellular ovalbumin by BAMs persists in the absence of both light cues and sleep, suggesting that it may instead be under direct regulation of circadian clock machinery.

**Figure 3.**
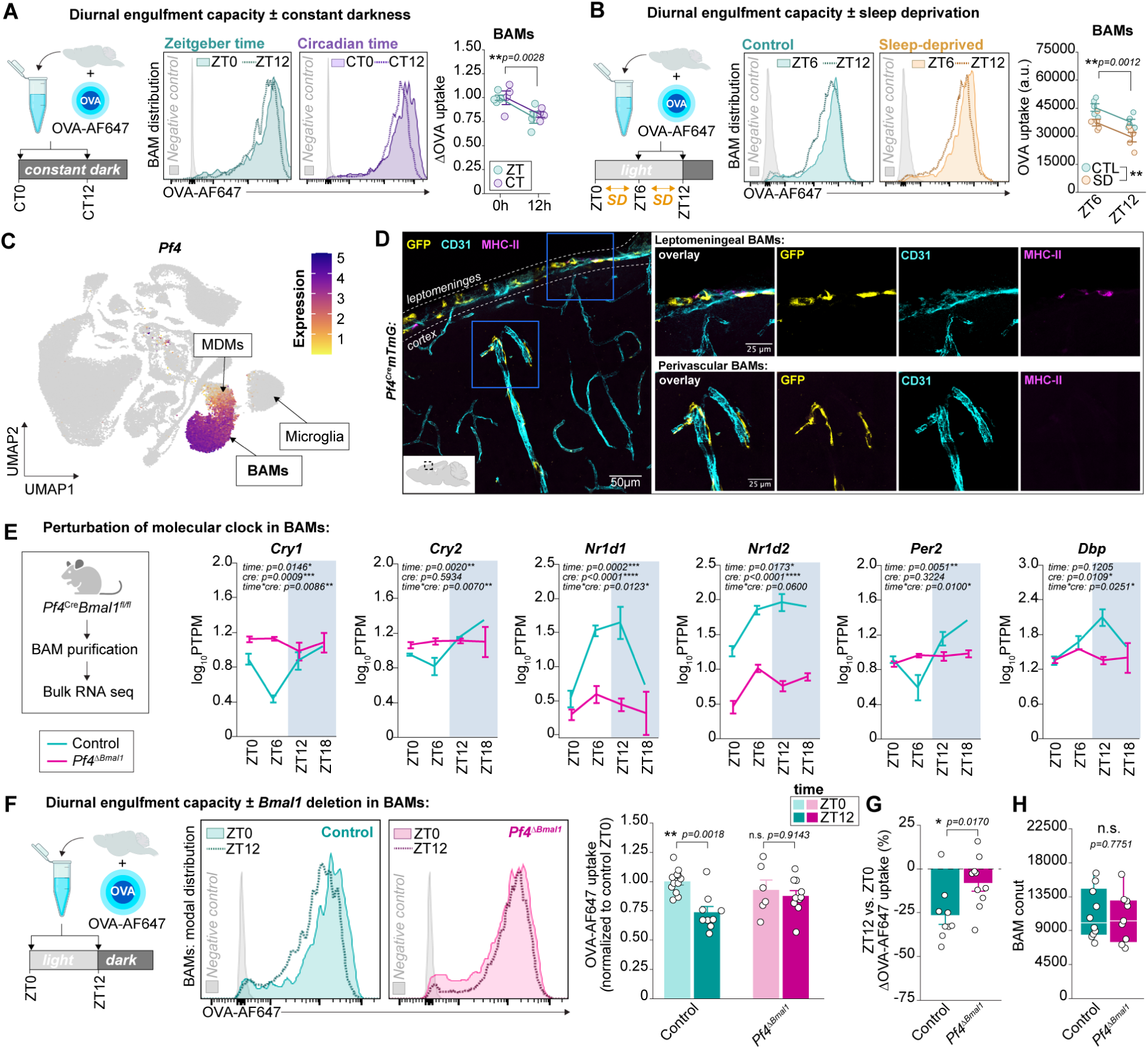
Diurnal change in BAM engulfment is regulated by the clock gene *Bmal1* (A) Experimental design, representative histograms, and group-level comparison of OVA-AF647 uptake by BAMs from mice under constant conditions versus controls. Main effect of time but not condition on OVA-AF647 uptake (two-way ANOVA, n of 4 per condition per time point). OVA-AF647 uptake is normalized to the 0h mean per condition. (B) Experimental design, representative histograms, and group-level comparison of OVA-AF647 uptake by BAMs following 6 hours of total sleep deprivation (SD). Main effect of time (p=0.0012) and condition (p=0.0010), but no interaction effect between the two (p=0.9988) on OVA-AF647 uptake (two-way ANOVA; n of 5 per condition per time point). (**C and D**) Feature plot of *Pf4* expression within single-cell atlas showing expression restricted to BAMs, a subset of MDMs, and a small number of proliferating cells/HSPCs, and IHC showing perivascular and leptomeningeal GFP+ cells from *Pf4*^Cre^*mTmG* mice. (E) Experimental design for validation of circadian perturbation and group-level quantification of pseudo-transcripts per million (PTPM) for clock genes of interest in BAMs from *Pf4^ΔBmal1^* mice versus their littermate controls. Shaded areas represent dark cycle, line and error bars represent mean and SEM per genotype per time point. Model results shown are from two-way ANOVA between time and genotype. (F) Experimental design for diurnal engulfment assay, representative histograms, and group-level quantification of OVA-AF647 uptake by BAMs in *Pf4^ΔBmal1^* mice versus their littermate controls. Genotype*time interaction: p=0.0425 (two-way ANOVA). Results pooled from two independent experiments (n of 21 control and 16 *Pf4^ΔBmal1^* mice). (G) Group level quantification of % change in OVA-AF647 uptake between ZT0 and ZT12 in BAMs from *Pf4^ΔBmal1^* mice versus their littermate controls (comparison using student’s t-test). Only the ΔOVA-AF647 uptake from control, but not *Pf4^ΔBmal1^*BAMs, is significantly non-zero (one-sample t-tests against hypothesized mean of zero, p=0.0009 and p=0.1136, respectively). Results shown from two independent experiments. Points represent individual mice (n of 9-10 per genotype). (H) BAM count from controls versus *Pf4^ΔBmal1^* mice. Cell count is unchanged by genotype (p=0.7751, Mann-Whitney U test). Points represent individual mice (n of 9-10 per genotype). *For all panels: Bars and error bars represent mean and SEM. Box and whisker plots represent median and IQR. Post-hoc testing with Tukey’s HSD. Illustrations made with BioRender. ****p<0.0001, ***p<0.001, **p<0.01, *p<0.05. See also **Figure S3***.

To test whether the molecular clock regulates diurnal changes in BAM engulfment, we generated mice lacking *Bmal1* in BAMs. Differential gene expression analysis across our scRNA-seq atlas nominated *Pf4* as a BAM-enriched gene, and the *Pf4*^Cre^ line had previously been shown to target BAMs^63,64^ (**Figure 3C**). When bred to a reporter line expressing membrane-bound GFP in the presence of Cre (*Pf4*^Cre^*mTmG*), we observed robust recombination in almost all BAMs, around 25% of MDMs, and less than 2% of microglia (**Figure 3D**, **Figure S3A-B**). We did not observe GFP expression above background noise in any other cells yielded by flow cytometry (**Figure S3B**). Because a bacterial artificial chromosome (BAC) was used to generate the *Pf4^Cre^* line^65^, we also assessed whether the BAC contents alone could confound any readouts. We found no effect of BAC presence on engulfment capacity of brain macrophage subsets nor on the number of microglia, BAMs, MDMs, nor GR-1+ cells in the brain (**Figure S3C-E**). Having established that the *Pf4*^Cre^ line was suitable for BAM-targeted genetic manipulations, we crossed it to the *Bmal1^fl/fl^* line^66^, where the eighth exon of *Bmal1* is flanked with *loxP* sites, to obtain mice lacking BMAL1 in *Pf4-*expressing cells (*Pf4^ΔBmal1^*). We validated that this approach successfully disrupted BAM circadian rhythms via bulk RNA sequencing of FACS-purified BAMs at six hour increments around the day-night cycle from *Pf4^ΔBmal1^* mice and their littermate controls (**Figure 3E, Figure S3F,G**). BAMs from *Pf4^ΔBmal1^* mice lost rhythmicity in the expression of multiple core clock genes including *Cry1, Cry2, Nr1d1, Nr1d2, Per2,* and *Dbp* (**Figure 3E**) and lacked reads mapping to the loxP-flanked eighth exon of *Bmal1* (**Figure S3G**). Thus, *Pf4^ΔBmal1^*mice are suitable for studying the effects of disrupting circadian rhythms in BAMs with minimal off-target effects in other brain and leptomeningeal cells.

We then performed acute OVA-AF647 engulfment assays on brains from *Pf4^ΔBmal1^* mice and their littermate controls at ZT0 and ZT12 (**Figure 3F**). *Bmal1* deletion in BAMs abolished the diurnal change in engulfment (**Figure 3F-G**), despite having no effect on the overall percentage of BAMs engulfing (**Figure S3H**). In control BAMs, OVA-AF647 uptake was reduced on average 26.40% (±5.18%; mean±SEM) at ZT12 relative to ZT0 (**Figure 3G**). In BAMs from *Pf4^ΔBmal1^* mice, the change between ZT0 and ZT12 was not significantly different from zero (8.04±4.59%; **Figure 3G**). Importantly, BAM count was unchanged by *Bmal1* deletion (**Figure 3H**), indicating that the loss of rhythmicity is not likely due to a change in cell viability or population size. Taken together, these data indicate that diurnal rhythmicity in BAM engulfment depends on *Bmal1* and likely, an intact circadian transcription-translation feedback loop.

### The mannose receptor CD206 mediates rapid engulfment by BAMs and its rhythmicity

We then explored the cellular mechanisms that may underlie the rapid scavenging capacity of BAMs and its rhythmicity by returning to our scRNA-seq atlas with a specific interest in genes upregulated at ZT0, the time of heightened engulfment capacity. To leverage the single-cell resolution and increase our power to detect small magnitude changes around the clock, we used MAST^67^, a gene enrichment analysis approach using a generalized linear framework that controls for gene detection rate per cell. The results were further filtered for time-of-day effects: DEGs were only included if significantly different at a minimum of two time points with a peak and trough in expression (see **Methods**). This pipeline successfully identified diurnal shifts in clock gene expression (e.g. *Bmal1, Clock*; **Table S4**) despite low cellular detection rates, highlighting the strength of this approach. This DEG analysis nominated multiple genes involved in clathrin-mediated endocytosis (CME), a highly efficient route by which macrophages internalize extracellular ligands (**Figure 4A**). Upregulated genes encoded components of the clathrin coat (*Cltc*), adaptor proteins and machinery involved in cargo recognition (*Ap2b1, Fcho2, Dab2, Cd2ap, Clint1, Itsn1*), scaffold proteins (*Eps15, Eps15l1*), proteins involved in vesicular trafficking (*Scamp1, Aftph*), and receptors known to internalize extracellular ligands via CME (*Mrc1, Msr1*). K-means clustering of BAMs from all scRNA-seq samples using only these 13 genes successfully ranked 10 out 12 samples by time, the remaining two samples were only off by a single time point (**Figure S4A**). BAMs from *Pf4^ΔBmal1^* mice exhibited perturbed rhythmicity in several of these transcripts (**Figure S4B**), suggesting that they may be under circadian regulation. To further interrogate the role of the molecular clock in regulating CME-associated genes, we re-analyzed published data from chromatin immunoprecipitation sequencing (ChIP-seq) of the liver acquired every four hours from CT0 to CT20^68^. Rhythmic BMAL1 binding was detected on multiple ubiquitously-expressed CME-associated genes including *Clint1, Cltc, Eps15l1, Itsn1,* and *Scamp1* with a period averaging 24 hours (24.01±0.32) and peaking during the rest phase (**Figure 4C**). These data closely align with our scRNA-seq and suggest that, across multiple organs, the circadian clock drives a coordinated wave of upregulation of CME-associated genes during the rest phase.

**Figure 4.**
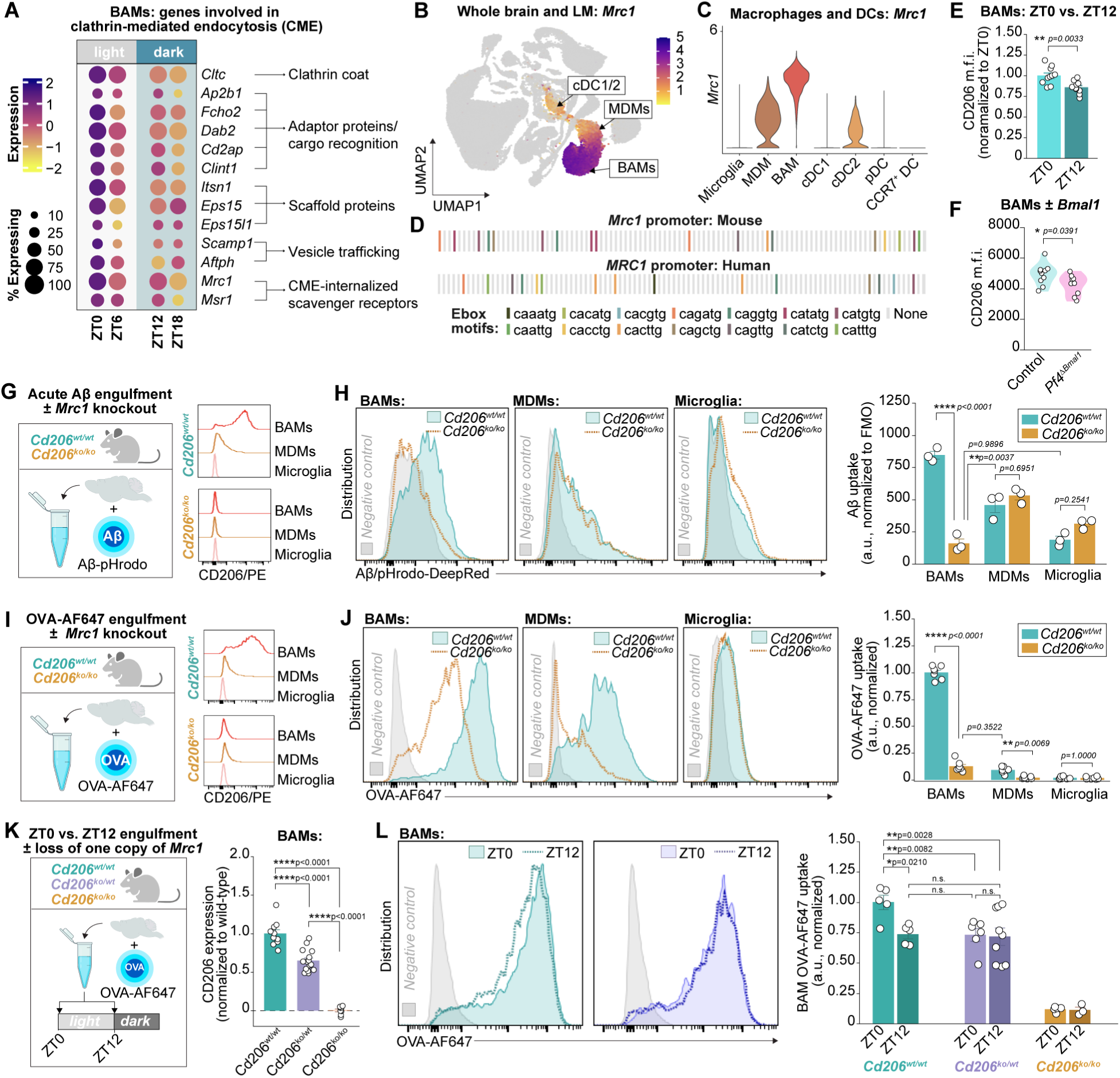
The mannose receptor CD206 mediates rapid engulfment by BAMs and its rhythmicity (A) Heat map of CME-associated gene expression across time in BAMs in scRNA-seq atlas (n of 3 mice per time point). All determined to be DEGs via time-peak filtered MAST analysis (see **Methods**). (**B and C**) *Mrc1* expression in scRNA-seq atlas and across cell types. (D) E-box domain mapping along murine and human *Mrc1/MRC1* promoter. (E) Protein-level CD206 expression is reduced at ZT12 relative to ZT0 in BAMs (t(16.21)=-3.44, p=0.0033; two-tailed t-test). Data are pooled from two independent replicate experiments, n of 10 mice per time point. (F) Protein-level CD206 expression is reduced in *Pf4^ΔBmal1^* mice versus their littermate controls (t(16.90)=-2.24, p=0.0391; two-tailed t-test; n of 9-10 mice per genotype). (**G and H**) Experimental design, representative histograms, and group-level quantification of Aβ-pHrodo uptake per cell type from *Cd206^ko/ko^*mice and littermate controls. Cell type*genotype interaction effect: p<0.0001 (mixed effect model with random effect of mouse; n of 3 mice per genotype). (**I and J**) Experimental design, representative histograms, and group-level quantification of OVA-AF647 uptake per cell type from *Cd206^ko/ko^*mice and littermate controls. Cell type*genotype interaction effect: p<0.0001 (mixed effect model with random effect of experimental replicate and mouse). N of 7 mice per genotype, results pooled across three experimental replicates. (K) Experimental design and group-level quantification of CD206 expression on BAMs per genotype. Main effect of genotype on CD206 expression: p<0.0001 (mixed effects model with random variable of experimental replicate). N of 8-15 mice per genotype, results pooled from two experimental replicates. (L) Representative histograms and quantification of OVA-AF647 uptake by BAMs at ZT0 vs. ZT12 across genotypes. Time*genotype interaction: p=0.0380 (mixed effects test with random effect of experimental replicate). N of 3-9 mice per genotype per time point, results pooled from two experimental replicates. *For all panels: points represent individual mice, bars and error bars represent mean and SEM. Post-hoc testing with Tukey’s HSD. Illustrations made with BioRender. ****p<0.0001, ***p<0.001, **p<0.01, *p<0.05. See also **Figure S4, Table S4***.

We were particularly interested in the nomination of *Mrc1* as upregulated at ZT0 given that it is a defining feature of other highly endocytic cell types^59–62^. *Mrc1* encodes the rapidly recycling macrophage mannose receptor CD206 that can internalize extracellular ligands on the order of minutes^61,62^. In the brain, *Mrc1* is primarily expressed by BAMs, though a low level of expression is detected in MDMs and cDC2s (**Figure 4B-C**). The actual representation of CD206+ cells in the CNS is by far enriched for BAMs, which accounted for ∼90% of all CD206+ cells yielded from the brain and leptomeninges of healthy young adult mice following enzymatic digestion (**Figure S4E**). *Mrc1* exhibits high levels of evolutionary conservation across mammals^69^ and both the murine and human promoter regions of *Mrc1* are enriched for E-box motifs (**Figure 4D**), the binding sites for the BMAL1-CLOCK heterodimer and clock-controlled transcription factors that drive rhythmic gene expression. A 24-hour rhythm in *Mrc1* expression peaking during the rest phase had also been observed in peritoneal macrophages^26^, and ChIP-seq analysis of the liver has identified rhythmic Ser5P binding to the *Mrc1* promoter^68^. Ser5P labels the C-terminal domain of the RNA polymerase RNAPII during transcription initiation, thus indicating *de novo* transcription. In line with our results, rhythmic Ser5P recruitment to the *Mrc1* promoter in the liver peaked at CT0 with a period of 24.19 hours (**Figure 4D**)^68^, demonstrating that rhythmic *Mrc1* transcription is recapitulated in other organs and likely driven by *de novo* transcription that is heightened at the onset of the rest phase. On the protein-level, CD206 expression was similarly higher at ZT0 versus ZT12 in BAMs and was reduced overall in BAMs lacking BMAL1 (**Figure 4E,F**). Together, our transcriptional and protein-level data, along with converging lines of evidence from other research groups, indicate that *Mrc1* and its encoded receptor CD206 are rhythmic with a peak during the rest phase.

To test whether *Mrc1* contributes to BAM engulfment and its rhythmicity, we obtained mice lacking CD206^70^ and performed acute Aβ engulfment assays in CD206-knockout mice versus their littermate controls (*Cd206^ko/ko^* and *Cd206^wt/wt^* respectively, **Figure 4G**). Flow cytometric assessment confirmed complete loss of CD206 on BAMs and MDMs from the knockouts (**Figure S4F**). Deletion of CD206 had no effect on the expression of independent markers used for BAM identification (CD38) nor on BAM count (**Figure S4G-H**). Loss of CD206 profoundly impaired the capacity of BAMs to engulf Aβ (**Figure 4H**). BAMs lacking CD206 no longer showed enhanced Aβ engulfment relative to microglia and MDMs. In contrast, Aβ uptake was unchanged in either MDMs or microglia in the absence of CD206, suggesting that this receptor is specifically important for acute BAM scavenging of fluid-borne Aβ. Similarly, CD206-deficient BAMs exhibited nearly a 90% reduction in OVA-AF647 uptake (**Figure 4I-J**). Parallel experiments using *E.coli* BioParticles as a substrate revealed no effect of CD206 deletion on BAM engulfment (**Figure S4I-K**). Together, these data indicate that the macrophage mannose receptor may be a critical component of the cellular machinery used by BAMs to endocytose small fluid-borne substrates, including Aβ, over short time frames.

Given that CD206 accounted for around 90% of the engulfment capacity of BAMs in acute assays and that the dynamic range in engulfment around the clock was nearly 30%, it was evident that at least some of the rhythmicity in BAM engulfment may be related to diurnal changes in CD206. To further explore this link, we digested brains from mice lacking either zero, one, or two copies of *Mrc1* (*Cd206^wt/wt^*, *Cd206^ko/wt^,* and *Cd206^ko/ko^*, respectively) with OVA-AF647 (**Figure 4K**). Loss of a single copy of the gene resulted in approximately a 40% reduction in cell-surface CD206 on BAMs and a 25% reduction in acute OVA-AF647 engulfment relative to wild-type littermates (**Figure 4K-L**). While BAMs from *Cd206^wt/wt^* mice showed a consistent reduction in OVA-AF647 uptake at ZT12 relative to ZT0, rhythmicity was lost in *Cd206^ko/wt^* BAMs (**Figure 4L**). Furthermore, levels of engulfment by BAMs lacking one copy of *Mrc1* at both ZT0 and ZT12 phenocopied those of wild-type BAMs at ZT12. Thus, there is a dose-dependent effect of *Mrc1* expression on both BAM engulfment and its rhythmicity: reduced availability of CD206 perturbs the dynamic range of BAM engulfment around the day-night cycle. It is important to note that the diurnal change in *Mrc1* expression is far smaller than loss of an entire copy of the gene: our scRNA-seq data suggest a 15-20% dynamic range in transcript expression around the clock. Diurnal change in BAM engulfment is likely related to the coordinated upregulation of multiple components of endocytic machinery. These data nominate CD206 as a crucial component of BAM engulfment that is also necessary for its rhythmicity.

### Aged BAMs exhibit perturbed clock gene expression and engulfment capacity

Across species, aging is accompanied by diminished circadian amplitude^71–75^. Aged peritoneal macrophages have been reported to lose circadian rhythmicity in phagocytic function^26^, and aged microglia lose circadian rhythmicity in transcriptional phenotype^16^. To explore whether aging may similarly impact the phenotype and function of BAMs, we performed scRNA-seq of the brain and leptomeninges of aged mice (2 years old; **Figure S5A-C**) in tandem with our scRNA-seq in adult mice (2.5 months old). We sampled aged mice at both ZT0 and ZT6, rather than just ZT0, to ensure any changes would not simply be due to shifted circadian rhythms.

Microglia, BAMs, and MDMs were all detectable in the aged brain (**Figure S5D-E**) and aged BAMs were still distinguishable from MDMs and microglia by their expression of genes including *Cd38, Ms4a7,* and *Lyve1*. Flow cytometric analysis found no difference in BAM count between adult and aged brains, but a slight decrease in microglia and a substantial increase in MDMs (**Figure S5F**). Pseudobulk analysis comparing BAMs from adult versus aged brains revealed 289 differentially expressed genes between the age groups (**Figure 5A, Figure S5G,H, Table S5**). IPA on differentially expressed genes nominated clathrin-mediated endocytosis, fibrin clotting, binding and uptake of ligands by scavenger receptors, caveolar-mediated endocytosis, and extracellular matrix (ECM) organization as downregulated with age, and antigen presentation, interferon gamma signaling, and T cell interactions as upregulated with age (**Figure 5B**). K-means clustering of all differentially expressed genes indicated these signatures were still sufficient to distinguish adult from aged BAMs regardless of time (ZT0 and ZT6; **Figure 5I**).

**Figure 5.**
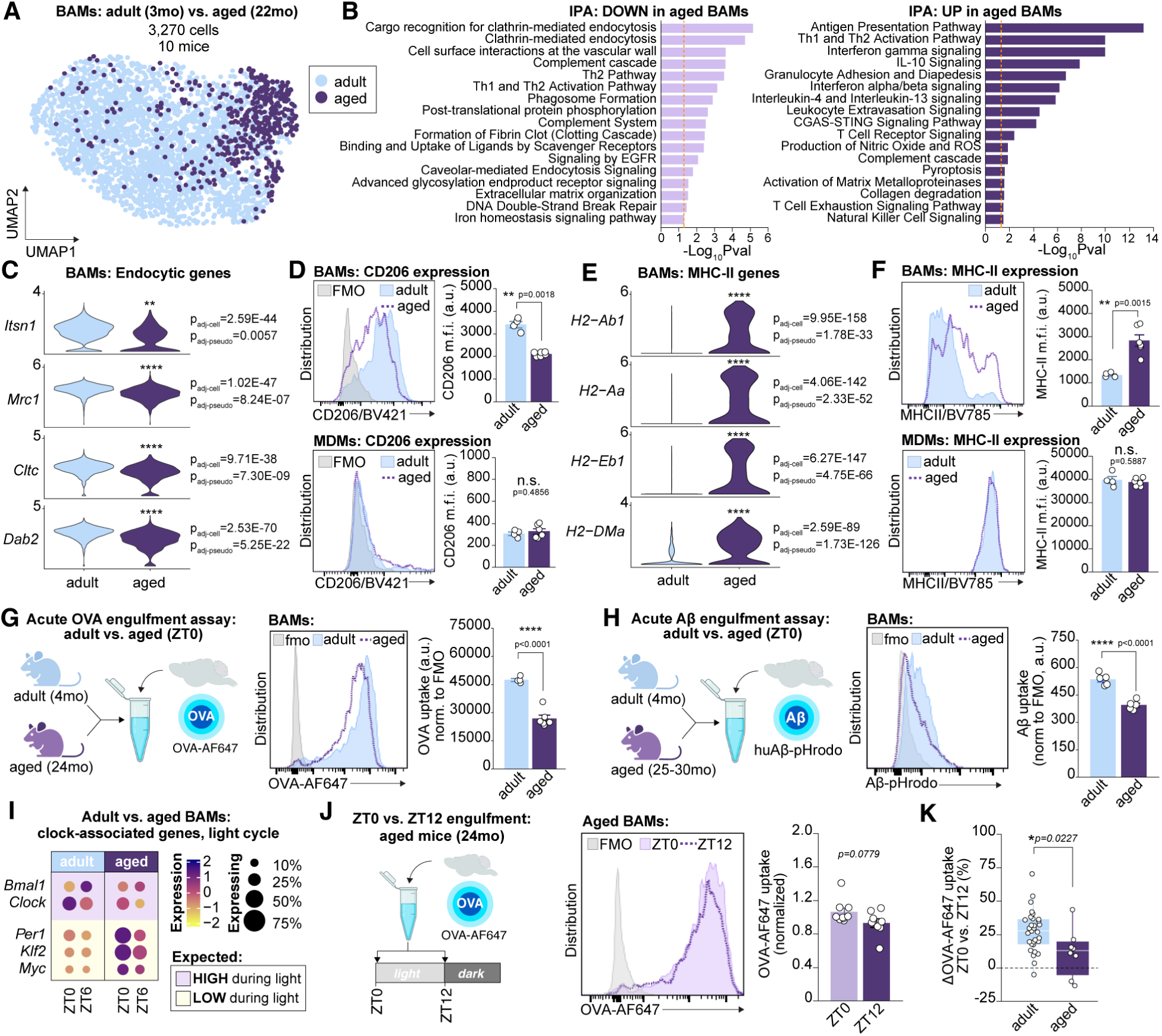
Aged BAMs exhibit perturbed clock gene expression and engulfment capacity (A) UMAP of BAMs from 6 adult and 4 aged mice (n of 3 adult and 2 aged per time point, time points = ZT0 and ZT6). All independent samples. (B) IPA of genes up- and down-regulated in adult vs. aged BAMs. DEGs were obtained from pseudobulk analysis. Selected pathways shown. Dotted line represents significance threshold. (C) CME-associated genes by age. Adjusted p-values shown for both cell-level and pseudobulk analyses; asterisks denote pseudobulk level significance. (D) Representative histograms and flow cytometric quantification of CD206 expression on BAMs and MDMs. CD206 is downregulated in aged BAMs (p=0.0018) but not aged MDMs (p=0.4856) relative to adult controls (two-tailed t-tests). N of 4 adult and 6 aged mice, all male B6N collected at ZT0. (E) MHCII-associated genes by age. Adjusted p-values shown for both cell-level and pseudobulk analyses; asterisks denote pseudobulk level significance. (F) Representative histograms and flow cytometric quantification of MHC-II expression on BAMs and MDMs. MHC-II is upregulated in aged BAMs (p=0.0015) but not aged MDMs (p=0.5587) relative to adult controls (two-tailed t-tests). N of 4 adult and 6 aged mice, all male B6N collected at ZT0. (**G and H**) Experimental design, representative histogram, and group-level quantification of OVA-AF647 or Aβ-pHrodo uptake in BAMs from adult versus aged mice. Both OVA-AF647 and Aβ uptake are reduced in aged BAMs (both p<0.0001; two-tailed t-tests). (I) Heatmap of clock and clock-associated genes at ZT0 and ZT6 in BAMs from adult vs. aged mice (n of 3 adult/time, n of 2 aged/time, independent samples). Plot background is pseudo-colored based on the theoretical pattern of clock gene expression based on time of day. (J) Experimental design, representative histogram, and group-level quantification of OVA-AF647 uptake at ZT0 versus ZT12 in aged BAMs. Results pooled from two independent experiments. Marginal difference in uptake across time (p=0.0779; mixed effects model accounting for experimental replicate; n of 9 mice per time point). (K) Percent change (Δ) in OVA-AF647 uptake at ZT12 relative to ZT0 in adult versus aged BAMs. Δ is reduced in aged BAMs (p=0.0227; Wilcoxon rank-sum). Only Δ from adult, but not aged, BAMs is significantly non-zero (p<0.0001 and p=0.1152, respectively; one-sample t-tests against hypothesized mean of 0). *For all panels: points represent individual mice, bars and error bars represent mean and SEM, box and whisker plots represent median and IQR. ****p<0.0001, ***p<0.001, **p<0.01, *p<0.05. All normalization to background using age-matched negative controls to account for age-related changes in cellular autofluorescence. Illustrations made with BioRender. See also **Figure S5, Table S5***.

Multiple genes belonging to the endocytic signature of adult BAMs during the light cycle were massively downregulated in aged BAMs, including *Itsn1, Mrc1, Cltc,* and *Dab2* (**Figure 5C**). Flow cytometric analysis confirmed that surface level CD206 expression was decreased by nearly 50% in aged versus adult BAMs (**Figure 5D**). In contrast, MDMs expressed a low but consistent amount of surface CD206 in the adult and aged brain. As previously reported^36,76,77^, aged BAMs instead exhibited a transcriptional signature vastly enriched for genes involved in class II antigen presentation, including *H2-Ab1, H2-Aa, H2-Eb1,* and *H2-DMa*, that was mirrored by nearly doubled surface MHC-II expression relative to adult BAMs (**Figure 5E-F**). Thus, transcriptional and flow cytometric analysis suggests that BAMs shift from a highly endocytic to an antigen-presenting state in aged mice.

To test whether downregulation of endocytic machinery was associated with decreased engulfment, we performed acute engulfment assays using both OVA-AF647 and pHrodo-conjugated Aβ (**Figure 5G-H**). In both cases, aged BAMs exhibited profoundly decreased substrate uptake compared to adult BAMs: OVA-AF647 engulfment was reduced by approximately 50%, and Aβ-pHrodo engulfment was reduced by approximately 30%. This was likely not attributable to differences in substrate diffusion during the engulfment assay across ages as both the proportion of BAMs positive for OVA-AF647 was unchanged between adult and aged mice, as was BAM yield (**Figure S5F,J**). There was a reduction in Aβ+ BAMs in aged mice relative to adults, though this was likely due to reduced signal-to-noise ratio due to the relatively dimmer Aβ-pHrodo signal combined with the increased autofluorescence of aged BAMs. MDMs also exhibited aging-related decreases in OVA-AF647 engulfment, and both aged MDMs and microglia showed reduced capacity for Aβ uptake relative to adult cells (**Figure S5J-M**). Thus, acute engulfment of fluid-borne substrates by macrophages decreases in aged mice. In BAMs, this is mirrored by downregulation of cellular machinery involved in CME, and specifically by downregulation of the mannose receptor CD206.

Aged BAMs also exhibited aberrant expression of clock-associated genes characterized by antiphase dominance during the light cycle (**Figure 5I**). Furthermore, aged BAMs upregulated known circadian disrupters such as *Myc,* a transcription factor that binds to E-box domains and competes with components of the circadian clock machinery such as BMAL1^78^. To test whether perturbed circadian gene expression has a functional consequence on aged BAMs, we performed a diurnal engulfment assay. Although there was still a trend toward reduced OVA-AF647 at ZT12 relative to ZT0 in aged mice, it was no longer statistically significant (**Figure 5J**). The dynamic range in engulfment between ZT0 and ZT12 was reduced in aged relative to adult BAMs (∼30% decrease in adult versus ∼10% decrease in aged BAMs; **Figure 5J**) and was only significantly non-zero in adult BAMs.These data indicate that aged BAMs exhibit perturbed circadian rhythms characterized by aberrant clock gene expression and impaired functional rhythmicity.

### BMAL1 deletion in BAMs worsens perivascular and leptomeningeal plaque burden

We next aimed to determine whether circadian perturbation of BAMs had a physiological consequence on the brain and leptomeninges. Because of our data indicating that BAM engulfment of fluid-borne substrates is under circadian regulation, we were specifically interested in how circadian dysregulation of BAMs may affect the accumulation of Aβ in the brain and its borders. To this end, we crossed the *Pf4^ΔBmal1^* line into the 5xFAD mouse model of AD (5x:*Pf4^ΔBmal1^*) and assessed amyloid plaque burden early in the development of pathology (P12). This time point was chosen as it coincides with initial plaque deposition in the cortex but precedes neuron loss by several months^53,79,80^, precluding effects of neurodegeneration on further circadian dysregulation and enabling anatomical segmentation of plaque localization (parenchymal versus perivascular). In the 5xFAD model, transgene expression is limited to neurons via the Thy1.2 promoter. We further verified that full-length human APP was undetectable in the bone marrow of 5xFAD mice to ensure that megakaryocytes (which undergo *Pf4*^Cre^-mediated recombination) are not a source of human APP/Aβ in this model (**Figure S6A**).

We assessed amyloid pathology across whole-brain sagittal sections using a tripartite labeling strategy: the Amylo-Glo dye to label fibrillized amyloid deposited in amyloid plaques (Biosensis), the 6E10 antibody to label human Aβ and APP, and the 1D1 antibody to label human amyloid precursor protein (APP; see **Methods; Figure S6B**). Congo red, a commonly used dye for amyloid plaque detection, shows non-specific binding to large perivascular structures in wild-type mice^81^, possibly to fibrillar collagen structures surrounding large vessels. To ensure that this was not the case for our pathological assessment, we also ensured that Amylo-Glo did not show non-specific binding in wild-type mice (**Figure S6C**). Three regions of interest were specifically interrogated: the leptomeninges (segmented dorsally), the hippocampal formation, and the cortex.

While there was no difference in brain parenchymal plaque burden, leptomeningeal plaque burden showed a striking increase in 5x:*Pf4^ΔBmal1^* mice relative to their 5x:*Bmal1^fl/fl^* littermate controls (**Figure 6A-C**). This result was highly consistent both within- and between-mice (**Figure 6C**). Increased leptomeningeal plaque burden in 5x:*Pf4^ΔBmal1^* mice held both when normalizing to cortical and hippocampal plaque burden (**Figure 6D,E**) and when Aβ was labeled with an antibody versus a dye (**Figure 6F**). We observed no difference in APP signal between 5x:*Pf4^ΔBmal1^* mice and their littermates (**Figure S6D,E**), suggesting that differences in plaque burden were driven by clearance of Aβ peptides themselves rather than via production of APP. Generally, leptomeningeal plaques were smaller than both cortical and hippocampal plaques, and plaque size was independent of genotype (**Figure S6F**). Instead, the difference in leptomeningeal pathology was driven by a substantial increase in plaque number in 5x:*Pf4^ΔBmal1^* mice (**Figure 6G**). This effect was dependent on conditional *Bmal1* deletion: the expression of *Pf4*^Cre^ alone in the 5xFAD line (5x:*Pf4*^Cre^) had no effect on leptomeningeal plaque burden, vascular plaque burden, nor APP signal (**Figure S6G**).

**Figure 6.**
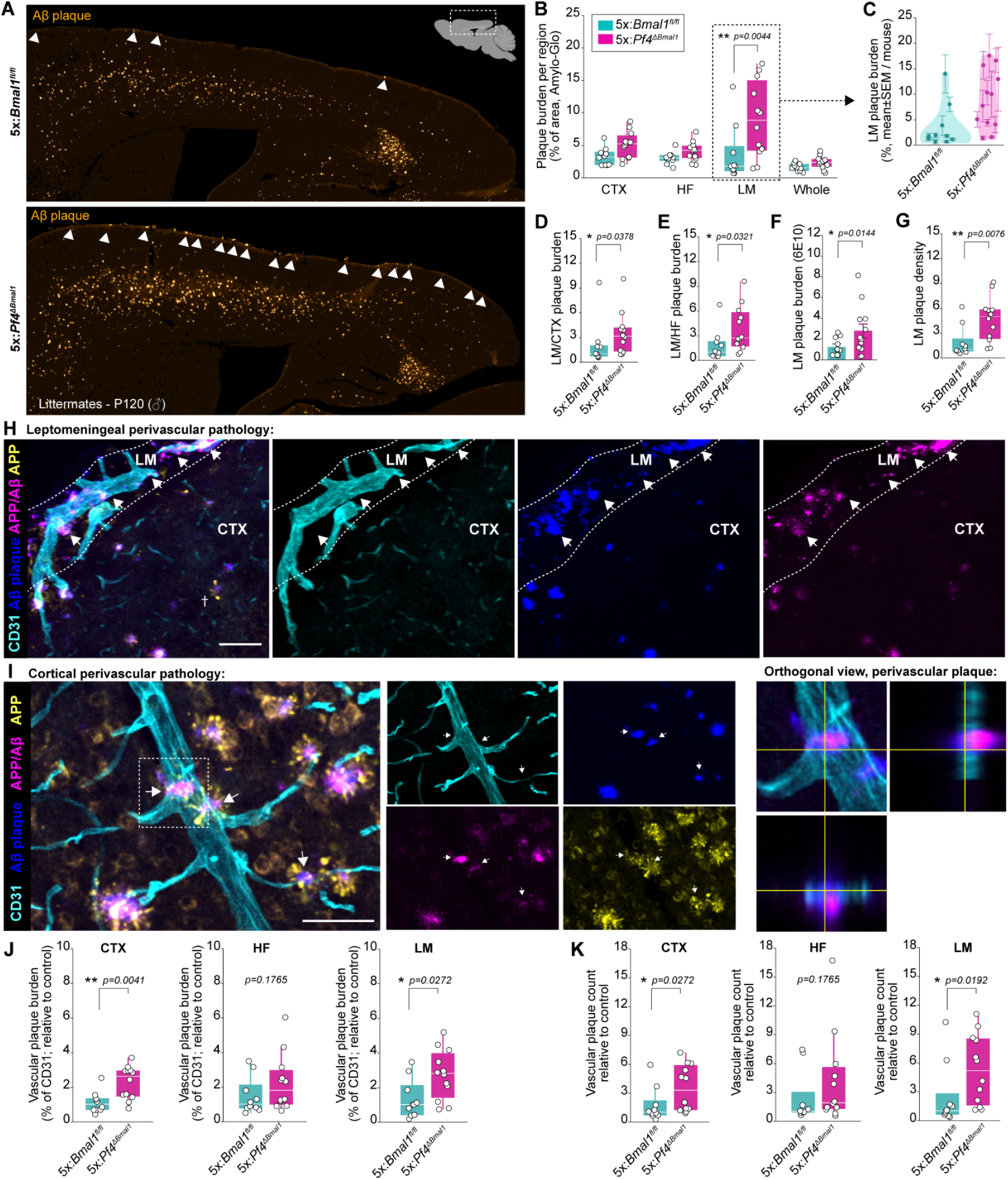
BMAL1 deletion in BAMs precipitates perivascular and leptomeningeal plaque burden in the 5xFAD model of Alzheimer’s Disease (A) Representative sagittal sections from male 5x:*Bmal1^fl/f^* and 5x:*Pf4^ΔBmal1^* littermates (P120). Aβ plaque is stained with Amylo-Glo, white arrowheads point to leptomeningeal plaque. (**B and C**) Group level-quantification of plaque burden per region in 5x:*Pf4^ΔBmal1^*mice versus their littermates (n of 10-12 per genotype). Plaque labeled with Amylo-Glo. CTX = cortex, HF = hippocampal formation, LM = leptomeninges. Region*genotype interaction effect: p=0.0108 (mixed effects model controlling for variance due to individual mouse and sex). Contour plot in (C) shows mean and SEM of LM plaque burden across sections per mouse. (**D and E**) Group-level quantification of LM plaque burden normalized to same-section CTX plaque and HF plaque burden. In both cases, relative plaque burden is increased in 5x:*Pf4^ΔBmal1^* versus their littermate controls. Comparisons using Mann-Whitney U test, n of 10-12 per genotype. (F) Group-level quantification of LM plaque burden stained with antibody (normalized to littermate controls; n of 10-12 per genotype). LM plaque burden is increased in 5x:*Pf4^ΔBmal1^* mice (mixed effects model controlling for variance due to sex). (G) LM plaque density (calculated as plaque number/area, normalized to controls) is increased in 5x:*Pf4^ΔBmal1^* versus their littermates. Comparisons using Mann-Whitney U test, n of 10-12 per genotype. (**H and I**) Representative confocal microscopy images of perivascular plaque along pial vessels (H) and a cortical venule (I). Perivascular plaques are indicated via white arrowheads. † Denotes axonal swelling adjacent to parenchymal plaques, characterized by heightened APP signal; these features are absent from leptomeningeal plaques. Scale bars = 50μm. (**J and K**) Group-level quantification of vascular plaque burden (% of CD31 signal co-localized with Aβ plaque; normalized to littermate controls) and vascular plaque count (number of plaques colocalizing with CD31 per area; normalized to littermate controls). Vascular plaque burden and count are increased in 5x:*Pf4^ΔBmal1^* mice versus their 5x:*Bmal1^fl/fl^* littermates in the CTX and LM, but not HF (n of 10-12 per genotype; comparisons using Mann-Whitney U test). *For all panels: points represent individual mice. Bars and error bars represent mean and SEM. Post-hoc testing with Tukey’s HSD. Box-and-whisker plots represent median and IQR (points unconnected to whiskers are considered outliers by IQR). ****p<0.0001, ***p<0.001, **p<0.01, *p<0.05. See also **Figure S6***.

Furthermore, we observed substantial plaque deposition along brain vasculature in 5x:*Pf4^ΔBmal1^* mice (**Figure 6H-I, Figure S6H**). These “perivascular plaques” resembled small spherical plaques situated on or around blood vessels, as previously described in models of capillary amyloidosis^82^. Perivascular plaques were detectable along large pial arteries, diving cortical arterioles, ascending cortical venules and pial veins, and smaller vessels within the cortex and hippocampus. Perivascular plaques were identified by partial colocalization with CD31, an endothelial cell marker, as well as by double-positivity for both Amylo-Glo and an anti-Aβ antibody to preclude non-specific signal (**Figure 6H-I**). Peri-arterial plaques were highly compacted, thus anti-Aβ antibody staining tended to localize to the plaque periphery (**Figure S6H**). Generally, the leptomeninges were far more enriched for perivascular plaque compared to the cortex and hippocampal formation; nearly half of all Aβ plaque signal directly colocalized with CD31 in the leptomeninges (**Figure S6I**). Within both the cortex and the leptomeninges, 5x:*Pf4^ΔBmal1^* mice exhibited increased vascular plaque burden (**Figure 6J**) driven by a heightened number of perivascular plaques (**Figure 6K**). Taken together, these results indicate that *Bmal1* loss associated with a loss of circadian BAM function precipitates brain border amyloidosis.

## DISCUSSION

Circadian rhythms entrain cellular physiology to time-of-day by synchronizing biological processes at the level of entire organs and organisms. Circadian rhythm disruption perturbs this synchrony and increases the risk of developing AD, yet the mechanisms underlying this risk are incompletely understood. By transcriptionally and functionally assessing the brain’s immune compartment around the day-night cycle, we identified BAMs as highly rhythmic cells. We found that BAMs are remarkably efficient at engulfing extracellular, fluid-borne substrates including Aβ and share features of highly endocytic cells. Importantly, we show that BAM engulfment is regulated by the circadian clock and that, in a mouse model of AD, loss of *Bmal1* in BAMs precipitates brain border plaque deposition. Our work also adds to a handful of studies linking BAM perturbations to amyloid pathology^31,32^, highlighting that not only BAM functions but also the time at which BAMs perform their functions critically influence brain homeostasis.

The exceptional scavenging capacity of BAMs has long been recognized, historically via studies of ink injection to the rat brain and more recently via injection of fluorescently conjugated substrates to the murine brain^31,83,84^. We show that this heightened capacity is in part mediated by the macrophage mannose receptor CD206. Open questions remain regarding the interaction between CD206 and Aβ. We used synthetic Aβ peptides and fibrils for our acute engulfment assays. While this approach removes substantial variability from our readouts, it is also reductionist given growing evidence that Aβ is not only glycosylated but also potentially differentially glycosylated in disease^85^. The extracellular domain of CD206 contains C-type carbohydrate recognition domains that bind to glycoproteins for their rapid uptake and clearance^69^. Whether different Aβ glycosylation patterns, as well as possible diurnal rhythmicity in those patterns, affect the ability of BAMs to engulf Aβ is an unanswered but interesting question.

Microglial phagocytosis has been a primary focus of research on Aβ clearance in the brain. Our study highlights that endocytosis, and specifically scavenger receptor-mediated CME by brain border-resident macrophages, may also contribute to the clearance of fluid-borne Aβ. We find that expression of the transcriptional machinery involved in CME (genes including *Dab2, Itsn1, Cltc, Mrc1*) as well as protein-level expression of CD206 (which is internalized via CME), both deteriorate with aging. Given the many AD risk genes associated with endocytosis^86–91^ (e.g. *PICALM, SORL1, CD2AP, BIN1*), as well as previous reports of age-related decline in CME-associated genetic signatures in the human brain borders^92^, we believe further interrogation of the role of endocytosis by brain border immune cells will deepen our understanding of AD pathogenesis with the potential of uncovering new therapeutic targets.

In our study, BAMs exhibited coordinated upregulation of phagocytic genes during the active phase and endocytic genes during the rest phase. Rhythmicity in macrophage phagocytosis has been previously reported^26^, but comparatively little is known of rhythmicity in macrophage endocytosis. In line with our results, large-scale ChIP-seq profiling of the liver shows that ubiquitously-expressed endocytic genes peak during the rest phase, and in many cases are direct transcriptional targets of BMAL1^68^. Thus, heightened endocytic capacity is likely a conserved feature of the rest phase across cell types and organs. Because BAMs far exceeded microglia and MDMs in scavenging capacity, our ability to sensitively detect diurnal changes was likely increased within BAMs. It is possible that microglia endocytosis is also rhythmic, but that we were not powered to detect it in acute assays. It is also probable that heightened endocytic uptake of extracellular solutes is more physiologically relevant for BAMs. Microglia are highly ramified cells dwelling in the brain parenchyma that can contact and phagocytose surrounding substrates. In contrast, BAMs must instead sample draining fluid-borne solutes exiting the brain. Phagocytosis is a time- and energy-consuming process requiring extensive membrane re-arrangement. The rapid process of CME by BAMs may enable scavenging of fluid-borne substrates that are only present for a brief period of time in brain border regions.

Why might phagocytosis and endocytosis peak at different points in the circadian cycle? One possibility is that the segregation of tissue defense and repair would allow innate immune cells to anticipate the time of highest pathogen exposure risk (i.e., during waking behavior) rather than simply respond to it. Similarly, heightened endocytosis during the rest phase may prime tissue-resident macrophages for the clearance of metabolites and debris. Converging evidence from multiple research groups and model systems indicates that Aβ and tau accumulate in brain ISF during wakefulness and peak in the CSF at the onset of the rest phase^17,46,47^, and that fluid egress to the brain borders is increased during sleep^48–52^. BAMs live along routes of fluid egress from the brain; thus, circadian anticipation of solute drainage may be critical for enforcing efficient BAM engulfment of substrates that are differentially abundant across the day-night cycle. This could represent one mechanism to explain the increased risk of dementia and AD associated with circadian misalignment, such as that enforced by night-shift work^11,12^.

In summary, we generated a single-cell atlas of transcriptional changes in both parenchymal and brain border immune niches around the day-night cycle. This atlas and our subsequent functional experiments highlighted the remarkable scavenging capacity of BAMs and its dynamic range around the clock. We propose that circadian perturbations, for example due to aging or night-shift work, may disrupt CNS homeostasis in part by impairing the timing of critical brain border innate immune functions, including the clearance of pathogenic proteins.

### Limitations of this study

While we characterize one clock-controlled BAM function – engulfment – *Bmal1* deletion in BAMs likely worsens amyloid pathology by dysregulating the timing of many cellular functions beyond engulfment, such as ECM maintenance^36^. Further studies, for example deleting E-box domains from genes of interest, would be needed to assess the specific contributions of rhythmicity in a single cellular pathway to AD pathogenesis. However, this is less translationally relevant, as circadian perturbations involve global rather than gene-specific disruption. Additionally, we are limited by the tools available to genetically manipulate BAMs. Existing Cre lines can be used to target brain myeloid cells broadly (*Cx3cr1-Cre, Cx3cr1-CreER*), but due to the specific differences between BAMs and microglia within the scope of this work, we opted for an approach with minimal microglial recombination: the *Pf4*^Cre^ line. The *Pf4*^Cre^ line was originally developed to target megakaryocytes but recombines in BAMs due to their high level of CXCL4 expression. While we can and do control for the effects of the BAC transgene, we cannot exclude the possibility that *Bmal1* deletion in megakaryocytes may contribute to the phenotype observed in the 5x:*Pf4^ΔBmal1^* mice. Megakaryocytes are located in the bone marrow and do not contribute to human APP overexpression in the 5xFAD model of AD: transgene expression is driven by the Thy1.2 promoter and limited to subsets of neurons. We confirmed that full-length human APP is absent from the bone marrow of 5xFAD mice. We also verified that APP expression in the brain is unchanged in 5x:*Pf4^ΔBmal1^* mice versus their 5x:*Bmal1^fl/fl^*littermate controls, suggesting that it is not the production of APP but rather the clearance of cleaved Aβ that is impacted in our model. Complementary approaches in other mouse lines, such as a *Cd206-CreER* model^33,93^, would also be informative (albeit with other off-target effects in distal cell types such as liver sinusoidal and lymphatic endothelial cells). Finally, the work included in this study was performed in mice. Mouse models of amyloidosis do not fully recapitulate the rich complexity of AD, but they allow us to specifically assess how genetic manipulations affect anatomical patterns of Aβ plaque deposition in an overproduction model. In addition, although the cellular mechanisms we study are highly conserved across mammalian species (CD206, BMAL1), large open questions remain in the field regarding how the circadian clock differentially regulates cellular physiology in diurnal versus nocturnal species. As technological advances eventually enable experimentation in rare human brain cell types, the circadian regulation of human BAMs would be an interesting question to explore.

## MATERIALS AND METHODS

### Mice

All experimental procedures were carried out following protocols approved by the Institutional Animal Care and Use Committee (IACUC) of Boston Children’s Hospital or the Broad Institute, in accordance with guidelines established by the NIH for the humane use and treatment of laboratory animals. All mice were group-housed except in cases of cage-mate aggression; both females and males were used in experiments. Mice were housed in Optimice rack-mounted cages (Animal Care Systems #C79112PF; Boston Children’s Hospital) or Sealsafe Plus cages (Techniplast #GM500; Broad Institute) following temperature, capacity, and environmental recommendations from the American Association for Accreditation of Laboratory Animal Care (AALAC). All mice had *ad libitum* access to food and water and were kept in standard 12-hour-light, 12-hour-dark cycles in either a regular or reverse light-cycle animal housing housing rooms. For experiments performed in circadian time, mice were kept in temperature- and humidity-monitored circadian cabinets (Actimetrics) in the Animal Behavior & Physiology Core of Boston Children’s Hospital for up to 12 hours following the expected onset of light exposure. Mice transferred to reverse light cycle rooms were given a minimum of 2 weeks to habituate prior to experimentation.

Transgenic mice were obtained from the Jackson Laboratory and bred in-house on a C57BL/6J background. The *Pf4*^Cre^ line^65^ (Jax #008535) was used for BAM-targeted crosses. This strain was crossed to either the Ai14 tdTomato line^94^ (Jax #007914) or the ROSA^mT/mG^ line^95^ (Jax #007676) for fluorescent reporter expression, or to the Bmal1^lox^ line^66^ (Jax #007668) for BMAL1 deletion. For all *Pf4*^Cre^ crosses, only male carriers were used for breeding. The CD206-knockout line^70^ was obtained out of cryorecovery from the Jackson Laboratory and maintained on a C57BL/6 background (Jax #007620). For large circadian experiments in wild-type mice, aged-matched cohorts of C57BL/6 mice were obtained from the Jackson Laboratory (C57BL/6J, Jax #000664) and kept in the animal facility for a minimum of two weeks prior to experimentation to ensure environmental habituation. Aged C57BL/6 wild-type mice were obtained from the National Institute of Aging (NIA) and adult controls were obtained from either Charles River (C57BL/6NCrl, Strain #027) or the Jackson Laboratory (C57BL/6NJ Jax #005304). The 5xFAD strain^55^ was used for modeling amyloidosis. 5xFAD mice on a C57BL/6J background were obtained from Mutant Mouse Resource & Research Centers (MMRRC; #034848-JAX) and bred in-house, maintaining the C57BL/6J background. The 5xFAD line was subsequently crossed to the *Pf4*^Cre^ and Bmal1^lox^ lines for BAM-targeted BMAL1 deletion. For all 5xFAD crosses and maintenance, only male carriers were used and no homozygotes were generated to prevent effects of paternal versus maternal inheritance on amyloid pathology^53^. Transgenic mice were genotyped using tissue samples sent to Transnetyx.

### Sample preparation for flow cytometric processing

#### Brain preparation for flow cytometric analysis

Mice were deeply anesthetized via intraperitoneal injection of Avertin (2,2,2-Tribromoethanl in 2-Methyl-2-butanol; Millipore Sigma #T48402 and #240486) diluted in Hank’s Balanced Salt Solution (HBSS, ThermoFisher Scientific #14175145). The plane of anesthesia was verified by loss of withdrawal reflex to the paw pinch test and loss of palpebral reflex to the eye-blink test. Then, mice were transcardially perfused with ice-cold HBSS to a total volume of ≥1mL per gram of body weight. Following decapitation, the skull cap was carefully removed, ensuring the dural meninges remained attached to the skull cap rather than the surface of the brain. The resulting brain sample (with leptomeninges and choroid plexus included) was placed in ice cold RPMI-1640 (no phenol red, ThermoFisher Scientific #11835030) containing 10mM HEPES (Sigma-Aldrich #H3537-100, “RPMI-H”). Brains were minced to a fine paste on 100mM petri dishes in 200uL of cold RPMI-H using a razor blade then transferred to 5mL capped FACS tubes (Corning #352058) and topped with an additional 2mL of RPMI-H. Then, samples were centrifuged for 5 minutes x 500g at 4C, decanted, and tissue pellets were re-suspended in a pre-heated digestion buffer containing Collagenase P (0.5 mg/mL, Sigma #11213865001), Dispase II (0.8 mg/mL, Worthington #LS02104), and DNAse-1 (250 U/mL, Worthington #LK003172) in RPMI-H. Whole brains were digested in 4mL of buffer and half brains in 2mL. After ensuring samples were thoroughly resuspended in the digest buffer, we placed tubes on a light-protected orbital rocker at 37C for gentle agitation throughout the 30-minute enzymatic digestion. Following digestion, samples were centrifuged for 5 minutes x 500g at 4C, the supernatant was discarded, and tissue was resuspended in 1mL of ice-cold FACS buffer composed of 0.5% Bovine Serum Albumin (Sigma-Aldrich #A2153) and 2mM EDTA (Research Products International #E14000) in HBSS. Brains were then triturated using a P1000 pipette tip, while carefully avoiding bubble formation, and passed through a 70µm filter into a 15mL tube. The resulting sample was subsequently diluted in 5mL of 25% BSA in HBSS and centrifuged for 10 minutes x 1200g at 4C for cellular enrichment and debris removal. The myelin layer and supernatant were carefully removed using a vacuum and subsequently a P1000 pipette tip, and the resulting cell pellet was re-suspended in FACS buffer and transferred to a 2mL tube for washing and staining of the resulting single-cell suspension.

#### Brain preparation for FACS prior to bulk or single-cell RNA sequencing

For FACS prior to sequencing, brains were prepared as above with the addition of transcriptional and translational inhibitors to several steps, an approach that has been shown to prevent *ex vivo* cellular activation signatures^37^ and in our case additionally ensured transcriptional profiles accurately reflected sample collection time. To this end, mice were transcardially perfused with a buffer containing Actinomycin D (5µg/mL, Sigma-Aldrich #A1410) and Triptolide (10µM, Sigma-Aldrich #T3652) in RPMI-H. Then, samples were incubated and minced in a solution containing 5µg/mL Actinomycin D, 10µM, Triptolide, and Anisomycin (27.1µg/mL, Sigma-Aldrich #A9789) in RMPI-H. For the diurnal scRNA-seq atlas and aged mouse scRNA-seq dataset, sample digestion was performed with the addition of the inhibitors and a cell-permeant DNA dye to label living cells (Vybrant DyeCycle Ruby Stain, Invitrogen #V10309), for a final digest buffer containing 0.5 mg/mL Collagenase P, 0.8 mg/mL Dispase II, 250U/mL DNAse-1, 5µg/mL Actinomycin D, 10µM Triptolide, 27.1µg/mL Anisomycin, and 5µM DyeCycle Ruby in RPMI-H. For the bulk RNAseq of purified BAMs, no DyeCycle Ruby was included in the digest. Samples were digested for 30 minutes at 37C, and subsequently underwent trituration, BSA enrichment, and generation of a single-cell suspension as described above. All reagents were kept light-protected throughout the preparation.

#### Acute labeling of circulating immune cells prior to perfusion

Following verification of plane of anesthesia and prior to cardiac perfusion, some mice received intravenous delivery of a conjugated antibody for labeling circulating immune cells (anti-CD45-PE, clone 30-F11, Biolegend #103106, 0.2µg/g), either via retro-orbital or intracardiac injection. The antibody was allowed to circulate for only three minutes prior to the beginning of transcardial perfusion to ensure cells were labeled while preventing antibody leakage into the CNS. For scRNA-seq datasets, 200uL of blood was acutely collected from the right atrium of each mouse to verify efficacy of the intracardiac (IC) anti-CD45 antibody delivery. Blood was immediately diluted into 10mL of FACS buffer on ice. Then, blood samples underwent two rounds of red blood cell lysis for 5 minutes each using ACK Lysing Buffer (ThermoFisher Scientific #A1049201) to obtain single-cell suspensions of leukocytes.

#### Antibody labeling of single-cell suspensions

Single-cell suspensions were centrifuged for 5 minutes x 500g at 4C, decanted, re-suspended in 200uL of FACS buffer, and transferred to 96-well U-bottom plates (Thermo Scientific #262162). Following another 5 minutes x 500g spin at 4C, samples were resuspended in 50µL of an Fc-blocking and viability dye solution composed of 2% Fc-blocking antibody (anti-mouse CD16/CD32 clone 2.4G2, BD Biosciences #553141) and 0.1% fixable viability dye (eBioscience Fixable Viability Dye eFluor 780, ThermoFisher Scientific #65-0865-18) in FACS buffer, and incubated for 10 minutes on ice. Then, 50µL of antibody cocktail solution containing 20% Brilliant Stain Buffer (BD #566385) as well as the desired antibody combination was added into each sample, carefully pipette-mixed, and incubated for 20 more minutes on ice. All antibody clones, catalog numbers, and dilutions are listed in **Table S6**. Following staining, samples were washed twice in FACS buffer and either directly underwent flow cytometric analysis or a 20-minute fixation in 100µL of intracellular (IC) fixation buffer (eBioscience #00-8222-49) at room temperature, followed by two more washes in FACS buffer. Fixed cells were stored at 4C in FACS buffer until flow cytometric analysis.

### Flow cytometric analysis and sorting

#### Flow cytometric analysis

Flow cytometric analysis was performed using a FACSSymphony S6 flow cytometer with FACSDIVA v9.5, a FACSAria SORP II flow cytometer with FACSDIVA v8.0.1, a FACSCanto II SORP with FACSDIVA v8.0.1, or a Beckman Coulter Cytoflex LX with CytExpert 2.6 software. For the FACSSymphony, FACSAria, and FACSCanto, single-cell suspensions in FACS buffer were filtered into FACS tubes through a 35µm filter cap and loaded into the machines for analysis. For the Beckman Coulter Cytoflex LX, single-cell suspensions in FACS buffer were directly loaded into the machine using a plate-reader attachment. For a subset of experiments aiming to obtain absolute cell counts, counting beads (CountBright Absolute Counting Beads, Thermo Fisher Scientific #C36950) were added to samples prior to acquisition. All samples within individual experiments were run together in a single flow cytometry session. Following acquisition, exported FSC files were analyzed using FlowJo v10.10.0.

#### Fluorescence-activated cell sorting (FACS)

For generation of scRNA-seq datasets as well as for BAM purification for bulk RNA sequencing, cells were sorted using a FACSSymphony S6 flow cytometer with FACSDIVA v9.5 equipped with a 100µm nozzle. A srting buffer composed of HBSS with 0.5% BSA was used instead of FACS buffer for all sorting steps. For the diurnal and aged scRNA-seq datasets, single-cell suspensions were transferred to pre-coated FACS tubes (Corning #352235) via a 35µm filter cap and incubated with DAPI (0.2µg/mL, BioLegend #422801) for five minutes prior to loading on the FACSSymphony. Viable singlets were identified based on DAPI^neg^, DyeCycle Ruby^pos^, and morphological features, and any contaminating peripheral immune cells remaining after perfusion were excluded as IC-CD45-PE^pos^. Finally, cells were gated for sorting using a combination of anti-CD45 and anti-CX3CR1 antibodies (**Figure S1B, Table S6**). For bulk RNA sequencing of purified BAMs, single-cell suspensions were transferred to pre-coated FACS tubes (Corning #352235) via a 35µm filter cap and loaded on the FACSSymphony. Viable singlets were identified based on morphological features and Viability Dye^neg^ (Invitrogen #65-0865-18), any remaining contaminating peripheral immune cells were excluded as IC-CD45-PE^pos^, and BAMs were identified using a combination of anti-CD45, anti-GR-1, anti-CD64, anti-CD11b, anti-CX3CR1, and anti-CD38 antibodies (**Figure S3J, Table S6**). All FACS samples were sorted into 2mL low-bind microcentrifuge tubes pre-filled with sorting buffer.

### Flow cytometric engulfment assays

#### Assessment of intrinsic engulfment capacity via acute ex vivo engulfment assays

For assessment of intrinsic engulfment capacity, we designed acute *ex vivo* engulfment assays enabling normalization of substrate concentration and exposure duration across samples. Following cardiac perfusion as described above, brains and attached leptomeninges were carefully dissected to ensure the dura mater was not adhered then minced to a fine paste in 200uL of ice cold RPMI-H using a razor blade. For circadian experiments, the time of sample dissection/collection was used for categorization (e.g. ZT0 vs. ZT12) to standardize across preparations. Tissue was pelleted via a 500g spin at 4C for 5 minutes then resuspended in a pre-warmed digest buffer containing Collagenase P (0.5 mg/mL, Sigma Aldrich #11213865001), Dispase II (0.8 mg/mL, ThermoFisher Scientific #17105041), and DNAse-1 (250 U/mL, Worthington #LS002007) with the addition of an exogenous substrate to engulf. Engulfed substrates included AF647-conjugated ovalbumin (OVA-AF647, 0.01mg/mL; Invitrogen #O34784), AF488-conjugated ovalbumin (OVA-AF488, 0.01mg/mL; Invitrogen #O34781), AF594-conjugated ovalbumin (OVA-AF594, 0.01mg/mL; Invitrogen #O34783), AF488-conjugated dextran (dextran-AF488, 10,000MW, 0.01mg/mL, Invitrogen #D22910), AF488-conjugated E. coli bioparticles (K1-12 strain, 0.01mg/mL, Invitrogen #E13231), and human synthetic Aβ_1-42_ pre-formed fibrils (1µM, StressMarq Biosciences #SPR-487E). Aβ_1-42_ fibrils were conjugated with pHrodo Deep Red (Invitrogen #P35358) following manufacturer instructions. We performed prior experiments using AF647-conjugated or unconjugated Aβ_1-42_ (0.01mg/mL; Anaspec #AS-64161 and AS-20276, respectively), however, the majority of Aβ signal in those experiments was from peptides adhered to the cell surface rather than internalized. Thus, we only used pHrodo-conjugated Aβ for acute engulfment assays. For whole brain engulfment assays, 4mL of digest buffer was used. For assays on half brains, 2mL was used. We did not further divide brains beyond hemispheres to reduce variability introduced by uneven starting tissue amounts. Once re-suspended in the digest engulfment buffer, samples were then light-protected and digested for 30 minutes at 37C under continuous gentle rotation from an orbital rocker. Immediately after the 30 minutes, samples were spun down for 5 minutes at 4C and the supernatant containing residual engulfment substrate was discarded. Samples were then processed for flow cytometry as described above, with the specification that samples engulfing fixable substrates were always fixed prior to flow cytometric analysis. Samples engulfing pHrodo-conjugated substrates were kept on ice and run live. For circadian engulfment assays, all samples from individual experiments were run on the cytometer together. Thus, for pHrodo-conjugated substrates, experiments could only be performed using reverse light cycle rooms. Engulfment assays were only performed during the digest rather than after obtention of a single-cell suspension to ensure the highest consistency in starting cell/tissue amount.

#### Assessment of in vivo Aβ engulfment using FEAST

For assessment of *in vivo* Aβ engulfment in the 5xFAD model of AD, we used the fixed-cell FEAST protocol^54^ as live cell preparations are highly susceptible to artifactual *ex vivo* engulfment by BAMs. Briefly, mice were transcardially perfused using 10mL of HBSS followed by 20mL of fixation buffer (BioLegend #420801). All perfusion materials were kept ice cold throughout the procedure. Brains were dissected and incubated in a PFA quenching solution composed of HEPES (5mM, Sigma-Aldrich #7365-45-9), Tris (250mM, Millipore Sigma #GE17-1321-01), and Glycine (250mM, Millipore Sigma #G7126) in HBSS. The cortex and corpus callosum were microdissected using forceps and chopped to a fine paste in 100uL of the quenching solution using a razor blade. Samples were then spun for 3 minutes at 300g and the supernatant was discarded prior to resuspension in a digest buffer composed of 800U/mL of Collagenase IV (Worthington #LS004189) in RPMI-H. Samples were digested for 2 hours at 37C with gentle agitation on an orbital rocker. We found through prior experiments that micro-dissection of the cortex and corpus callosum in addition to assessment at an early stage of amyloid pathology were necessary to appropriately yield BAMs from the 5xFAD brain using the fixed cell FEAST protocol. Using too much starting tissue or too late of a pathological stage led to the formation of large Aβ plaques aggregates during the 2 hour digest that vastly reduced yield. After digestion, samples underwent the trituration, BSA enrichment, acid wash, and blocking steps as described in the fixed cell FEAST protocol to obtain a single-cell suspension free of contaminating substrates adhered to the cell surface^54^. Finally, cells underwent a second fixation in IC fixation buffer (eBioscience #00-8222-49). Following fixation, cells were permeabilized in a 1X dilution of permeabilization buffer (eBioscience #00-8333-56) and incubated for 10 minutes with 2% Fc-blocking antibody (anti-CD16/CD32, BD Bioscience) before staining with a panel of primary-conjugated antibodies composed of anti-CD45 (PE-Cy5.5, eBioscience), anti-GR1 (AF488, BioLegend), anti-CD68 (APC-Cy7, BioLegend), anti-CX3CR1 (BV711, BioLegend), anti-CD38 (PE-Cy7, BioLegend), anti-MHCII (BV785, BioLegend), and anti-Aβ_1-16_ (clone 6E10, AF647, BioLegend), as well as 1:20,000 DAPI (BioLegend #422801). All antibody clones, catalog numbers, and dilutions are listed in **Table S6**. Wild-type littermates were used as a negative control for intracellular Aβ staining. Following 20 minutes of staining, samples were washed twice in 200uL of permeabilization buffer, then once in 200uL of FACS buffer prior to resuspension in FACS buffer and transfer to FACS tubes through a 35µM filter cap (Corning #352235). All FEAST samples were analyzed using a FACSSymphony S6 flow cytometer with FACSDIVA software (v9.5). Exported FCS files were analyzed using FlowJo v10.10.0.

#### Single-cell RNA sequencing

#### Single Cell Partitioning & Library Generation

Following FACS, cells were centrifuged 500 x g for 5 minutes at 4C in PCR strip tubes. The supernatant was then removed until ∼30μl remained. The cell pellet was resuspended first by gentle pulse vortex and then gentle pipetting (x15). FACS buffer was then added to the cell suspension to a final volume of 38.7μl and the entire volume was carried forward for droplet generation. scRNA-seq experiments were performed using 10X Genomics 5’ Single-cell Version 2 and NextGEM Chromium Controller. Following droplet generation, barcoded libraries were generated following manufacturer’s specifications with the following experiment specifics steps: cDNA amplification 14 cycles, sample index PCR 14 cycles.

#### Next Generation Sequencing

Single-cell libraries were first shallow-depth sequenced using the Illumina NextSeq 500 with a High Output 150 cycle flowcell. All libraries were pooled at equimolar concentration following dilution and then denatured/diluted according to Illumina denature and dilution guidelines for NextSeq500 High Output flow cells. Reads were as follows according to Read 1: 26bp (16bp cell barcode, 10bp UMI); Index 1: 10bp (Illumina i7 sample index); Index 2: 10bp (Illumina i5 sample index); Read 2: 90bp (Transcript insert). For full depth sequencing, libraries were re-pooled based on cell number determined from shallow depth sequencing. Pooling was performed to create equal read depth per cell across all libraries. The library pool was then sequenced on a NovaseqX Plus 10B flowcell at the Broad Institute’s Genomics Platform. Reads were as follows according to Read 1: 26bp (16bp cell barcode, 10bp UMI); Index 1: 10bp (Illumina i7 sample index); Index 2: 10bp (Illumina i5 sample index); Read 2: 90bp (Transcript insert).

### Single-cell data processing and analysis

#### Single-cell Data Preprocessing

All Cell Ranger preprocessing of sequencing data was performed on Harvard Medical School’s O2 High Performance Compute Cluster. CellBender preprocessing was performed on Terra cloud environment using google cloud platform GPUs.

For shallow depth sequencing, raw Illumina bcl files were demultiplexed using Cell Ranger version 7.1.0 and bcl2fastq version 2.20.0.422 using ‘mkfastq’ step using default specifications. For full depth sequencing, raw Illumina bcl files were demultiplexed using bcl2fastq version 2.20.0.422 using specifications equivalent to Cell Ranger ‘mkfastq’ wrapper. Individual sample gene expression matrices were generated using Cell Ranger version 7.1.0 ‘count’ step using default mm10 genome supplied by 10X Genomics Cell Ranger 7.1.0 (reference annotation corresponds to filtered version of Ensembl v98; see 10X support website for further information). The only non-default settings used for Cell Ranger ‘count’ were setting ‘--include-introns’ flag to false, to exclude reads mapping to introns from quantification.

Following count matrix generation, the samples were processed using CellBender^96^ to remove ambient RNA and bulk PCR artifacts in unsupervised fashion. CellBender was run on Google Cloud GPUs through Terra platform using ‘remove-background’ workflow (Snapshot 13: corresponds to v0.3.0). The following settings were used for CellBender processing: ‘expected_cells’ was set equal to Cell Ranger called cells, ‘total_droplets_included’ set based on examination of barcode rank plots (via Cell Ranger output and DropletUtils/scCustomize^97,98^ ‘barcodeRanks’ and ‘Iterate_Barcode_Rank_Plot’), ‘fpr = 0.01’, ‘epochs=300’. CellBender corrected counts matrix containing all droplets was loaded in R via scCustomize along with filtered CellBender and Cell Ranger matrices. A union of all droplet barcodes called as cells between Cell Ranger and CellBender was used to filter the full CellBender corrected matrix for further QC and analysis.

#### Quality control

All further processing and analysis was conducted using Seurat v5^99,100^. The dataset was initially filtered to only include cells with > 500 detected genes, as well as > 0% and < 10% of transcripts mapping to mitochondrial genes. The filtered object was then clustered following the standard Seurat v5 workflow (normalization, variable feature identification, data scaling, linear dimension reduction, determination of dimensionality, and clustering). The output was visualized using non-linear dimension reduction via uniform manifold approximation and projection (UMAP). Broad immune and non-immune cell types were identified based on established markers and subsetted for further quality control (QC).

To identify doublets, stripped nuclei, artifacts, and low-quality cells, subsetted cell types were processed via sequential high-resolution clustering^101^. Outlier clusters were identified based on 1) lack of ribosomal genes (e.g. *Rpl29,* likely stripped nuclei), 2) aberrantly high feature count and RNA count in conjunction with incongruent cell type-specific gene expression (evident multiplets, e.g. co-expression of *Ms4a7* and *Atp13a5* for a macrophage-pericyte doublet), or 3) aberrantly high mitochondrial gene expression in conjunction with low gene number (low quality cells). Outliers were removed and high-resolution clustering was repeated for each cell type until all low-quality cells had been identified and excluded. This biology- and evidence-based approach ensured that every cell in the dataset had been examined for multiple QC metrics. All cells that passed this QC pipeline were subsetted from the initial filtered object to create the final Seurat object.

#### Integration of adult and aged mouse datasets

All single-cell samples included in this study were acquired in one experiment (both the diurnal scRNA-seq atlas and the aged brain sequencing). Aged wild-type mice were obtained from the National Institute of Aging and were on a slightly different genetic background (C57BL/6N) than adult mice used in the circadian single-cell atlas (C57BL/6J). Thus, although samples from both age groups were acquired and processed for scRNA-seq concurrently, we opted to perform QC separately for these two datasets to preclude any biologically relevant differences between adult and aged brains affecting our QC pipeline. No computational integration was needed for examining the merged datasets after QC; all cell types clustered together across adult and aged samples and no batch effect was present given the concurrent acquisition and sequencing. All protein-level and functional validation experiments were performed with adult C57BL/6N as control counterparts to aged C57BL/6N mice.

#### Atlas annotation and data visualization

The final Seurat object was processed using the standard Seurat v5 workflow^99,100^. Two rounds of annotation were performed: broad versus fine. For broad cell annotation, cluster markers were obtained for the whole atlas. For fine annotation, cells were divided into “immune” and “non-immune” categories for further clustering and annotation. In both cases, the FindAllMarkers function from Seurat v5 was used with a minimum expression of 25% and a LogFC of 0.405. Genes, clusters, and metadata parameters were visualized using UMAPs as well as dimensional reduction plots (DimPlots), feature plots, heat maps, violin plots, and dot plots using scCustomize (version 3.0.1)^98^. Volcano plots were made using EnhancedVolcano (version 1.24.0)^102^.

#### Differential gene expression

DEG analysis was performed using both cell-level and pseudobulk approaches; the employed approach is indicated in each figure legend. For comparison of absolute number of DEGs between ZT0 and ZT12 across immune cell types, we only included immune populations with at least 250 cells per time point. ILCs, pDCs, and MC/BPs were not included due to scarcity. Data were aggregated per biological replicate using the AggregateExpression function from Seurat v5, then samples from ZT0 and ZT12 were compared using DESeq2 with an adjusted p-value cutoff of 0.05 and a log_2_FC cutoff of 0.2. This lower FC cutoff was used because, unlike core circadian clock genes that exhibit large diurnal shifts in expression, the oscillations in many clock-controlled genes tend to have smaller amplitudes. Given that each time point only had an n of 3 biological replicates, the smallest log_2_FC we were powered to detect had an absolute value of 0.24, corresponding to approximately a 20% change in expression (**Table S2**). We also performed cell-level comparison, matching cell number per cell type by random subsetting of 500 cells, for immune cells between ZT0 and ZT12. This approach was just used to verify that the pattern of differential gene expression was comparable across analysis methods and was not used for biological interpretation. For this, we used FindMarkers with default Wilcoxon Rank sum, min.pct set to 0.05, and applying the same adjusted p-value cutoff of 0.05 and a log_2_FC cutoff of 0.2 as for the pseudobulk analysis. Both analyses identified the same trend of higher DEG count in BAMs.

For comparison of gene expression across time in BAMs, two approaches were used. First, we used the output from the pseudobulk analysis described above. Then, we leveraged the power of the single-cell resolution in our dataset to interrogate smaller magnitude changes occurring around the clock in BAMs. For this, we used MAST^67^, a gene enrichment analysis approach using a generalized linear framework that controls for gene detection rate per cell and provides enhanced rigor relative to alternative options. We employed MAST with an adjusted p-value cutoff of 0.05, a log_2_FC cutoff of 0.15 (corresponding to approximately a 10% change in expression) and a min.pct cutoff of 0.05 across all time points. Then, we further subsetted results to only include genes that 1) were considered differentially expressed at a minimum of two time points, and 2) had both a significant positive and a significant negative log2FC across time, meaning that both a positive shift and negative shift from the average with an adjusted p-value <0.05 had been detected around the day-night cycle. This pipeline successfully identified diurnal shifts in clock gene expression (e.g. *Bmal1, Clock*) despite low cellular detection rates (**Table S3**), highlighting the strength of this approach. Pathway enrichment analysis was performed using Ingenuity Pathway Analysis^103^ (Qiagen) on genes upregulated at ZT0 and ZT12 resulting from this analysis. The results of this filtered-MAST analysis were also used to identify biologically relevant genes, such as the CME-associated genes, changing around the clock in BAMs.

Finally, we used both pseudobulk and cell-level analyses to compare BAMs from adult versus aged mice. scRNA-seq datasets were merged as described in the scRNA-seq integration section. The AggregateExpression function from Seurat v5 was used in conjunction with DESeq2 to compare all adult and aged samples acquired at ZT0 and ZT6 (for each age, samples were pooled across time). An adjusted p-value cutoff of 0.05 and a log_2_ fold-change (FC) cutoff of 0.58 were employed, corresponding to approximately a 50% change in expression. Pathway enrichment analysis was performed using IPA (Qiagen) on genes both up- and down-regulated in aged BAMs, obtained from pseudobulk analysis.

### Bulk RNA sequencing

#### Sample preparation and sequencing

Following FACS purification, cells underwent a second round of FACS due to the high propensity of BAMs to form doublets. BAMs are a relatively scarce cell type (typical yield of ∼10,0000 per brain). Cell loss from double FACS purification and toxicity of the transcription/translation inhibitors left us with a final yield of only ∼500-1000 BAMs per sample. Final yielded cells were spun down and re-suspended in TRIzol (Invitrogen #15596026) for RNA extraction. For RNA isolation, 200uL of Chloroform was added to each sample. Samples were shaken, incubated at room temperature for 3 minutes, then spun down for 15 minutes at 12,000g at 4C for phase separation. The aqueous phase, containing RNA, was isolated and labeled with GlycoBlue (ThermoFisher Scientific #AM9516). Samples were subsequently incubated for 10 minutes following addition of 500uL of RNAse-free isopropanol (Sigma-Aldrich #67-63-0), then spun down for 10 minutes at 12,000g at 4C. Following supernatant removal, RNA pellets were resuspended in 1mL of 75% ethanol then centrifuged for 5 minutes at 12,000g at 4C. Finally, the supernatant was removed, samples were air dried for 5 minutes, and RNA pellets were re-suspended in a final volume of 10uL of RNAse-free water and heated to 55C for 5 minutes. Samples were then stored at -80C until shipping for ultra-low input bulk RNA sequencing (Novogene).

#### Bulk RNA sequencing analysis

Pairwise mRNA-seq analysis was performed on all samples. Sequence reads from each sample were aligned to the GENCODE M37 (GRCm39) mouse reference genome. Initial QC on fastq reads was performed with FastQC (v0.12.1) and MultiQC (1.0.dev0), and the reads were trimmed using Trim Galore (v0.6.10). The trimmed reads were aligned to the GENCODE M37 (GRCm39) mouse reference genome with STAR Aligner (v2.7.11a) and TPM values were calculated with RSEM (v1.3.3). To confirm that the Bmal1 exon 8 knockout was successful, we mapped mRNA reads from ZT0 samples using with ggsashimi (v1.1.5)^104^ with the coordinates: chr7:112,878,525-112,886,617 and grouped by genotype.

### Immunohistochemistry on brain sections

#### Tissue collection, processing, and sectioning

Mice were deeply anesthetized via intraperitoneal injection of Avertin (2,2,2-Tribromoethanl in 2-Methyl-2-butanol; Millipore Sigma #T48402 and #240486) diluted in Hank’s Balanced Salt Solution (HBSS, ThermoFisher Scientific #14175145). The plane of anesthesia was verified by loss of withdrawal reflex to the paw pinch test and loss of palpebral reflex to the eye-blink test. At this stage, a subset of mice received intravenous delivery of a conjugated antibody for labeling vasculature (anti-CD31-BV421, clone 390, Biolegend #102424, 1µg/g). The antibody was allowed to circulate for only three minutes prior to the beginning of transcardial perfusion. For all mice, transcardial perfusion was performed with ice-cold HBSS at a volume of ≥1mL per gram of body weight. Following decapitation, the skull cap was carefully removed, ensuring that the dura mater remained attached to the skull cap rather than the surface of the brain. The brain was then removed and drop-fixed for 24 hours in 4% PFA (diluted in HBSS from EM grade 16% PFA, Electron Microscopy Science #15710), and subsequently transferred to 30% sucrose (Sigma-Aldrich #S0389, in HBSS) for 48 hours. Brains were then transferred to a 2:1 mixture of 30% sucrose and O.C.T. (Sakura Finetek #4583) for 24 hours, and half brains (cut along the sagittal midline) were individually placed in embedding molds, frozen in the 2:1 mixture, and stored at -80C. Free-floating 40μm sagittal sections were obtained used a cryostat and stored in HBSS containing 0.1% sodium azide (Sigma-Aldrich #71289) at 4C prior to use.

#### Immunohistochemistry

Sagittal sections were washed 3 x 5min in HBSS on a gentle shaker. Sections were then incubated for 1 hour at room temperature in a blocking and permeabilization buffer composed of 20% donkey serum (Sigma-Aldrich #D9663) or horse serum (Vector Laboratories #S-2000-20) in 0.3% Triton-X100 (Sigma-Aldrich #9036-19-5) in HBSS. Then, sections were transferred to the primary antibody staining solution composed of the antibodies and 20% serum in 0.3% Triton-X100 and incubated overnight at 4C. The following day, sections were washed 3 x 10min in 0.3% Triton-X100. If a secondary antibody stain was needed, sections were then transferred to the secondary staining solution (with 20% serum in 0.3% Triton-X100) overnight, followed by another 3 x 10min in 0.3% Triton-X100 wash. Blocking solution, primary antibody solution, and secondary antibody solution were used at a volume of 100uL/section. For all samples, after the final 0.3% Triton-X100 wash, sections were washed 2 x 5min in HBSS and gently mounted onto microscope slides (Leica #3800200) using a fine-tipped paintbrush. Once the sections were dry, ProLong Diamond Antifade Mountant (Thermo Fisher #P36970) was applied and slides were cover-slipped. The exception to this protocol was for the use of the Amylo-Glo reagent (Biosensis #TR-300-AG). In this case, free-floating sections were stained following Amylo-Glo product specifications prior to the beginning of the immunohistochemistry pipeline. Then, all steps (blocking, primary stain, secondary stain, and washes) occurred as described above, with the added consideration of pH verification for all solutions to avoid fading of the Amylo-Glo signal. Furthermore, Amylo-Glo stained sections were mounted in Aqua-Poly/Mount (Polysciences #18606-20) due to pH sensitivity.

### Cell culture and immunocytochemistry

#### Primary culture of BAMs

Following FACS purification, cells were centrifuged at 500 xg for 5 min at 4C and resuspended in BAM culture media containing DMEM/F-12 + GlutaMAXTM (ThermoFisher #10565-042), 10% FBS (FBS heat inactivated prior to use; ThermoFisher #26140079), 1x Penicillin-Streptomycin (ThermoFisher #15140122), and 75 ng/ul recombinant mouse Macrophage colony stimulating factor (M-CSF; R&D Systems #416-ML-050). 10,000 cells were plated per well of 96-well glass-bottom plate (Cellvis #P96-1.5H-N, pre-coated for 1 hour in 1x Poly-D-Lysine). Cells were plated overnight at 37C with 5% CO2.

#### In vitro engulfment assay and immunocytochemistry

Cultured BAMs were incubated for 30 minutes at 37C with 50ug/mL of Beta-Amyloid (AnaSpec #AS-20276) in BAM culture media and immediately processed for immunocytochemistry. All steps were carried out within the 96-well glass-bottom plate precoated with Poly-D-Lysine (PDL). Following every step, supernatant was removed gently with an aspirator, careful not to disrupt cells. All washing steps used a volume of 200uL per well and blocking, permeabilization, and staining steps were done at 100uL per well. Cells were fixed with 4% PFA in HBSS for 20 minutes at 4C, 3x washed with HBSS, and blocked with in a permeabilization solution containing 1x IC permeabilization buffer (eBioscience #00-8333-56), Fc block (1:50; BD #553142), and 20% horse serum (Vector Laboratories; #S-2000-20) in HBSS for 10 minutes at room temperature. Cells were then stained for 1 hour at room temperature and light-protected with an antibody cocktail containing AF488-conjugated anti-F4/80 (1:100; BioLegend #123120), AF647-conjugated anti-β-Amyloid_1-16_ (clone 6E10; 1:100; BioLegend #803020), and DAPI (1:10,000; BioLegend #422801) in the permeabilization solution. Finally, cells were washed 2X with HBSS and stored at 4C in HBSS with a paraffin seal until imaging.

### Image acquisition and analysis

#### Confocal microscopy

Cover-slipped slides or paraffin-sealed plates were imaged using a Zeiss LSM880 confocal microscope with the Zen Black 2.3 software. The Zeiss LSM880 was used with 405nm, 458 nm, 488nm, 561 nm, 594 nm, 633 nm, and 730 nm laser excitation. Fluorescent signal was captured by five detectors: two external (fixed band pass filters; for far-red and infrared signal) and three internal (adjustable band pass filters). Individual z-stacks or tile scans were taken with either a 10x air objective, a 20x air objective, or a 40x oil-immersion objective. Within confocal microscopy imaging batches, tile region, z-resolution, zoom, laser power, detector gain, and aperture were all matched across samples.

#### Slide scanner microscopy

Cover-slipped slides were imaged using a Zeiss Axioscan 7 slide scanner with Zen 3.4 software. Sample detection and section segmentation was done using brightfield imaging. Whole-section images were acquired using a 20x objective and fluorescent imaging using the DAPI, AF488, AF555, AF647, and AF750 channels. Exposure duration and laser power were set per channel. Coarse focus was obtained throughout the section using the 5x objective and the onion skin strategy on the AF488 channel. Fine focus was obtained for each tile using the onion skin strategy on the AF488 channel. Within slide-scanner imaging batches, all imaging and focus parameters were matched across samples.

#### Analysis of amyloid pathology from immunohistochemical staining

Amyloid pathology was assessed in 3-5 whole sagittal sections per mouse via slide-scanner microscopy using three different pathological markers per panel. Aβ plaque was stained with Amylo-Glo, Aβ_1-16_ was labelled using an AF647-conjugated anti-β-Amyloid antibody (6E10 clone, Biolegend #803021), and APP was labelled using an anti-APP antibody (1D1 clone, EMD Millipore #MABN2287) followed by an AF555 donkey anti-rat secondary antibody. The use of dyes and antibodies in combination allowed within-section validation that signal was not driven by false-positive or non-specific binding. Tiled images of each whole section were saved as 8-bit TIFF files. The hippocampal formation (including subiculum), leptomeninges, cortex, and whole section were manually segmented using anatomical landmarks such as fiber tracks, vasculature, neuronal organization, and ventricle localization, and verified using the Allen Mouse Brain Atlas^105^ sagittal view. For example, the dorsal leptomeninges were confirmed via the presence of vasculature running parallel to the surface of the brain. Below the leptomeninges, the cortex was distinguished from subcortical regions via fiber tracks. All cortical layers were included in the cortex segmentation. Within individual mice, signal from the same region across different sections was highly replicable, indicating reliable region of interest (ROI) segmentation. Following segmentation, images were binarized using manual thresholding for which the parameters were set by channel and equally across all images of a batch. Amylo-Glo, 6E10, and 1D1 signal, including particle size, total area, and ROI coverage, were obtained per ROI. Then, CD31 staining was used as a vascular landmark, and its colocalization with Amylo-Glo, 6E10, and 1D1 signal was calculated per ROI. To account for inherent differences in ROI size across sections, all data were calculated as a percentage of the total ROI size, with the exception of plaque size and plaque number. Plaque size values were measured in μm^2^ and averaged across individual sections and mice. Plaque number values were divided by ROI area to normalize across sections and measured in mm^2^. All image analysis was automated and performed blind to genotype to prevent experimenter bias.

### Acute sleep deprivation

Mice were acutely and completely sleep deprived starting at either ZT0 or ZT6 for a duration of six hours. Sleep deprived mice all remained group-housed with *ad libitum* access to food and water throughout the procedure. Sleep deprivation was achieved via gentle handling, cage tapping, and novel object introduction^106^. Mice were continuously monitored by at least one experimenter to ensure continued wakefulness, and were not disturbed if showing spontaneous behaviors including movement, grooming, or food or water intake. Sleep deprivation continued all the way to the point of intraperitoneal Avertin delivery prior to cardiac perfusion for all mice. Control mice were similarly transported out of the animal colony housing room to control for effects of transit, but were allowed to sleep uninterrupted for the duration of the study.

### Western blot analysis

Tibial and femoral bone marrow and brain were removed from P136 male 5xFAD mice and wildtype littermates, frozen on dry ice and stored at -80. Tissues were lysed in RIPA buffer (PI89900 Thermo Scientific) supplemented with protease inhibitor cocktail (P8340, Merck) using the Precellys bead-milling method (Precellys soft tissue homogenizing lysis kit, Bertin Instruments). Insoluble material was removed by centrifugation (13,000 RPM, 4C, 10min) and protein concentration of the supernatant was measured using Pierce™ BCA Protein Assay Kit (Thermo Scientific 23225). 30ug of protein samples were run on a Tris-tricine SDS–PAGE gels (10–20%, Novex, Thermo Fisher). Gels were run at 100–120 V for approximately 1 h. Proteins were transferred onto low-fluorescent Immobilon-FL membrane (0.45 µm pore size, Merck) using the Bio-Rad wet-blot system (1.15 h, 500 mA) and blot transfer buffer (25 mM Tris and 190 mM glycine) containing 20% methanol. Blots were washed in water, and transferred protein was stained with Fastgreen (0.0005% Fastgreen FCF (Serva)) for whole protein loading control. Membranes were imaged using a ChemiDoc MP Imaging system. Membranes were rinsed and blocked in 5% BSA in TBS-T for 1 h at room temperature. Membranes were then incubated in primary antibodies in 5% BSA overnight at 4 °C on a rotating shaker (1:1,000; mouse, 6E10, BioLegend). Membranes were washed several times, and incubated in secondary antibody solution (5% BSA in TBS-T, 1:1000 goat anti mouse IRDye® 680 Goat) for 1h at RT, subsequently washed and imaged at the ChemiDoc system.

### Statistical analyses

Statistical analyses were performed using R (version 4.4.2) driving RStudio (version 2024.09.1+394, Posit PBC) or JMP Pro (version 18, JMP Statistical Discovery). Statistical testing on scRNA-seq data is described in the single-cell RNA sequencing methods section. For other comparisons, the statistical test used is indicated in figure legends. Two-tailed tests with an alpha cut-off of 0.05 were used unless otherwise specified. For comparisons between two independent groups, we used either parametric or non-parametric comparisons depending on data distribution and sample size: unpaired Student’s t-tests or Wilcoxon rank-sum tests, respectively. Grouped comparisons were performed using one- or two-way ANOVAs. In some instances, mixed effects models using restricted maximum likelihood (REML) approaches were used to account for random variables (for example, experimental batch), in which case model parameters are included in the figure legend. For all ANOVAs and mixed effects testing, main and interaction effects are indicated and post-hoc testing was performed with Tukey’s HSD (Honestly Significant Difference). Data visualization was performed using Rstudio, JMP Pro, and FlowJo.

## RESOURCE AVAILABILITY

A web-based interface for browsing the circadian scRNA-seq atlas will be made publicly available and all sequencing data will be deposited in the Gene Expression Omnibus (GEO) upon manuscript publication. No new reagents were generated in this study, however, requests for protocol sharing are gladly accepted.

## ACKNOWLEDGEMENTS

We thank Arnaud Frouin for lab management, Joel Cuadrado for animal care, Emma Connolly for coordinating experiments at the Broad Institute, and Daniel Wilton, Tomasz Kula, Jordan Doman, and Nils Korte for thoughtful feedback on this manuscript. Ronald Mathieu from the HSCI-BCH Flow Cytometry Research Laboratory provided generous support with flow cytometric experiments. Liza Curtis, Krishna Narayanan, Anna Kane, and Bronwen Brown provided key administrative support. We also thank Nathaniel Hodgson and the IDDRC Animal Behavior and Physiology Core for support with circadian experiments and the Harvard Medical School O2 High Performance Compute Cluster for use in data preprocessing. We also thank the Simons Foundation SCPAB team and the many members of the Stevens Lab who provided guidance from this project’s inception. This work was generously supported by grants from the Simons Foundation Collaboration on Plasticity and the Aging Brain (B.S.), the Cure Alzheimer’s Fund (B.S.), a NIH Training Grant on Molecular Biology of Neurodegeneration and Alzheimer’s Disease T32AG000222 (H.J.B.), and the Howard Hughes Medical Institution (B.S.). Illustrations created in BioRender (Frouin, A. (2025) https://BioRender.com/lysfc8t).

## AUTHOR CONTRIBUTIONS

Conceptualization, H.J.B. and B.S. with input from C.D., A.J.W., and J.L.; Methodology, H.J.B., C.D., A.J.W., S.E.M., S.F., J.L.; Investigation, H.J.B., S.K.C., C.D., A.J.W., S.E.M., A.S.G., J.P, M.H., S-O.P., S.F.; Software, H.J.B., J.M.R., S.E.M.; Supervision, H.J.B. and B.S.; Formal analysis, H.J.B., J.M.R., S.E.M.; Visualization, H.J.B., J.M.R.; Project administration, H.J.B.; Writing – original draft, H.J.B. and B.S.; Writing – review & editing, all authors; Funding acquisition, H.J.B., A.J.W., C.D., B.S.

## SUPPLEMENTAL INFORMATION

Document S1. Figures S1-6 and Tables S1-6.

---

**Figure S1.**
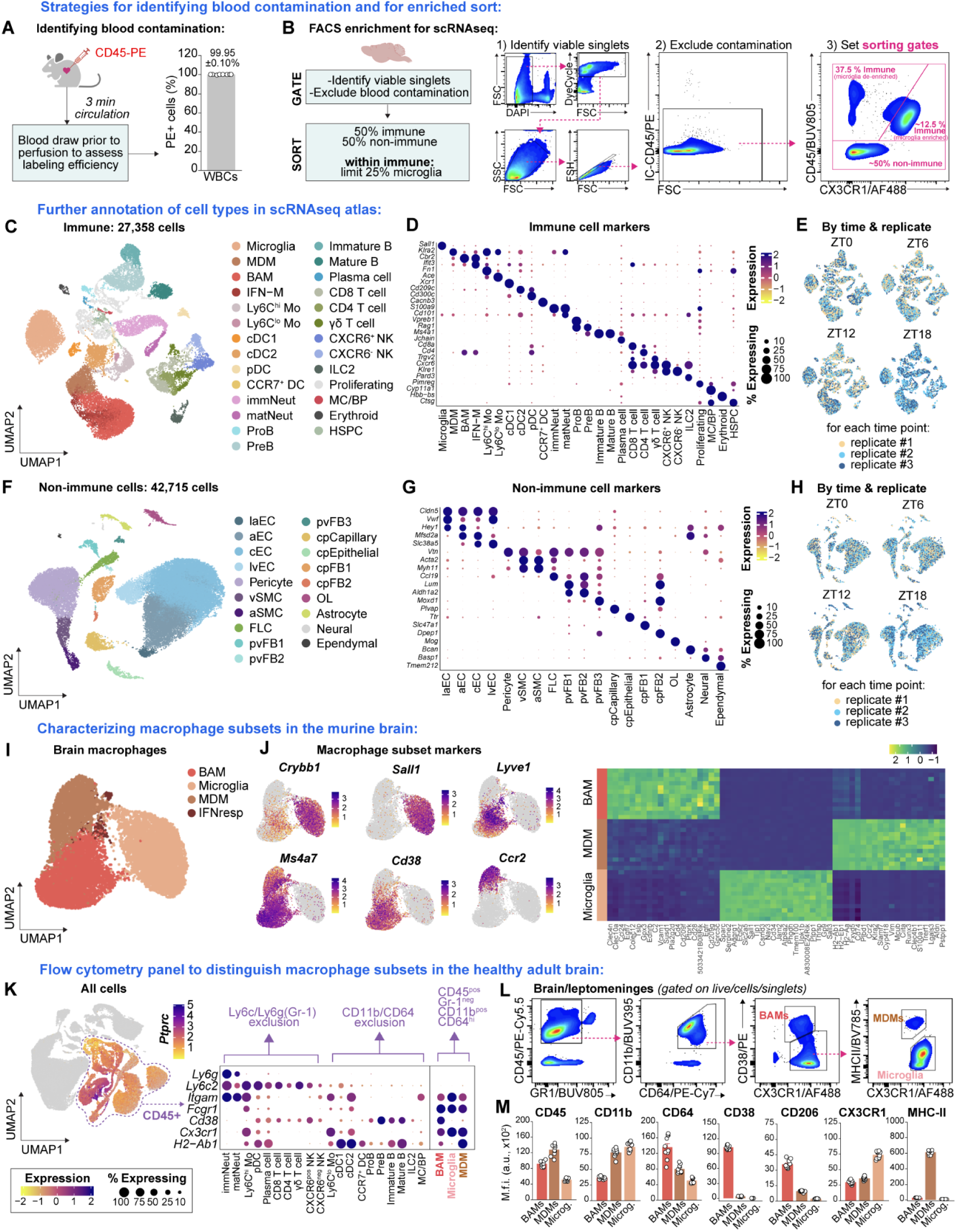
Additional annotation of circadian single-cell atlas and evidence-based flow cytometry panel for macrophage subset distinction in the healthy brain, related to Figure 1 (A) Experimental approach for labeling circulating immune cells prior to cardiac perfusion via intracardiac injection of CD45-PE antibody. Following 3 minutes circulation, and prior to cardiac perfusion, blood was acutely sampled to verify white blood cell (WBC) labeling efficacy. Quantification shows mean ± SEM. (B) Gating and sorting strategy prior to single-cell RNA sequencing to de-enrich microglia and enrich other brain-resident immune cells and their niches. Live, viable singlets were dually identified as DAPI^neg^ and Vybrant DyeCycle Ruby^pos^. (**C-E**) UMAP of annotated immune cell subsets, heat map of identification genes per cell type, and immune cells split by time and colored by replicate (n of 12 mice, 3 independent samples per time). IFN-M = interferon-responsive macrophages, immNeut and matNeut = immature and mature neutrophils. (**F-H**) UMAP of annotated non-immune cell subsets, heat map of identification genes per cell type, and non-immune cells split by time and colored by replicate (n of 12 mice, 3 independent samples per time). laEC = large arterial EC, aEC = arterial EC, cEC = capillary EC, lvEC = large venous EC, vSMC = venous SMC, aSMC = arterial SMC, pvFB = perivascular FB, cpFB = choroid plexus FB. (**I and J**) UMAP of macrophage subsets, feature plot of genes of interest, and heat map of macrophage subset-enriched genes across all 12 independent replicates. (**K-M**) Feature plot of *Ptprc* expression in whole scRNA-seq atlas, complex heat-map of immune cells demonstrating gene-level expression of markers used for macrophage subset differentiation by flow cytometry, recommended gating strategy, and group-level quantification of recommended surface markers from panel in BAMs, MDMs, and microglia. Points represent individual mice, n of 10 mice.

**Figure S2.**
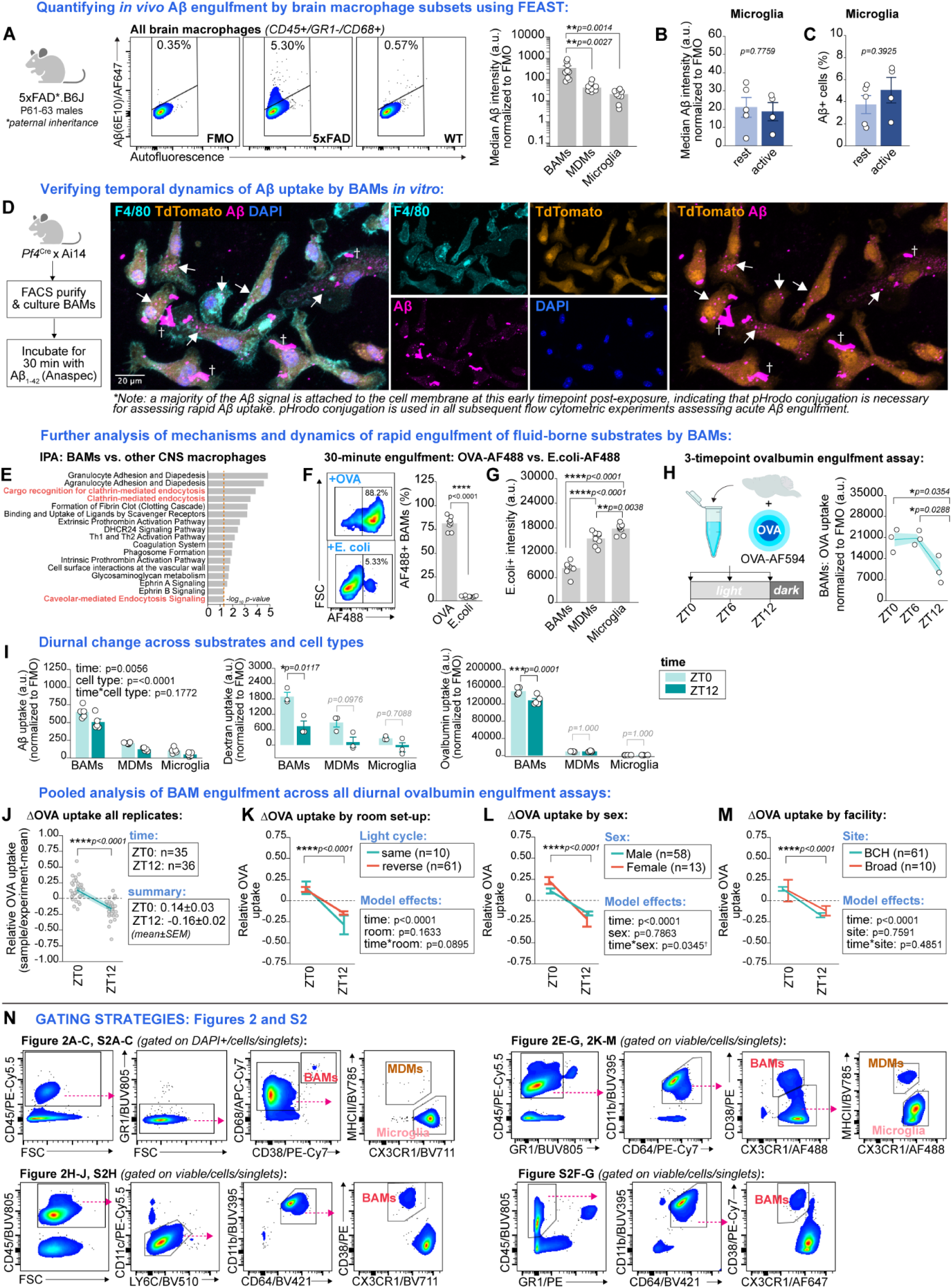
Further characterization of BAM engulfment and its rhythmicity, related to Figure 2 (**A**) Representative pseudocolored plots of Aβ signal in brain macrophages following the FEAST protocol on microdissected cortices from a young 5xFAD male versus a wild-type littermate control, and group-level quantification of Aβ intensity within each macrophage subset. Aβ signal varies by cell type (one-way ANOVA, F_2,24_=10.14, p=0.0006) and is highest in BAMs. N of 9 mice. The P61-63 timepoint is selected to maximize soluble versus insoluble Aβ. (**B and C**) Quantification of Aβ intensity and Aβ-positivity (%) within microglia, neither of which are significantly different across time at this early time point in amyloidosis. Quantification using two-tailed t-test, n of 4-5 mice per time point. (D) Experimental design and representative image of cultured BAMs following incubation with purified Aβ_1-42_. Aβ is detectable in intracellular vesicular structures within just 30 minutes of exposure (white arrowheads). Importantly, clumps of Aβ also adhered to the cell membrane (denoted by †), highlighting the need for pHrodo conjugation of all Aβ substrates used for measuring acute engulfment capacity. (E) IPA of top 100 genes upregulated in BAMs relative to microglia and MDMs (from cell-level analysis from all time points, n of 12, selected pathways shown). Dashed bar indicates significance threshold. (F) Representative pseudo-colored plots and quantification of substrate positivity in BAMs following 30 minutes digest with either AF488-conjugated ovalbumin or E.coli. Acute capacity for E.coli uptake is far reduced relative to OVA-AF488 uptake (t(7.18)=-29.86, p<0.0001, two-tailed t-test, n of 8). (G) Within E.coli-positive cells, signal varies by cell type (F_2,14_=135.74, p<0.0001). Signal is higher in both MDMs and microglia relative to BAMs (mixed effects model accounting for random variance due to individual sample, n of 8). (H) Experimental design and group-level quantification of OVA-AF594 uptake by BAMs at ZT0, ZT6, and ZT12. Uptake varies by time: F_2,6_=7.90, p=0.0209 (one-way ANOVA, post-hoc testing with Tukey’s HSD; n of 3 mice per time point), and is higher at both ZT0 and ZT6 relative to ZT12. (I) Group-level quantification of acute engulfment across cell types and substrates at ZT0 vs. ZT12. While BAMs consistently show a reduction in engulfment at ZT12 vs. ZT0 across substrates, MDMs and microglia do not. Analyzed with time*cell type full factorial mixed effects models accounting for random variance due to individual samples, n of 3-5 per time point). (**J-M**) Relative OVA uptake by BAMs at ZT0 versus ZT12 across 9 experimental replicates and 71 biological replicates (line and shaded area represent mean and 95% confidence intervals) and relative OVA uptake split by light cycle, sex, and animal facility. Model effects shown are for mixed effects design controlling for random variance due to independent experiments. Sample sizes denote biological replicates. There is a main effect of time on OVA uptake across all conditions. Lines and error bars represent mean and SEM. ^†^No significant post-hoc test for between-sex or within-time comparisons. (N) Flow cytometry gating strategies for data shown in Figure 2 and **Figure S2.** Note: the staining panel and gating strategy is adjusted for the fixed cell FEAST protocol due to epitope loss following sample preparation. *For all panels: points represent individual mice, bars and error bars represent mean and SEM. Normalization to background autofluorescence is done per cell type. Illustrations made with BioRender. ****p<0.0001, ***p<0.001, **p<0.01, *p<0.05*.

**Figure S3.**
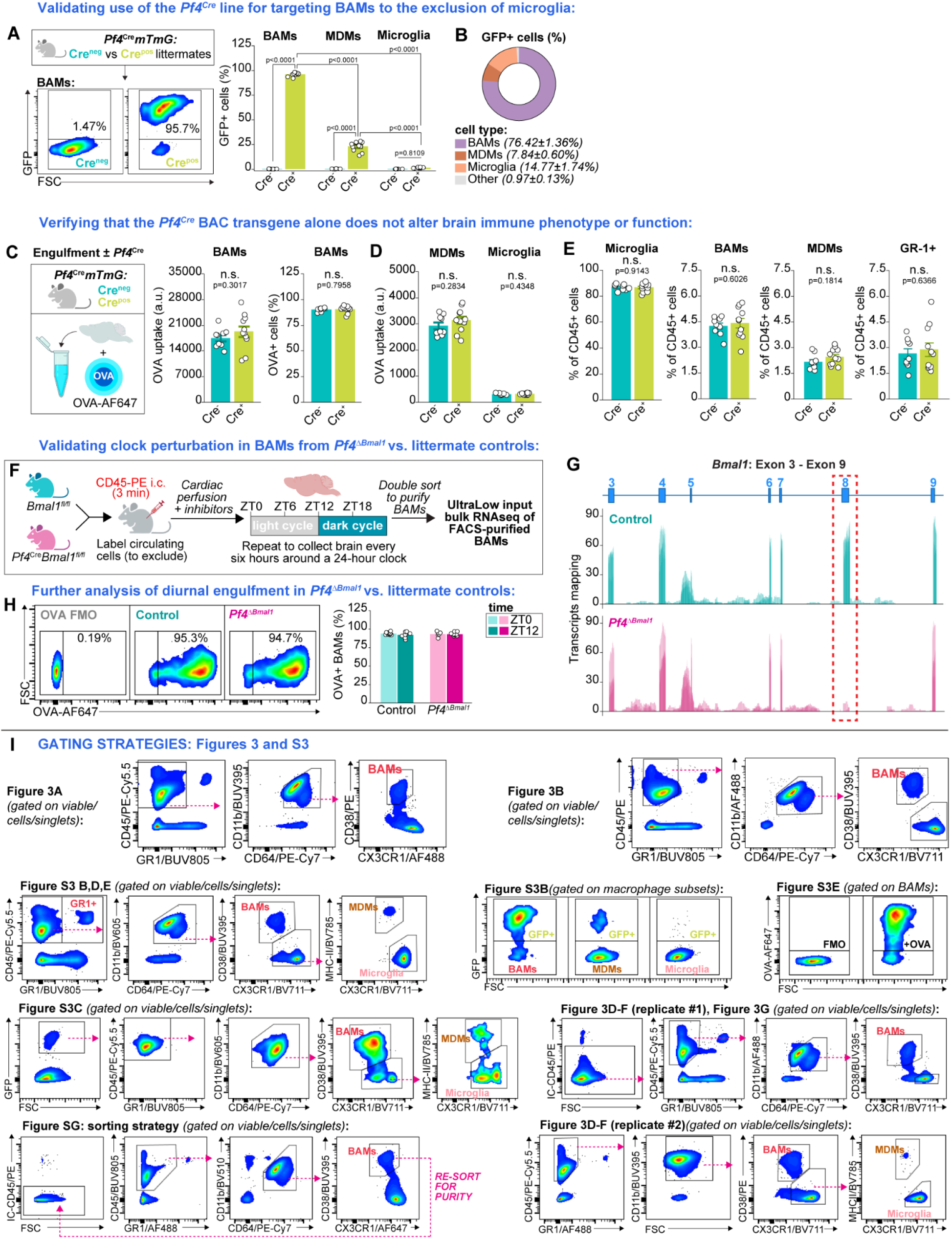
Validation of BAM-targeted circadian disruption, related to Figure 3 (A) Pseudo-colored plots and group-level quantification of GFP-positivity across cell types and genotypes in *Pf4*^Cre^*mTmG* mice. Cell type*genotype interaction: F_2,36_=3211.658, p<0.0001 (mixed effects model accounting for variance due to individual mouse). Selected post-hoc tests shown (all Tukey’s HSD; n of 9 Cre^neg^ and 11 Cre^pos^). (B) Cell type as % of all GFP+ cells in *Pf4*^Cre^*mTmG* mice. Data shown as average across all Cre^pos^ mice (n of 11). (**C and D**) Schematic of experimental design, group-level quantification of OVA-AF647 uptake and percent of BAMs positive for OVA-AF647, and group-level quantification of OVA-AF647 uptake by MDMs and microglia. There was no effect of genotype on OVA-AF647 uptake by BAMs, MDMs, or microglia, nor on the % of BAMs positive for OVA-AF647. All comparisons using student’s t-test (n of 9 Cre^neg^ and 11 Cre^pos^). (**E**) Cell count as % of all CD45+ by genotype. No effect of genotype on microglia, BAM , MDM , or GR-1 cell count (likely monocytes and granulocytes). All comparisons using student’s t-test (n of 9 Cre^neg^ and 11 Cre^pos^). (**F and G**) Schematic of experimental design for bulk RNA sequencing of BAMs from *Pf4^ΔBmal1^* mice and littermate controls and histogram of reads mapping to the loxP-flanked eighth exon of the *Bmal1* gene at ZT0 in both groups. (H) Pseudo-colored plots and group-level quantification of the percent of BAMs containing detectable OVA-AF647 across time and genotype. No effect of either time nor genotype of OVA-AF647 positivity (F_3,33_=0.6609, p=0.5820; ANOVA; n of 21 control and 16 *Pf4^ΔBmal1^*). (I) Flow cytometry gating strategies for Figure 3 and **Figure S3**. *For all panels: points represent individual mice, bars and error bars represent mean and SEM. Illustrations made with BioRender. ****p<0.0001, ***p<0.001, **p<0.01, *p<0.05*.

**Figure S4.**
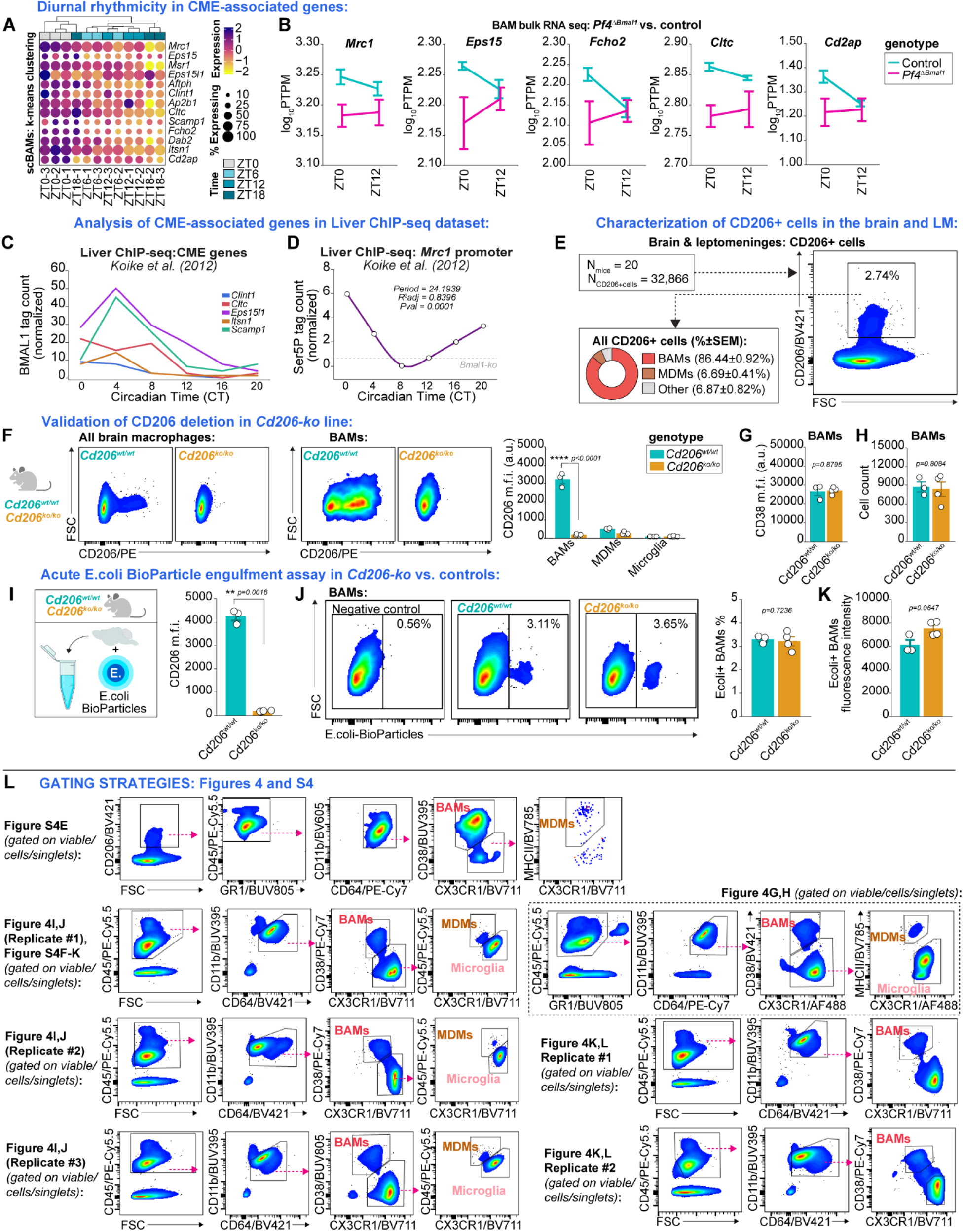
Further exploration of diurnal rhythmicity in CME-associated and validation of CD206 deletion, related to Figure 4. (A) Heatmap of CME-associated gene expression across time and biological replicates in BAMs (n of 3 mice per time point) with unbiased k-means clustering of samples by feature expression. (B) Group-level quantification of log_10_-transformed pseudo-transcripts per million (PTPM) of CME-associated genes at ZT0 and ZT12 in BAMs from Pf4^ΔBmal1^ mice versus their littermate controls. (**C and D**) Visualization of BMAL1 and Ser5P tagging of genes of interest across time in ChIP-seq dataset of liver (Koike et al., 2012). (E) Psuedo-colored plot of CD206+ cells in brain and quantification of cell type across 20 mice (data re-analyzed from Figure S2.3B-F). BAMs represent 85.59±0.90% of CD206+ cells in the brain (mean ± SEM). (F) Representative pseudo-colored plots and group-level quantification of CD206 expression on brain macrophage subsets from Cd206^wt/wt^ and Cd206^ko/ko^ mice. Cell type*genotype interaction: F_2,14_=151.84, p<0.0001 (mixed effects model accounting for individual mouse; post-hoc testing with Tukey’s HSD; selected post-hoc tests shown). Points represent individual mice (n of 3 Cd206^wt/wt^ and n of 4 Cd206^ko/ko^). (**G and H**) Quantification of CD38 expression and BAM yield from Cd206^wt/wt^ and Cd206^ko/ko^ mice. Both CD38 expression (t(2.64)=0.17, p=0.8795) and yield are unaffected by genotype (t(4.90)=0.26, p=0.8084; two-tailed t-tests). Points represent individual mice (n of 3 Cd206^wt/wt^ and n of 4 Cd206^ko/ko^). (**I-K**) Experimental design, representative pseudo-colored plots, and group-level quantification of E.coli BioParticle uptake in BAMs from Cd206^ko/ko^ mice and their wild-type littermate controls. The knockout was verified by loss of CD206 expression relative to wild-type littermates (t(2.02)=22.93, p=0.0018; two-tailed t-test). Both the proportion of BAMs positive for E.coli BioParticles and the intensity of signal within E.coli+ BAMs were unaffected by genotype (t(4.25)=0.38, p=0.7236 and t(3.69)=2.61, p=0.0647, respectively; two-tailed t-tests). Points represent individual mice (n of 3 Cd206^wt/wt^ and n of 4 Cd206^ko/ko^). (L) Flow cytometry gating strategies for Figure 4 and **Figure S4**. For all panels: points represent individual mice, bars and error bars represent mean and SEM. ****p<0.0001, ***p<0.001, **p<0.01, *p<0.05.

**Figure S5.**
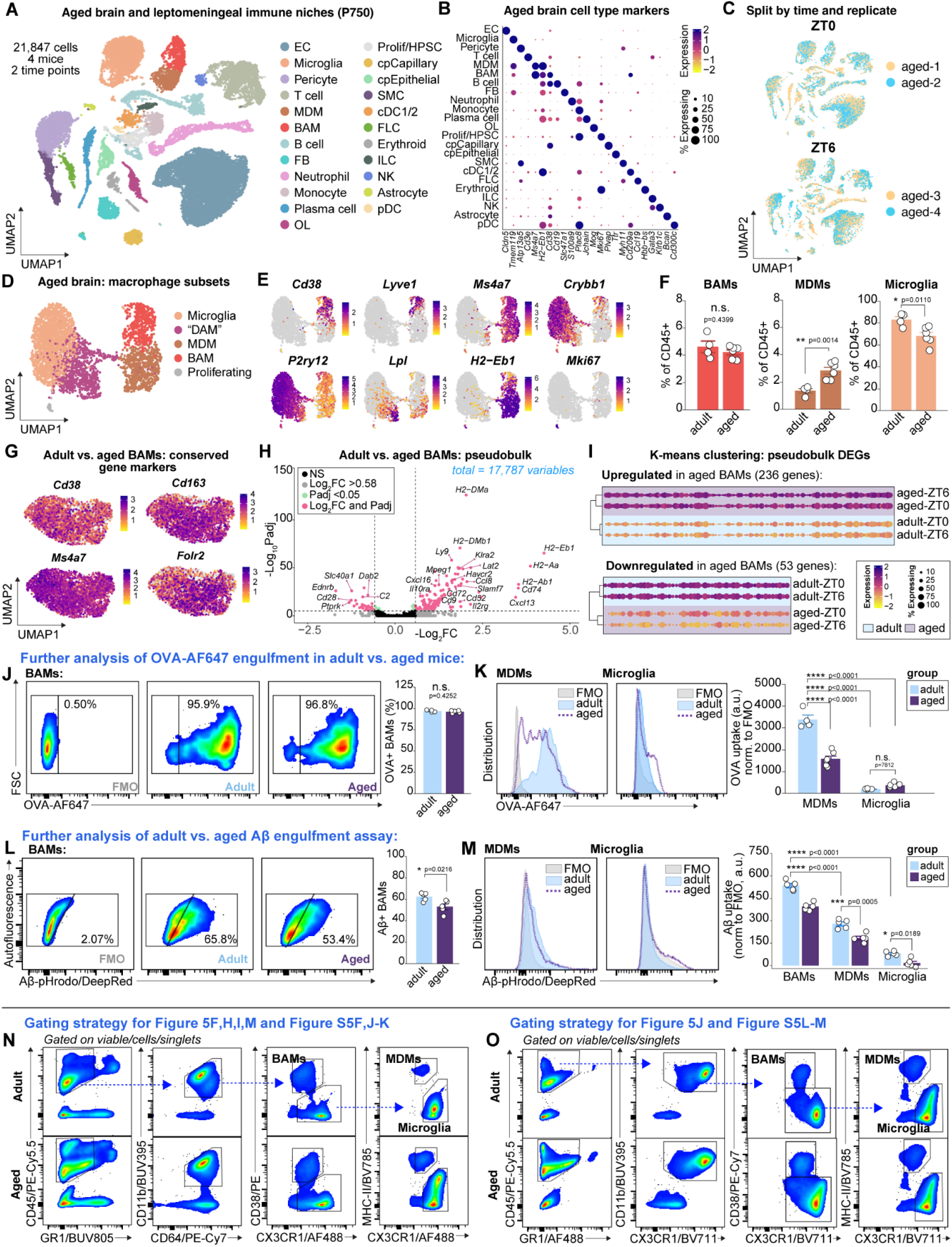
Additional analysis of macrophages subsets in the adult and aged brain, related to Figure 5 (**A-C**) UMAP of annotated cell types, heat map of cell type markers, and dataset split by time and replicate (n = 2 independent samples per time point). (**D and E**) UMAP of macrophage subsets and feature plots of identification genes per subset (from aged brain dataset; 3,573 macrophages). DAM = “disease-associated microglia”. (F) BAM, MDM, and microglia counts by flow cytometry, shown as % of CD45-positive cells in the brain and leptomeninges. BAM counts are unchanged by age (t(4.17)=-0.85, p=0.4399), but MDMs are increased (t(7.81)=4.84, p=0.0014) and microglia are decreased with age (t(7.84)=-3.31, p=0.0110; two-tailed t-tests). Points represent individual mice, n = 4 adult and 6 aged mice, all male B6N. (G) Feature plot of conserved genetic markers of BAM identity across age. (H) Volcano plot comparing BAMs from adult vs. aged mice via pseudobulk analysis; n = 10 total independent samples (6 adult, 4 aged, both ZT0 and ZT6 from each age included in comparison, all samples obtained in same experiment/preparation). FC threshold set to 0.58 (equal to a 1.5-fold change in expression). (I) K-mean clustering of DEGs from pseudobulk analysis, split by pattern of expression. For both up- and down-regulated genes, unbiased clustering still distinguishes adult from aged BAMs across ZT0 and ZT6. (J) Representative pseudo-colored plots and group-level comparison of the proportion of BAMs containing detectable levels of OVA-AF647 from adult vs. aged OVA engulfment assay (Figure 2**.5I**). The percent of OVA+ BAMs is unchanged by age (t(6.76)=-0.85, p=0.4252; two-tailed t-test with independent samples). Points represent individual mice, n = 4 adult and 6 aged mice, all male B6N collected at ZT0. (K) Representative histograms and group-level quantification of OVA-AF647 uptake in MDMs and microglia from adult vs. aged mice: age*cell type interaction (F(3,16)=129.60, p<0.0001; mixed effects model with random variable of individual mouse). Uptake was reduced in aged MDMs relative to adult MDMs (p<0.0001). Regardless of age, MDMs surpassed microglia in uptake (all p<0.0001). Microglia engulfment was unaffected by age (p=0.7812). (L) Representative pseudo-colored plots and group-level comparison of the proportion of BAMs containing detectable levels of Aβ-pHrodo from adult vs. aged Aβ engulfment assay. The percent of BAMs with detectable Aβ over background is reduced in aged BAMs (t(6.89)=-2.96, p=0.0216; two-tailed t-test with independent samples). Points represent individual mice, n = 5 mice per age. (M) Representative histograms and group-level quantification of Aβ uptake across cell types and ages. Age*cell interaction: F(2,16)=8.99, p=0.0024 (mixed effects model with random variable of individual mice, n = 5 mice per age). (**N and O**) Gating strategies for Figure 5 and **Figure S5**. The same gating strategy was applied to adult and aged samples from each experiment and was compatible with the increased autofluorescence and leukocyte infiltration present in the aged brain. *For all panels: points represent individual mice, bars and error bars represent mean and SEM. FMOs were generated per age to account for differences in background autofluorescence. Post-hoc testing with Tukey’s HSD. Illustrations made with BioRender. ****p<0.0001, ***p<0.001, **p<0.01, *p<0.05*.

**Figure S6.**
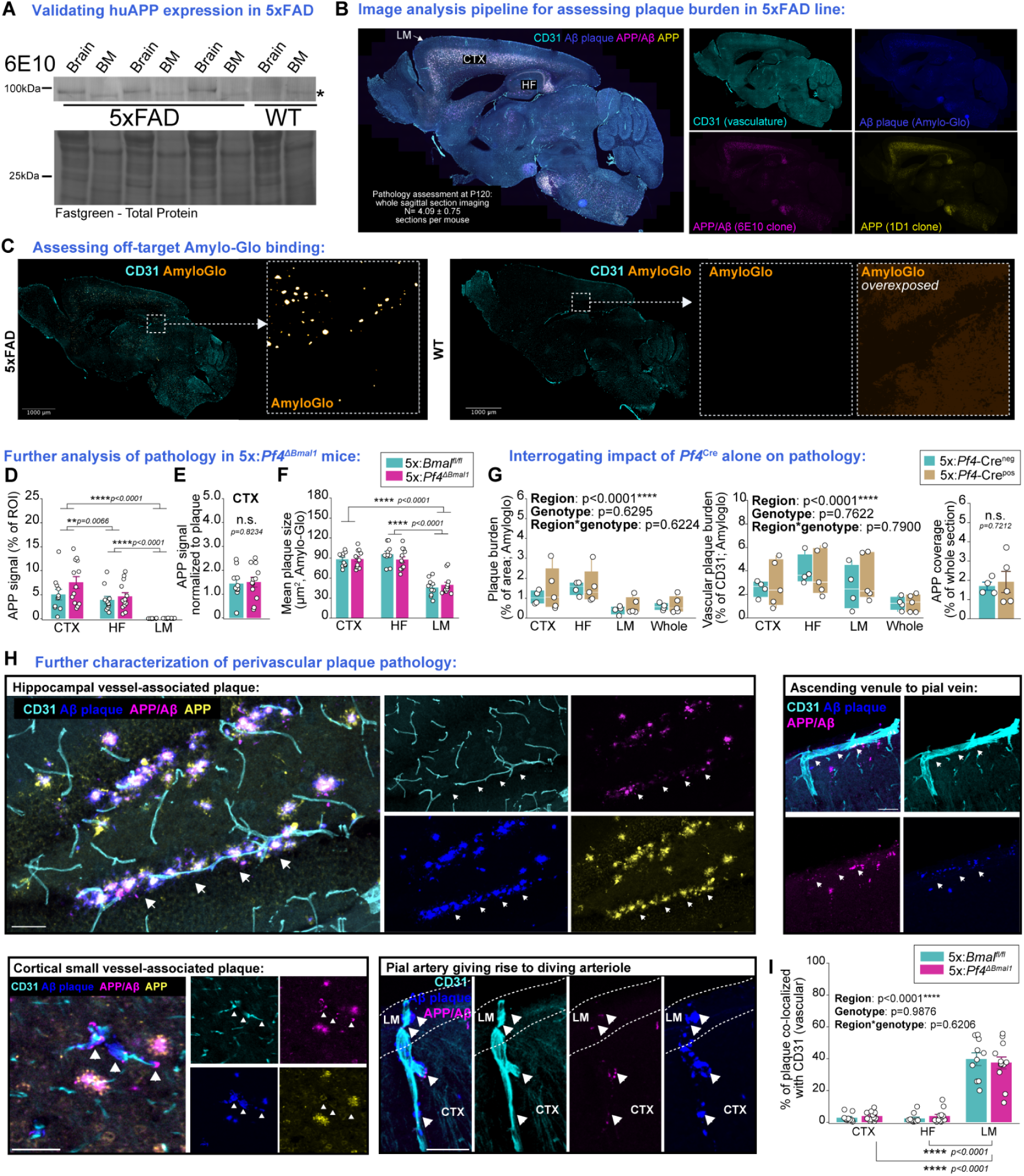
Further characterization of brain border amyloid pathology, related to Figure 6. (A) Human APP expression across brain and bone marrow (BM) of n of 3 5xFAD mice and a wild-type littermate control. Full-length human APP is only detected in the brain, but not bone marrow, of 5xFAD samples, and is not detected in the wild-type. *Denotes non-specific band. (B) Representative sagittal section slide-scanner microscopy image from a 5xFAD brain showing anatomical regions segmented for further analysis: the dorsal leptomeninges (LM), cortex (CTX), and hippocampal formation (HF). (C) Representative whole-section images of Amylo-Glo staining in 5xFAD versus wild-type mouse. No signal is detected in the wild-type mouse. (D) Quantification of APP signal across regions and genotypes. Very little APP is detectable above background in the leptomeninges. While there was a main effect of region on APP signal (p<0.0001), there was no effect of genotype (p=0.2419) nor region*genotype interaction effect (p=0.1373). Quantified using mixed effects model controlling for variance due to individual mouse and sex; post-hoc testing with Tukey’s HSD. N of 10-12 per genotype. (E) Cortical APP signal normalized to cortical plaque burden in 5x:*Pf4^ΔBmal1^* mice versus their littermate controls. Quantified with two-tailed t-test, n of 10-12 per genotype. (F) Group-level quantification of plaque size per genotype and region. Main effect of region (p<0.0001), but not genotype (p=0.7599), and no region*genotype interaction (p=0.2825) on plaque size. Quantified using mixed effects model controlling for variance due to individual mouse and sex; post-hoc testing with Tukey’s HSD. N of 10-12 per genotype. (G) Group-level quantification of amyloid pathology in 5xFAD mice expressing *Pf4*^Cre^ (5x:*Pf4*^Cre^) versus their Cre-negative littermates (5xFAD). CTX = cortex, HF = hippocampal formation, LM = leptomeninges. No effect of genotype on plaque burden or vascular plaque burden (mixed effects models controlling for mouse and sex), nor on APP coverage (two-tailed student’s t-test). N of 4 to 5 per genotype. (H) Further representative confocal microscopy images of perivascular plaque hippocampal small vessels, cortical small vessels, and diving pial arteries. Perivascular plaques are indicated via white arrowheads. Scale bars = 50μm. (I) Group-level quantification of plaque colocalized with CD31 per genotype and region. Main effect of region, but not genotype, on likelihood of plaque to be vessel-associated (quantified using mixed effects model controlling for variance due to individual mouse and sex; post-hoc testing with Tukey’s HSD). N of 10-12 per genotype. *For all panels: points represent individual mice, bars and error bars represent mean and SEM, box-and-whisker plots represent median and IQR (points unconnected to whiskers are considered outliers by IQR). ****p<0.0001, ***p<0.001, **p<0.01, *p<0.05*.

**Table S1.**
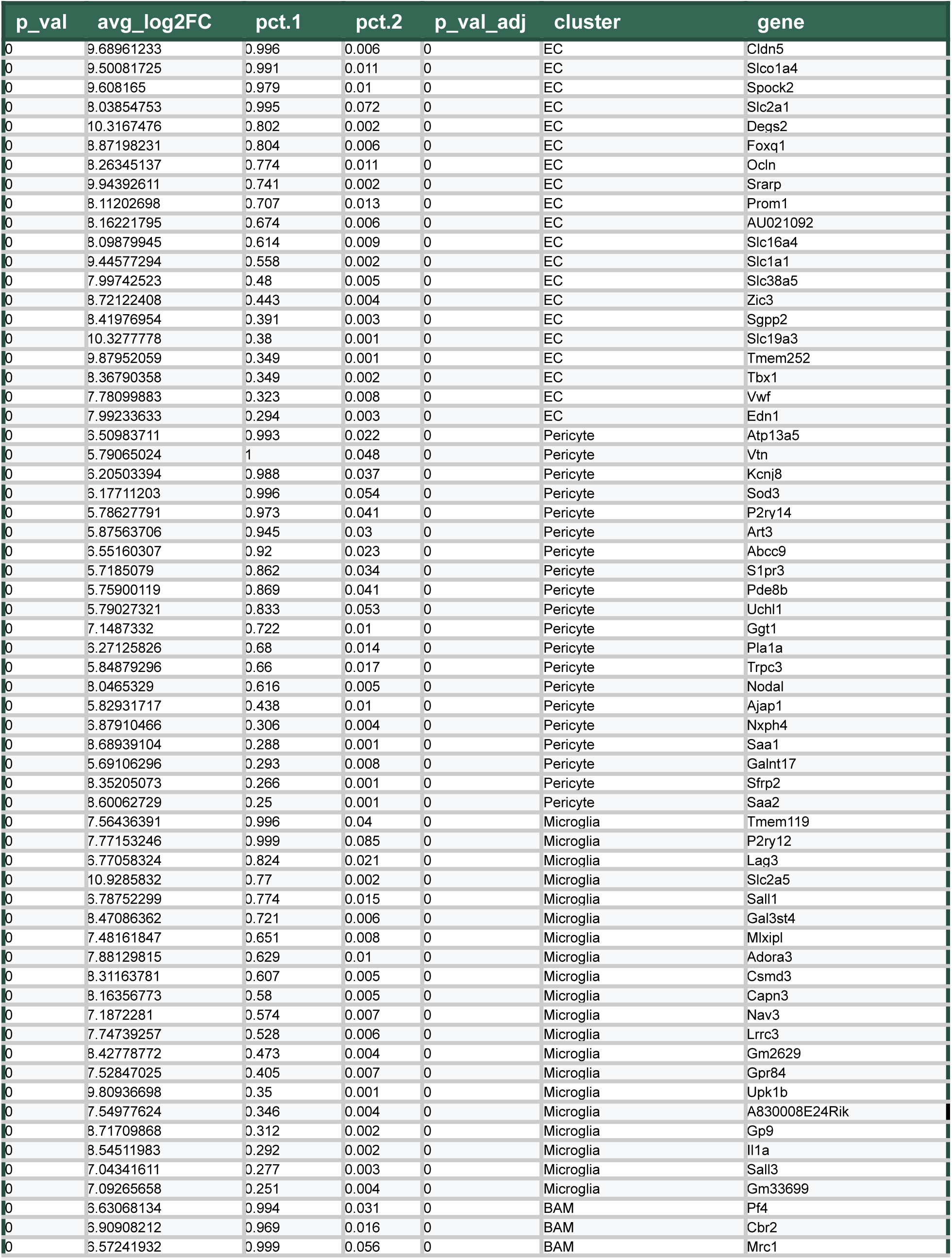
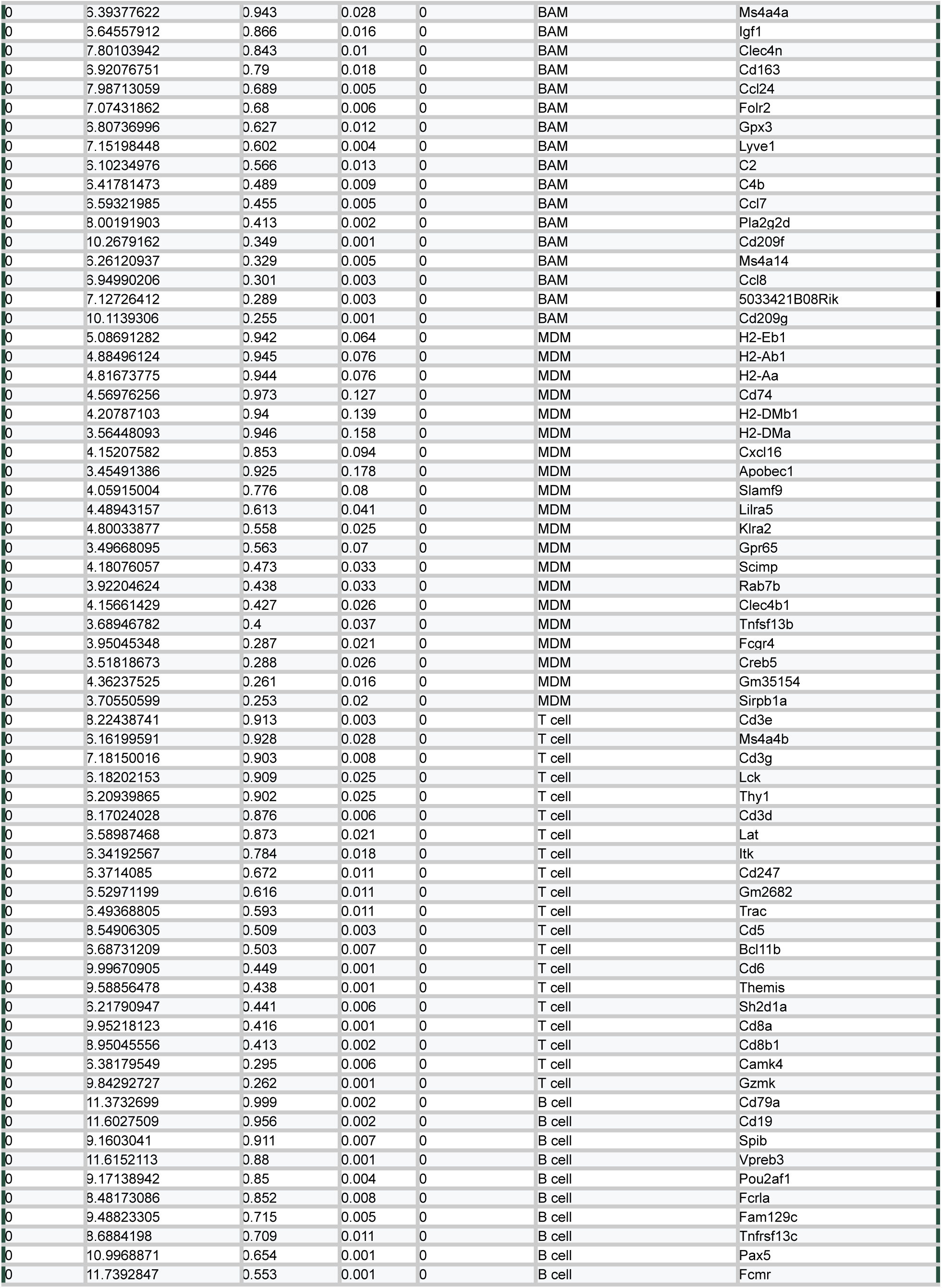
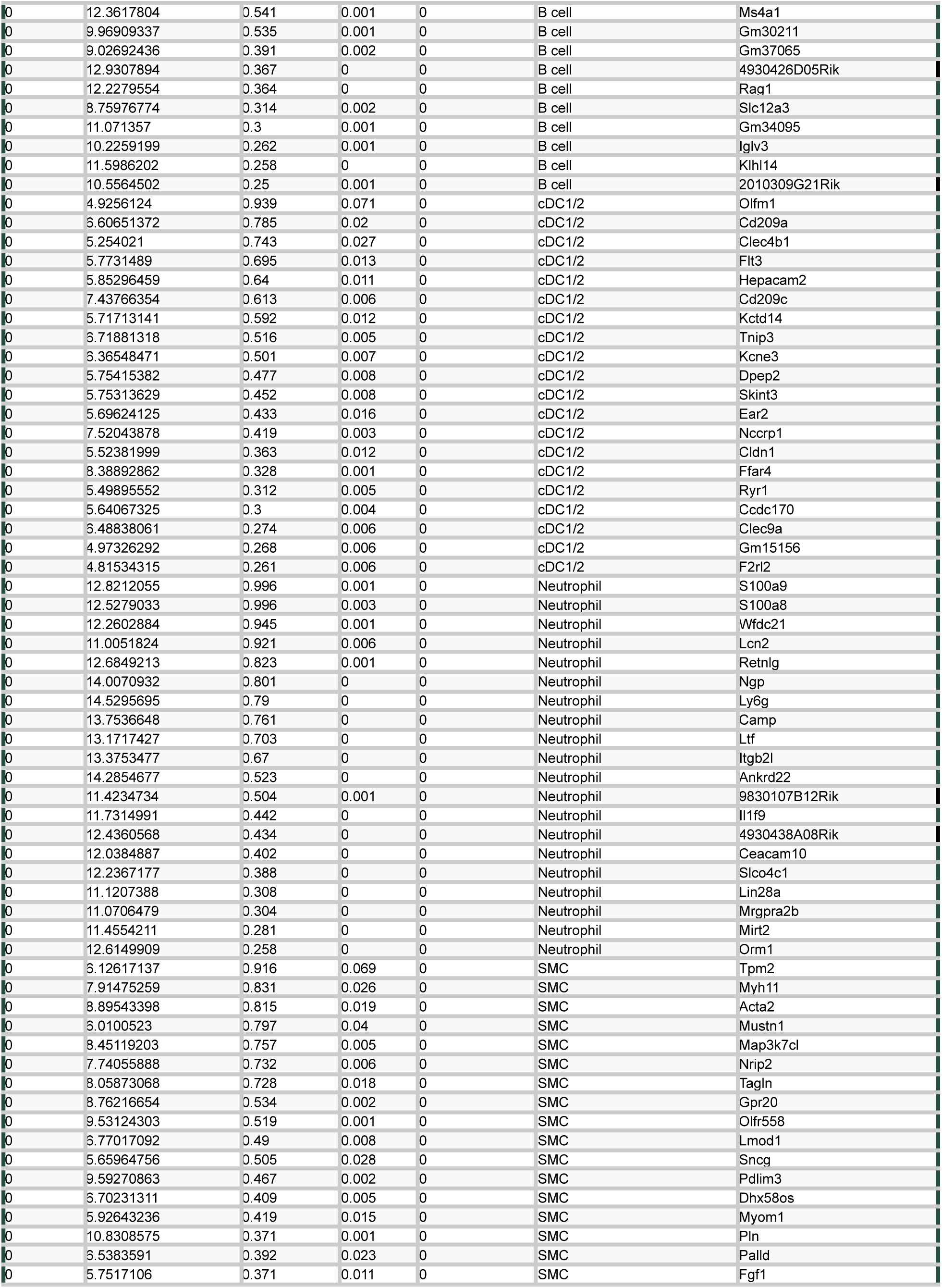
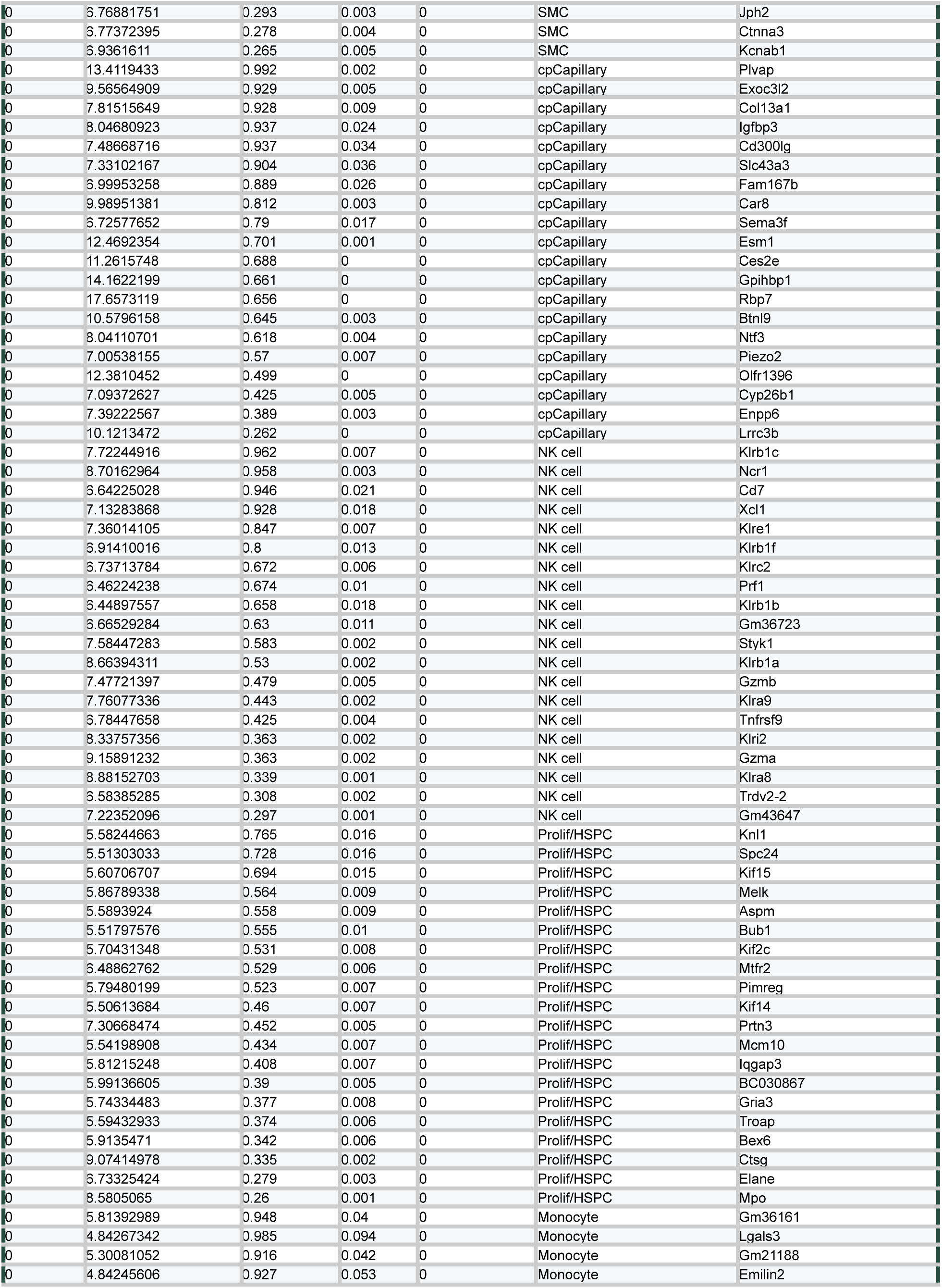
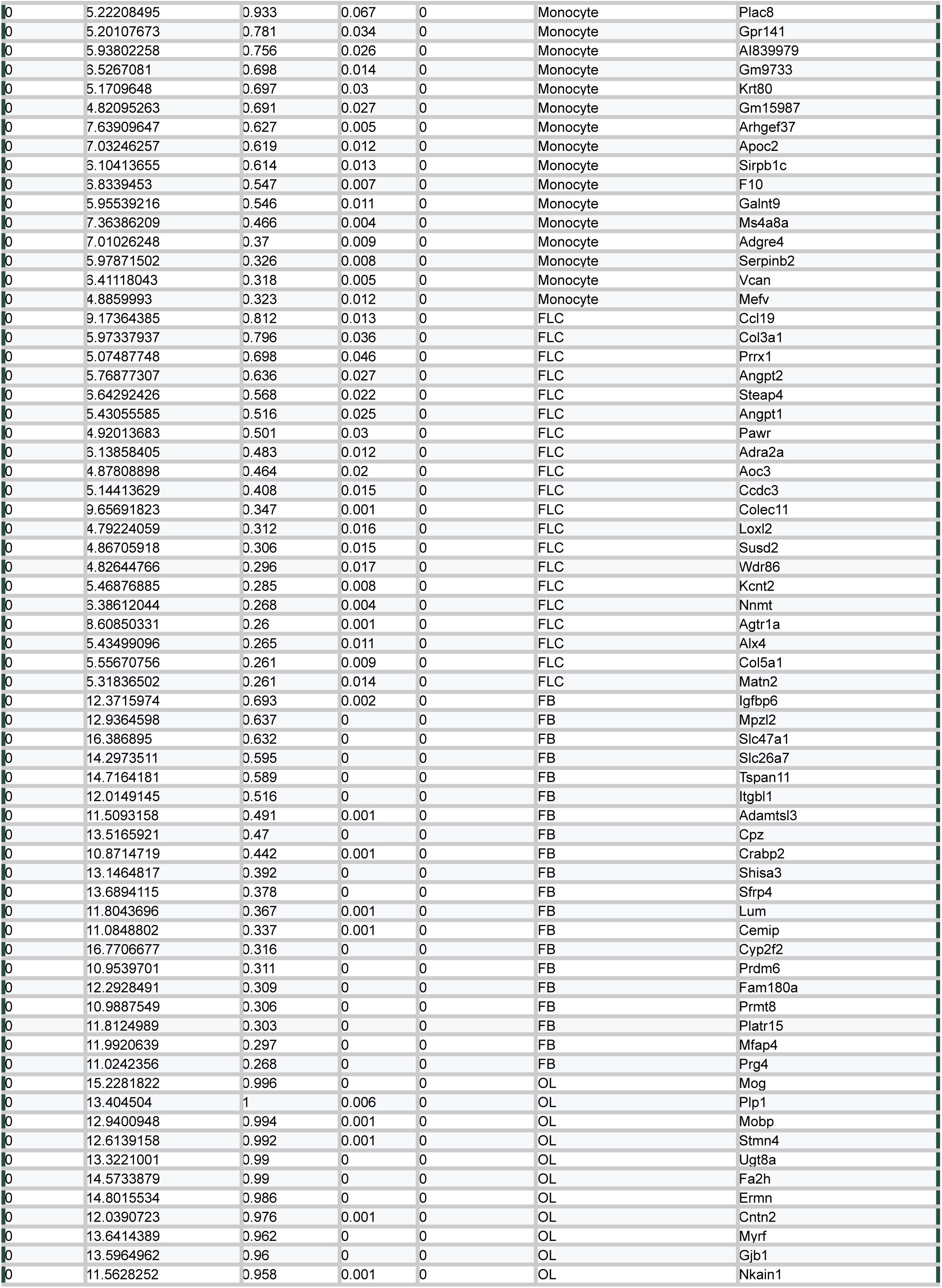
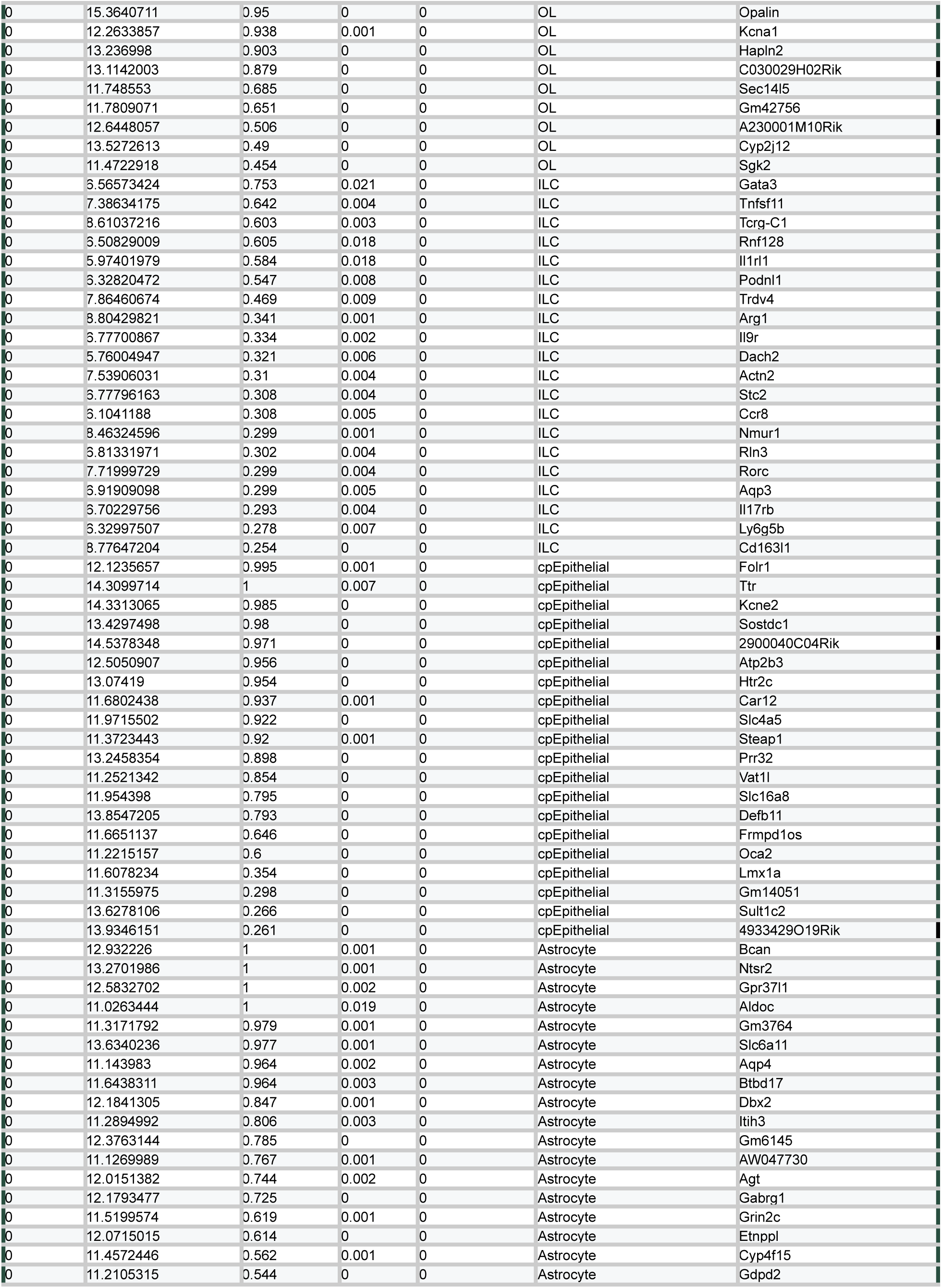
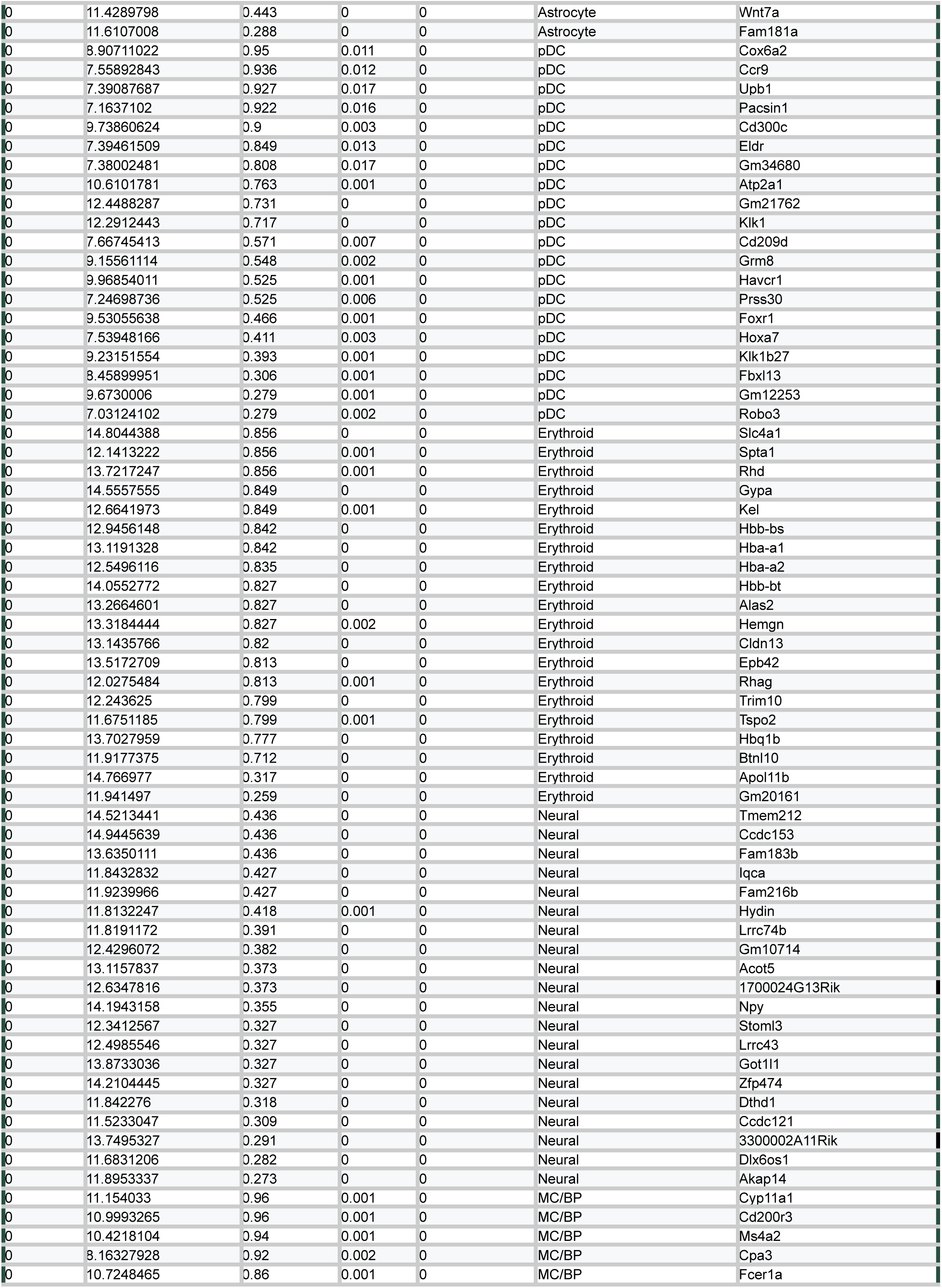
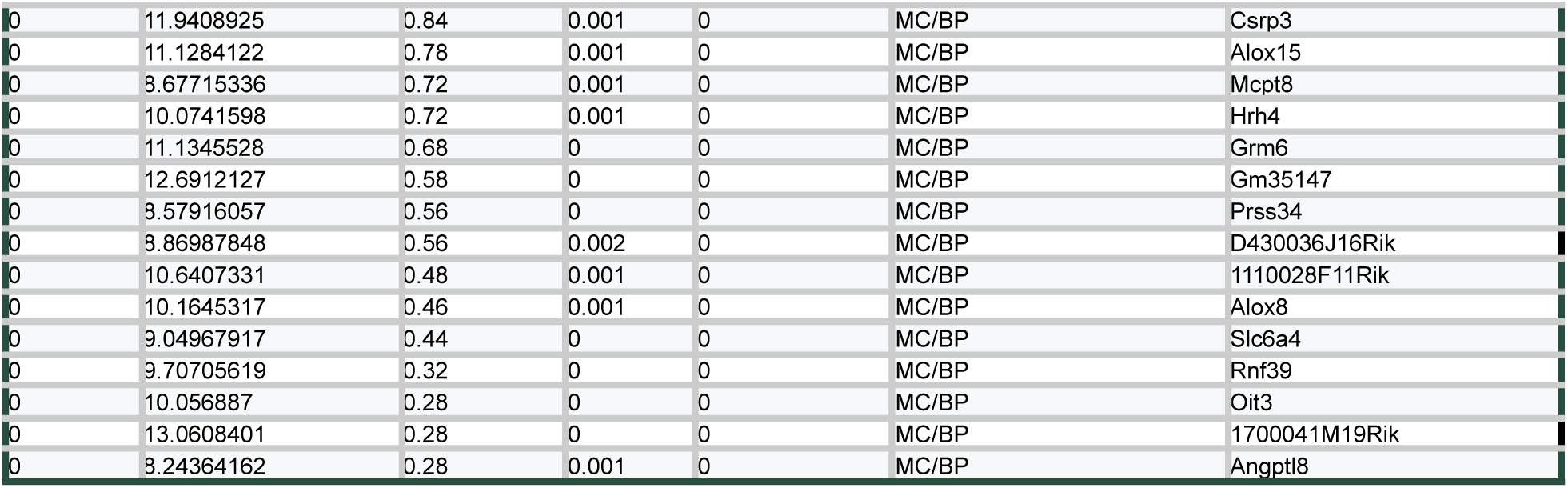
Cell type markers for scRNAseq atlas. Top 20 genes defining each broad cell type in the scRNA-seq atlas (**Figure 1B**). Fine annotation and resulting markers are located in **Figure S1**.

**Table S2.**
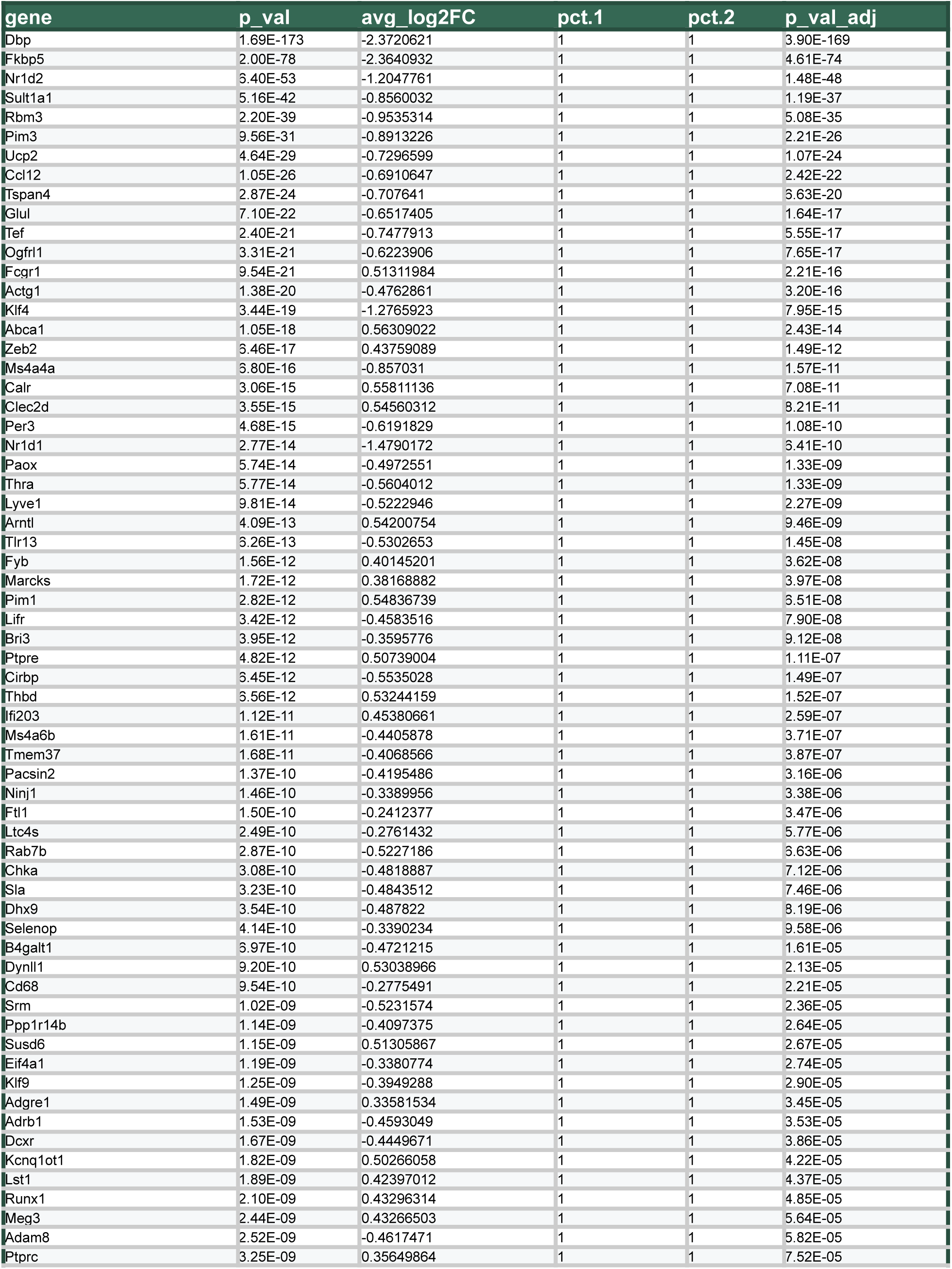
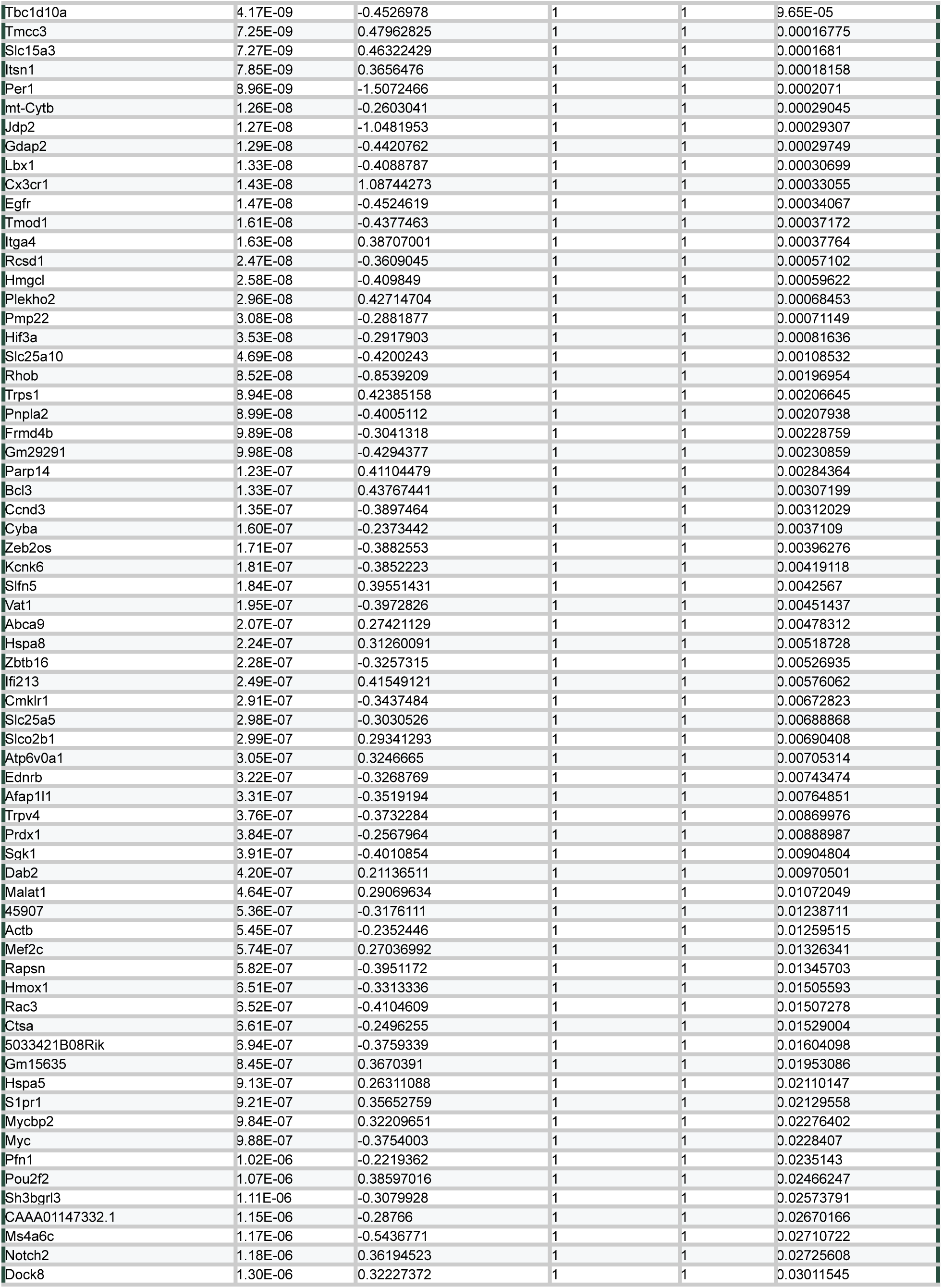
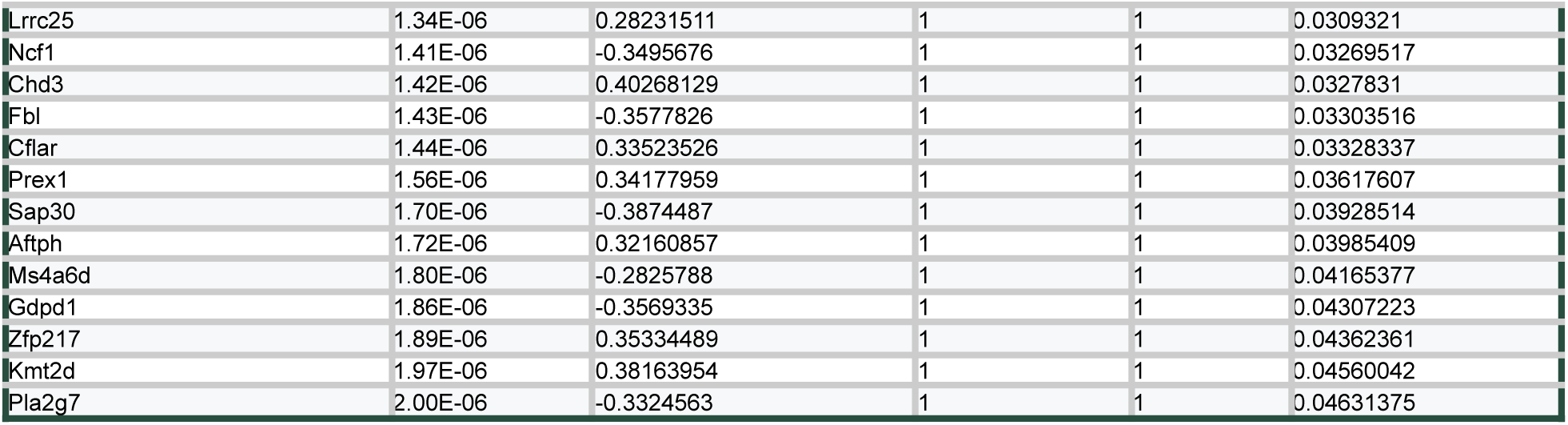
Pseudobulk comparison of BAMs from ZT0 and ZT12. Results of pseudobulk analysis comparing BAMs at ZT0 and ZT12, filtered for adjusted p-value <0.05. Negative values represent genes upregulated at ZT12 relative to ZT0.

**Table S3.**
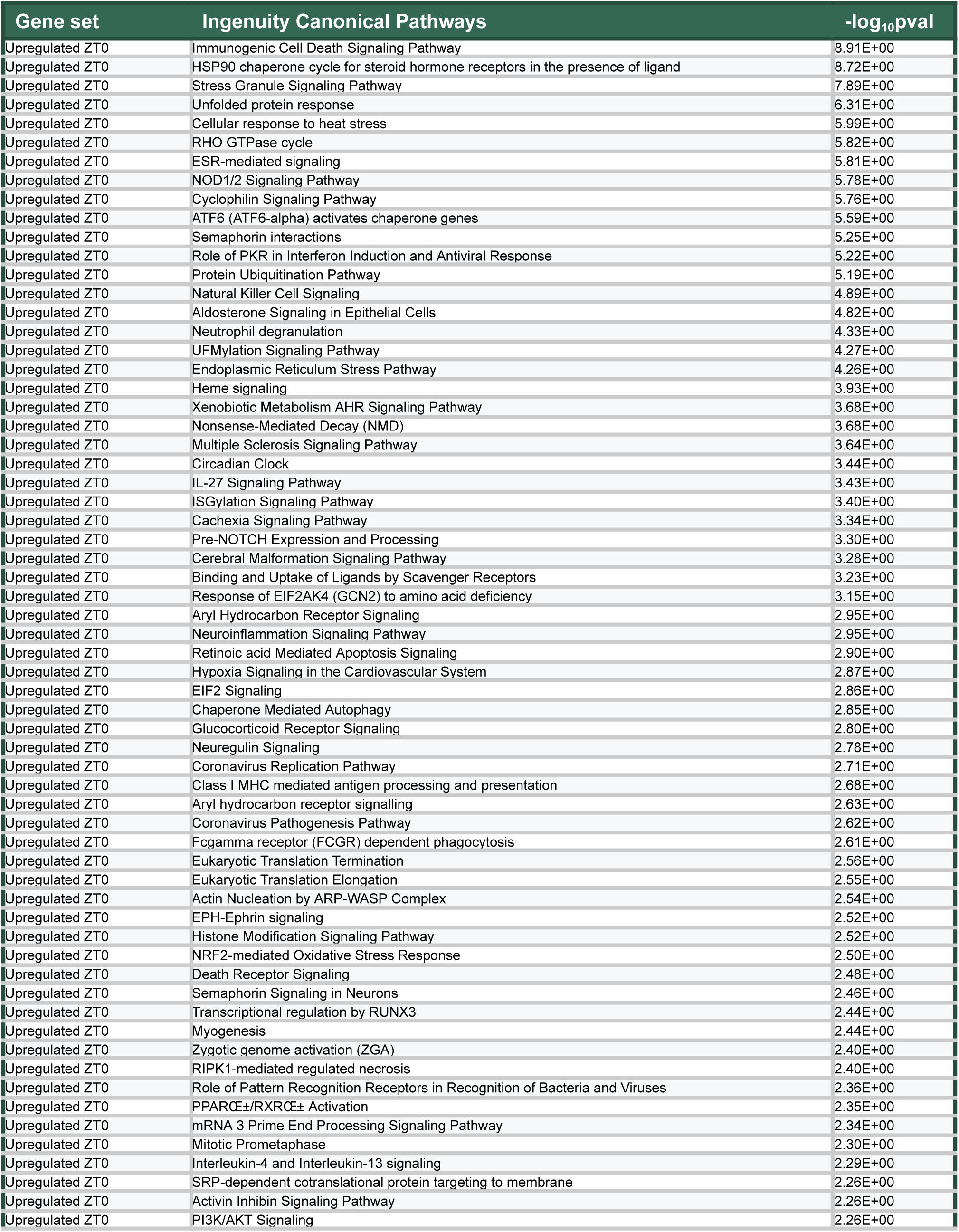
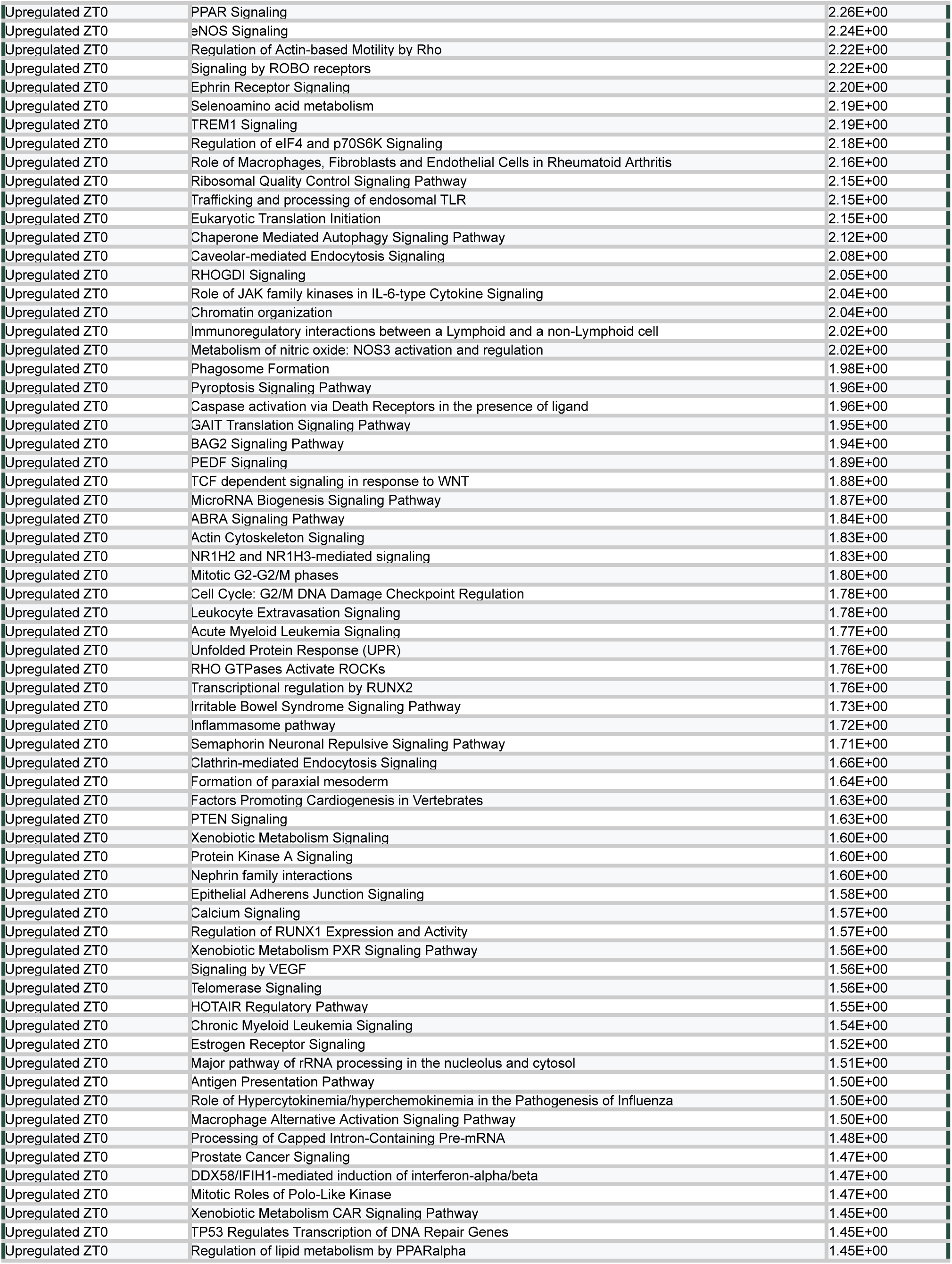
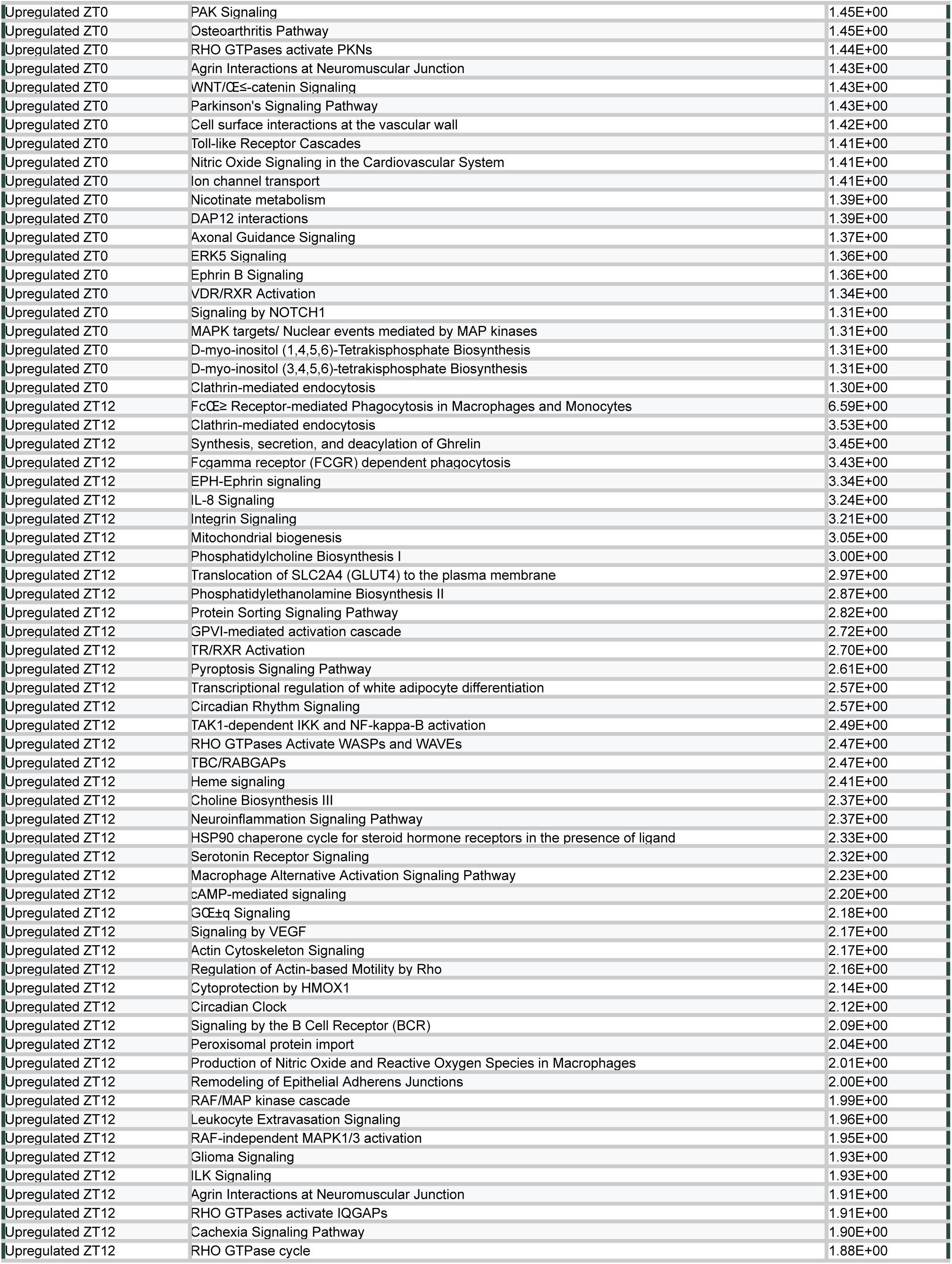
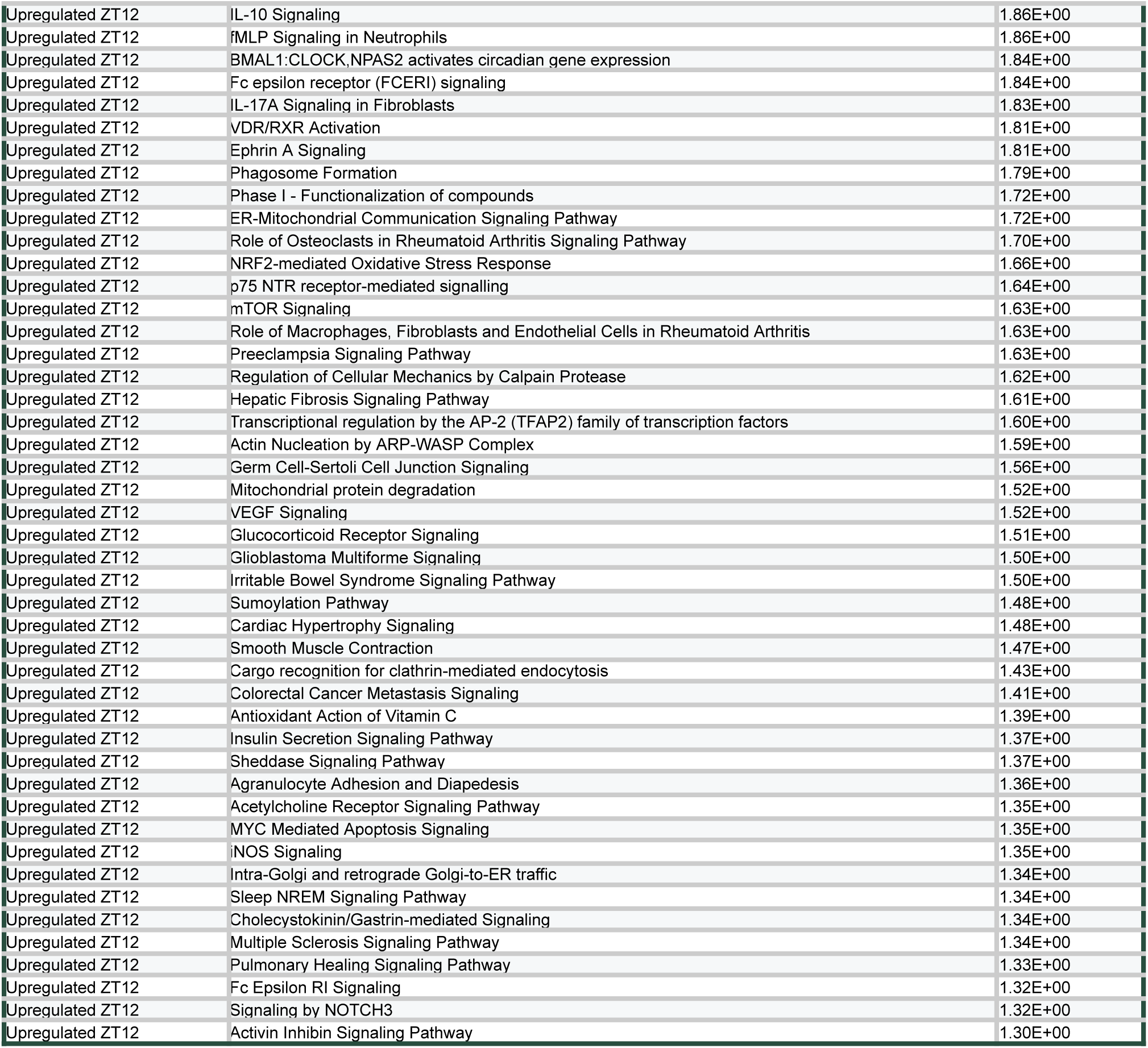
Ingenuity Pathway Analysis on genes upregulated in BAMs at ZT0 versus ZT12. All canonical pathways were identified by IPA analysis through the use of QIAGEN IPA (QIAGEN Inc.) on genes upregulated in BAMs at ZT0 versus ZT12. Input for IPA was DEGs identified by cell-level analysis. Output of IPA was filtered for -log_10_pvalue ≥ 1.3 and a minimum of 2 defining “molecules”.

**Table S4.**
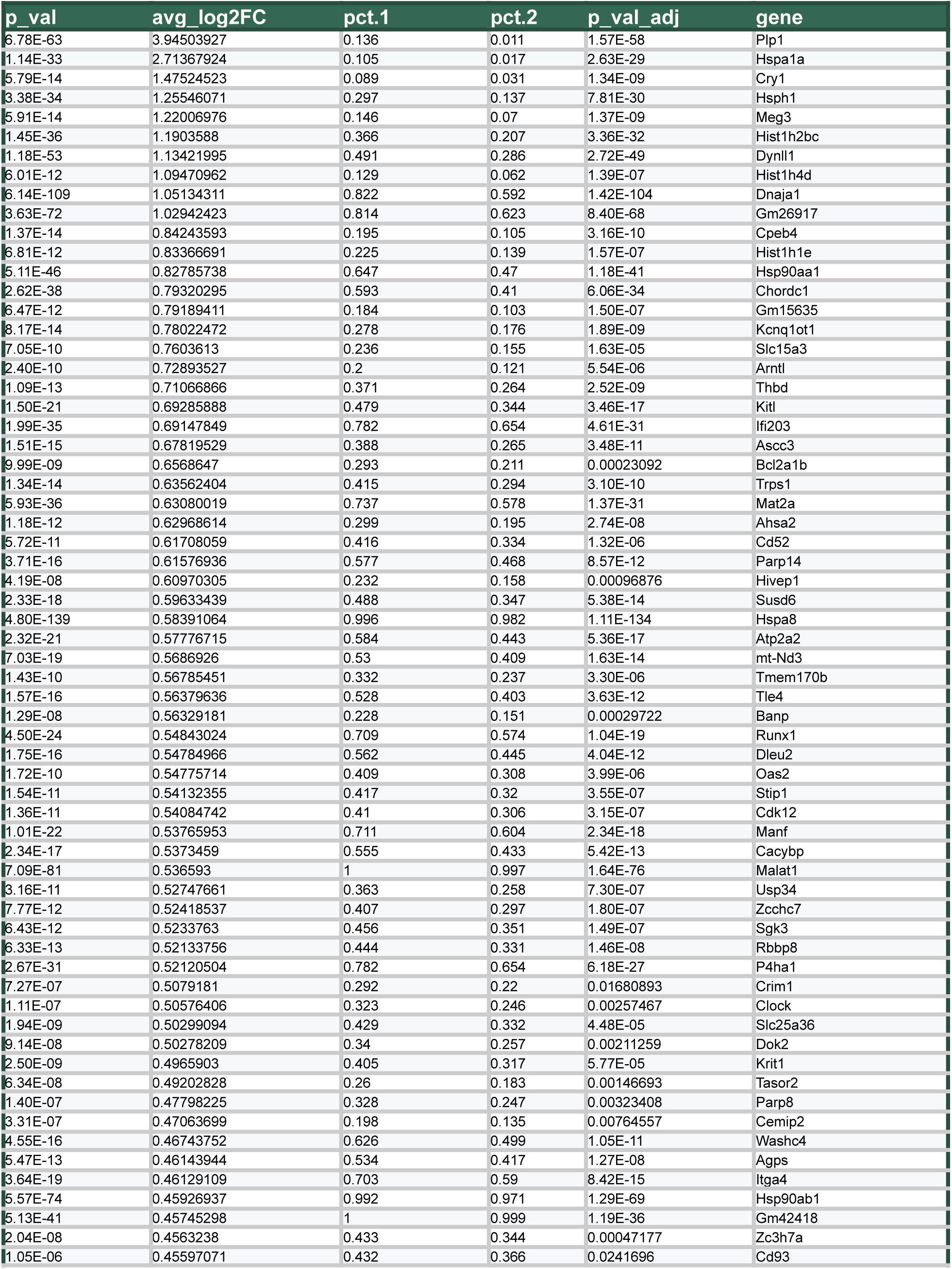
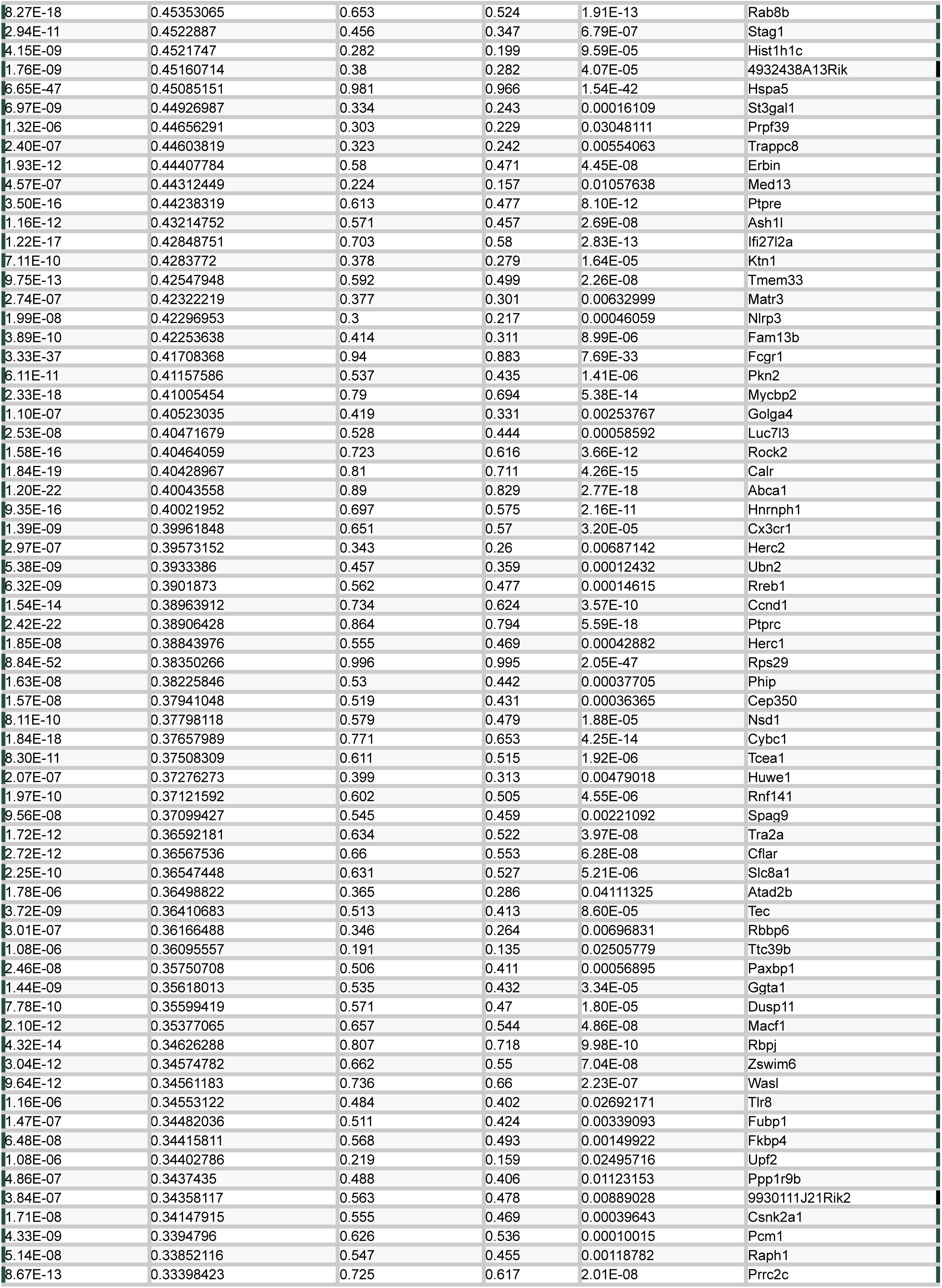
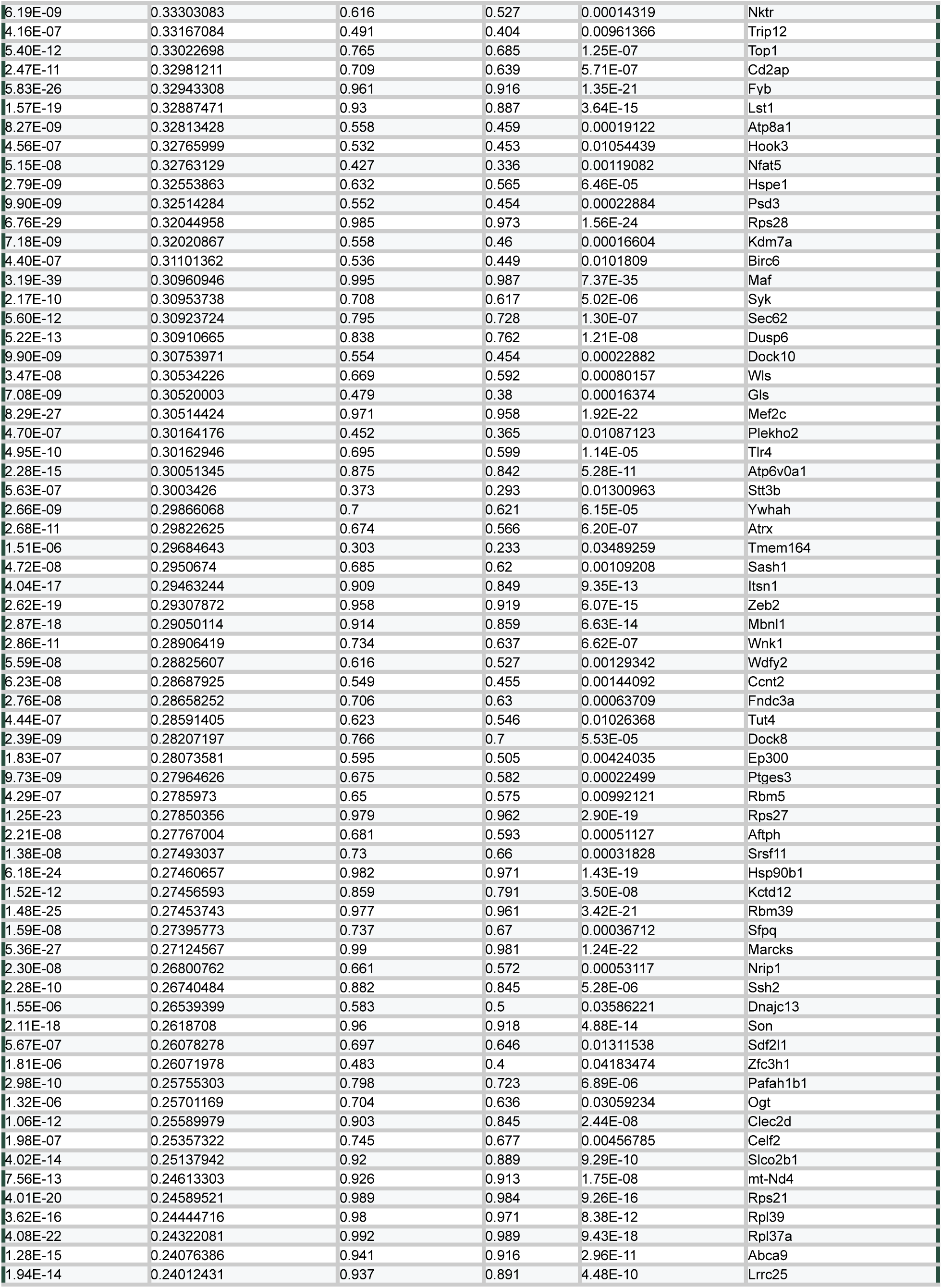
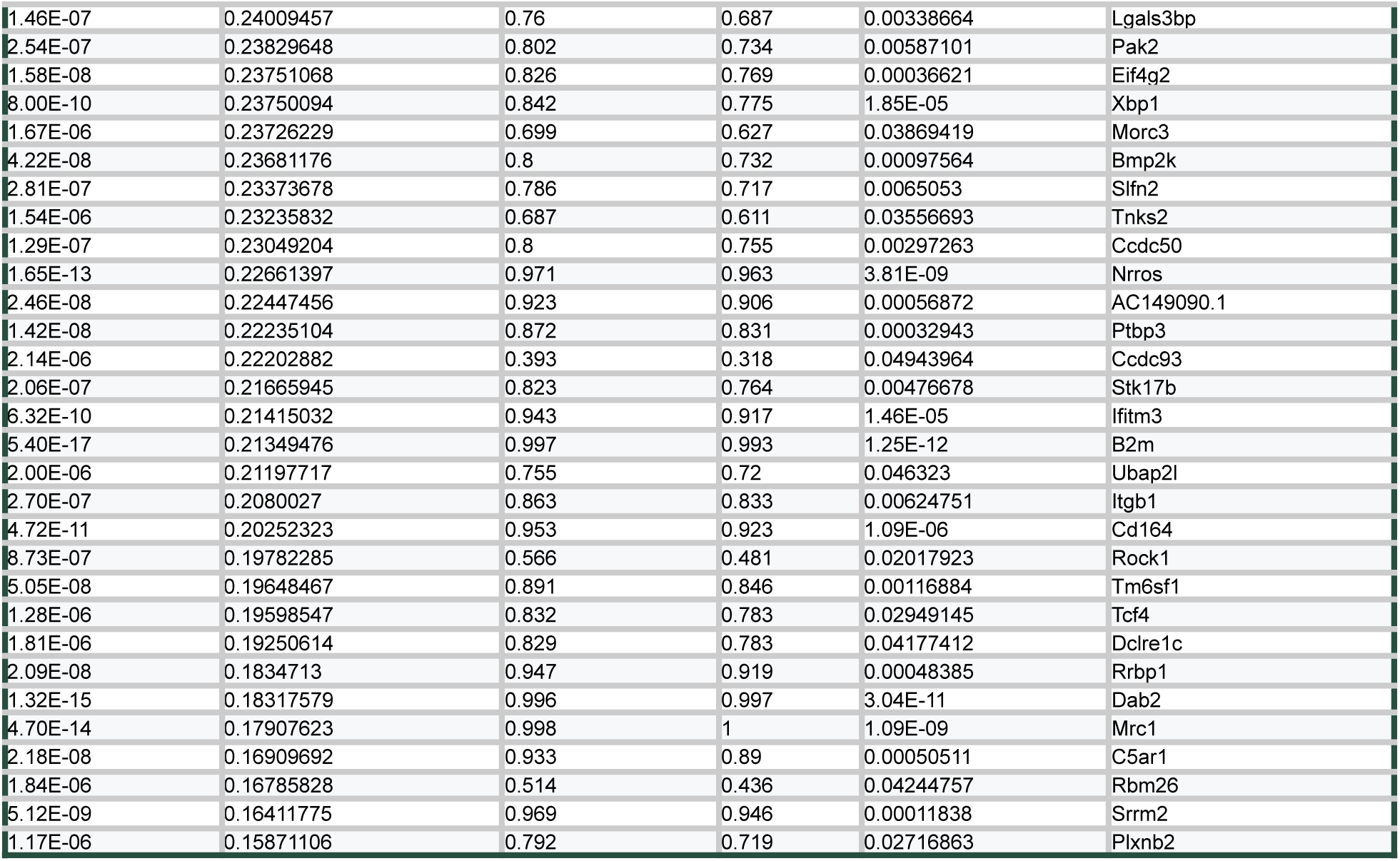
DEGs upregulated at ZT0 in BAMs from cell-level analysis. All genes upregulated at ZT0 from cell-level analysis of BAMs across all four time points. Gene list is filtered for adjusted p-value <0.05. Cell-level DEG analysis approach is described in **Methods**.

**Table S5.**
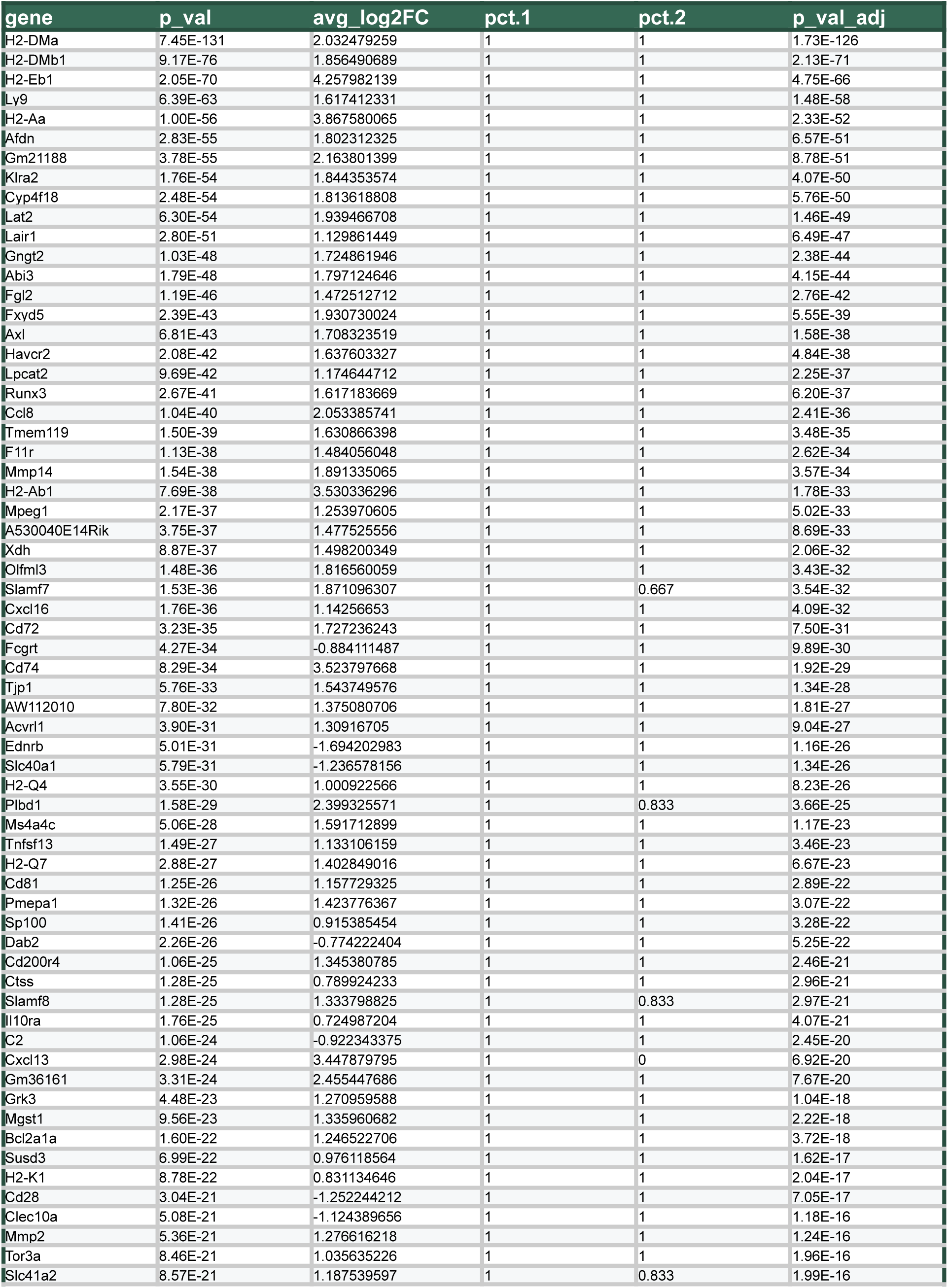
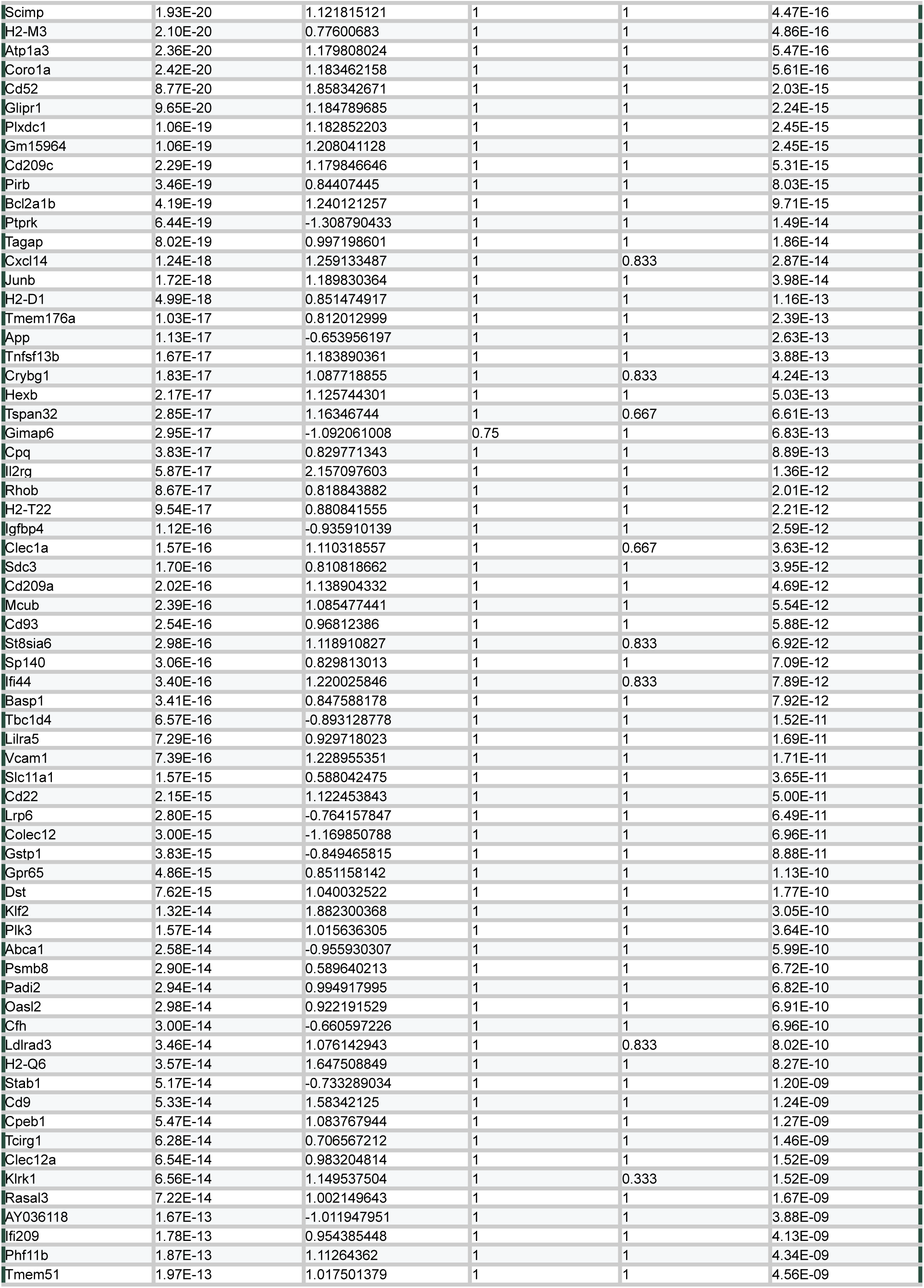
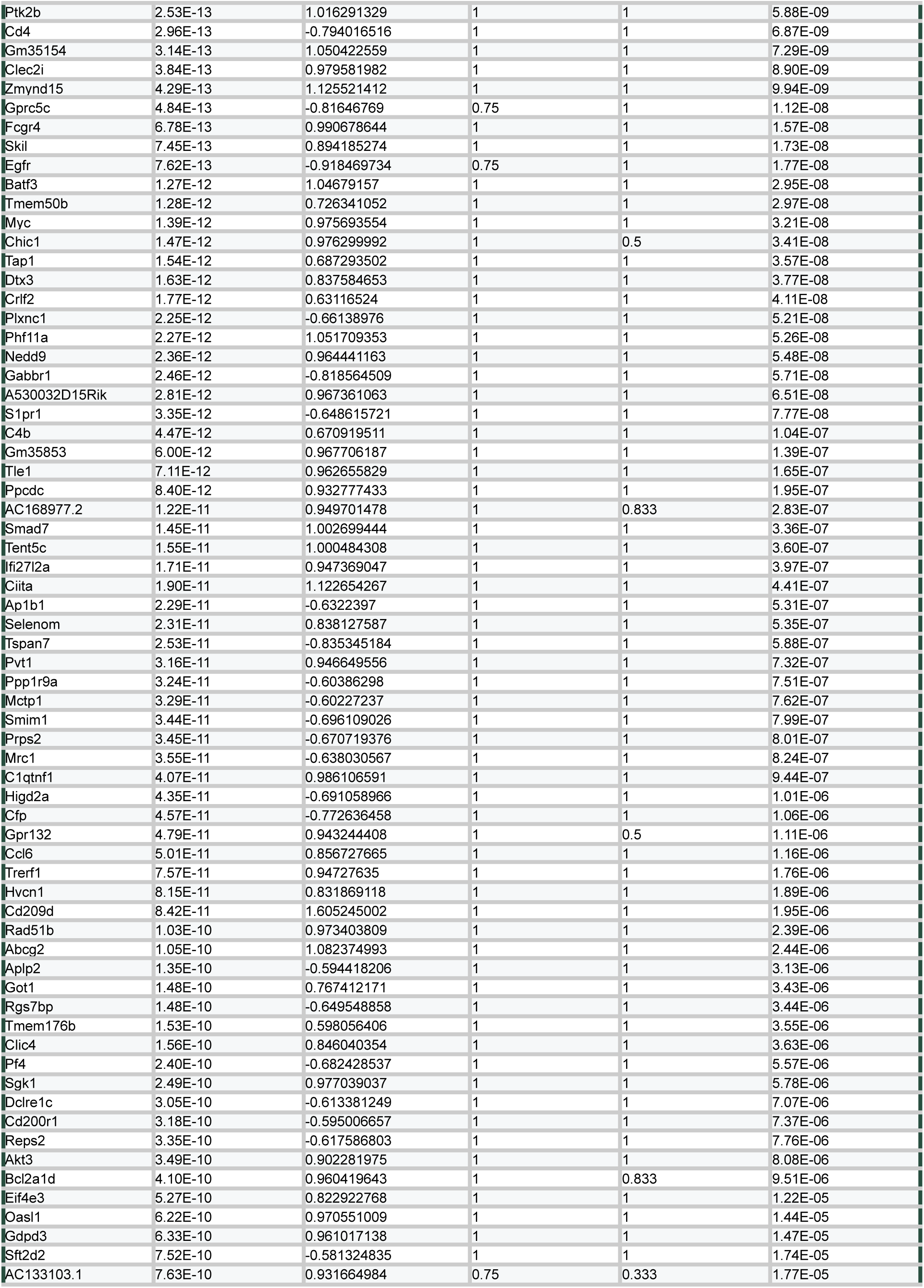
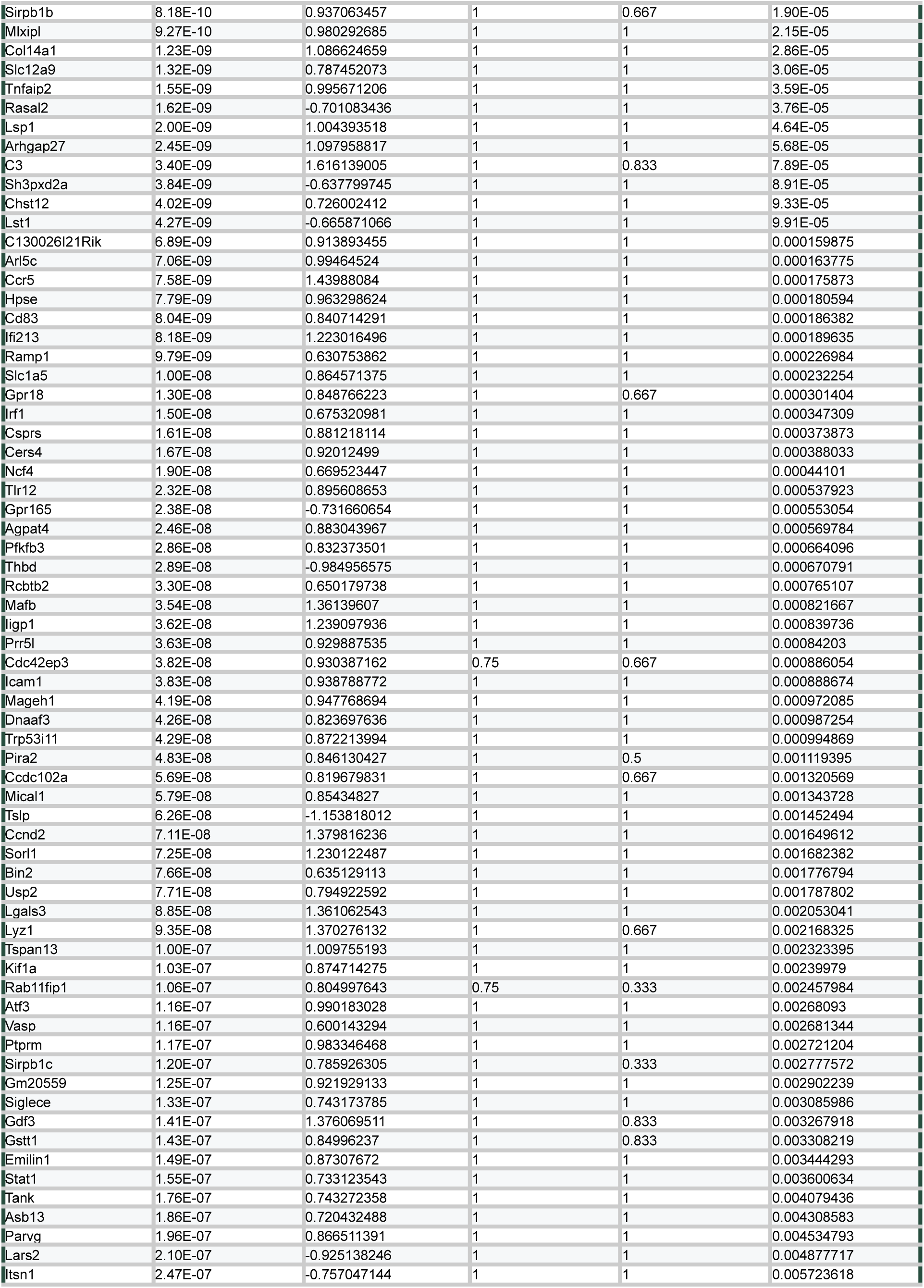
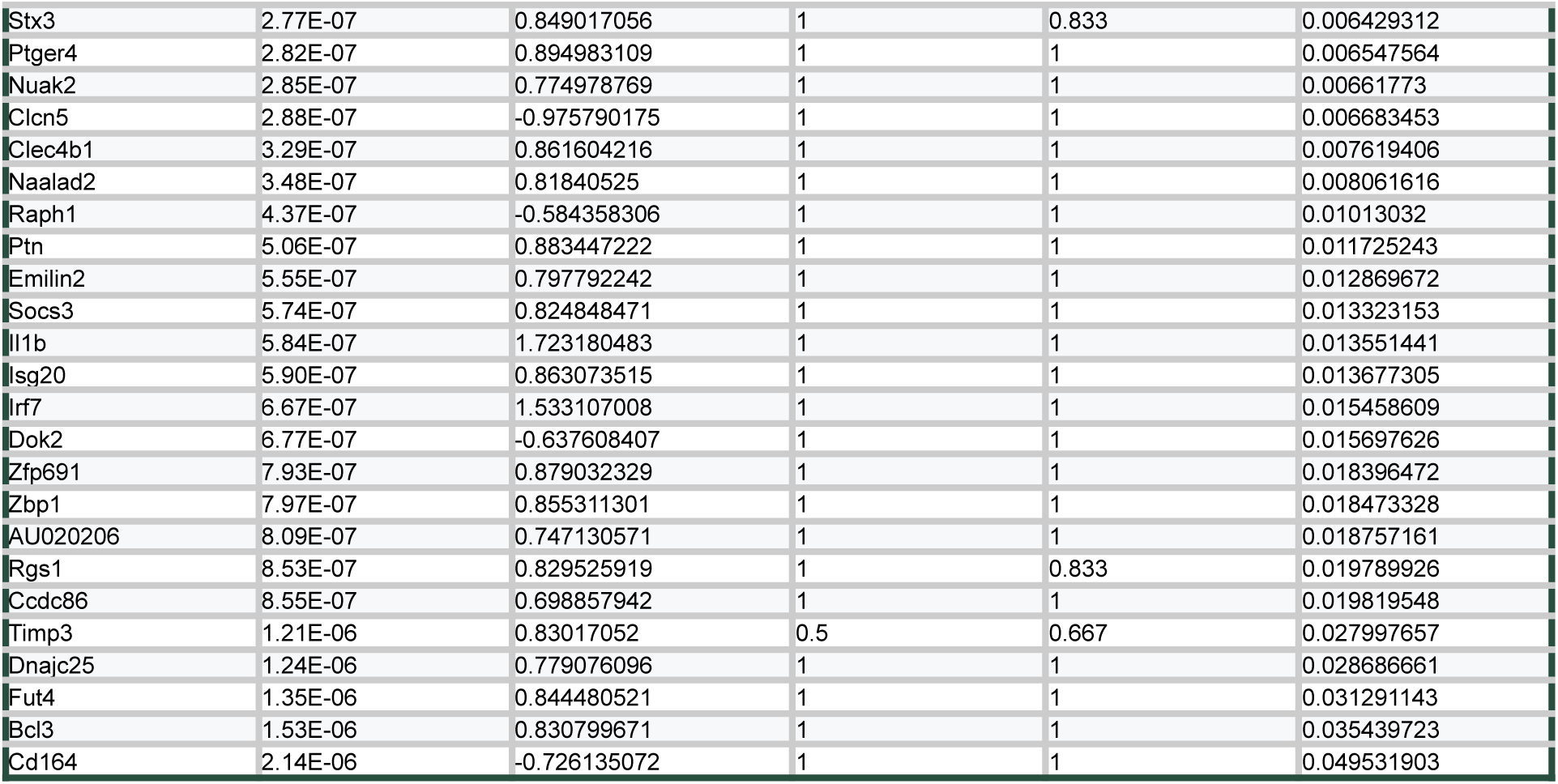
Adult versus aged BAM DEGs from pseudobulk comparison. Note: Positive fold changes represent upregulation in aged BAMs.

**Table S6.** Antibodies used for flow cytometry applications. Note: Panels and concentrations were adjusted for the fixed cell FEAST preparation.

| Antigen | Fluorophore | Clone | Manufacturer | Cat number | Dilution | Note |
| --- | --- | --- | --- | --- | --- | --- |
| CD11b | AF488 | M1/70 | BioLegend | 101217 | 200 |  |
| CD11b | BUV395 | M1/70 | BD Biosciences | 563553 | 200 |  |
| CD11b | BV510 | M1/70 | BioLegend | 101245 | 200 |  |
| CD11b | BV605 | M1/70 | BioLegend | 101237 | 200 |  |
| CD11c | PE-Cy5.5 | N418 | Thermo Fisher Scientific | 35-0114-82 | 200 |  |
| CD206 | BV421 | C068C2 | BioLegend | 141717 | 200 |  |
| CD206 | PE | C068C2 | BioLegend | 141720 | 100 |  |
| CD38 | PE | 90 | BioLegend | 102708 | 200 |  |
| CD38 | PE-Cy7 | 90 | BioLegend | 102718 | 200 |  |
| CD38 | BV421 | 90 | BioLegend | 102732 | 200 |  |
| CD38 | BUV395 | 90/CD38 | BD Biosciences | 740245 | 200 |  |
| CD38 | BUV805 | 90/CD38 | BD Biosciences | 741955 | 200 |  |
| CD45 | PE-Cy5.5 | 30-F11 | Thermo Fisher Scientific | 35-0451-82 | 500 | Used 1:200 for FEAST |
| CD45 | PE | 30-F11 | BioLegend | 103106 | 500 |  |
| CD45 | BUV805 | 30-F11 | BD Biosciences | 568336 | 200 |  |
| CD64 | PE-Cy7 | X54-5/7.1 | BioLegend | 139313 | 100 |  |
| CD64 | BV421 | X54-5/7.1 | BioLegend | 139309 | 100 |  |
| CD68 | APC-Cy7 | FA-11 | BioLegend | 137024 | 400 |  |
| CX3CR1 | BV711 | SA011F11 | BioLegend | 149031 | 800 |  |
| CX3CR1 | AF488 | SA011F11 | BioLegend | 149022 | 800 |  |
| CX3CR1 | AF647 | SA011F11 | BioLegend | 149004 | 800 |  |
| GR-1 | BUV805 | RB6-8C5 | BD Biosciences | 741920 | 100 |  |
| GR-1 | AF488 | RB6-8C5 | BioLegend | 108417 | 100 |  |
| GR-1 | PE | RB6-8C5 | BioLegend | 108408 | 100 |  |
| Ly-6C | BV510 | HK1.4 | BioLegend | 128033 | 200 |  |
| MHC-II | BV785 | M5/114.15.2 | BioLegend | 107645 | 200 |  |
| β-Amyloid, 1-16 | AF647 | 6E10 | BioLegend | 803021 | 200 |  |

